# The small-bat-in-summer paradigm: energetics and adaptive behavioural routines of bats investigated through a stochastic dynamic model

**DOI:** 10.1101/2022.09.28.509895

**Authors:** Mari Aas Fjelldal, Amandine Sophie Muller, Irja Ida Ratikainen, Clare Stawski, Jonathan Wright

## Abstract

Strong seasonality at high latitudes represents a major challenge for many endotherms as they must balance survival and reproduction in an environment that varies widely in food availability and temperature. Being heterotherms, bats spend long cold winters in hibernation, avoiding the challenges faced by many animals. To avoid energetic mismatches caused by limited foraging time and stochastic weather conditions, bats can also employ this energy-saving state of torpor during summer to save accumulated energy reserves. However, at high latitudes small-bats-in-summer face a particular challenge: as nocturnal foragers they rely on the darkness of the night to avoid predators and/or interspecific competition, but for many the summer involves short nights of mostly twilight, and even a lack of true night at the northernmost distributions of some bat species. To investigate optimal individual behaviour across diurnal cycles, we constructed a stochastic dynamic model of bats living at high latitudes. Using a detailed parameterized model framework with values that are representative for our study system, we show that individual energetic reserves are a strong driver of day-time use of torpor and night-time foraging behaviour alike, with these linked effects being both temperature and photoperiod dependent. We further used the model framework to predict survival probabilities at five locations across a latitudinal gradient (60.1°N to 70.9°N), finding that photoperiod is the main limiting factor to bat species distributions. To verify the accuracy of our model results, we compared predictions for optimal decisions with our own empirical data collected on northern bats (*Eptesicus nilssonii*) from two latitudes in Norway. The similarities between our predictions and observations provide strong confirmation that this model framework incorporates the most important drivers of diurnal decision-making in bat physiology and behaviour. Our model findings regarding state-dependent decisions in bats should therefore contribute to the understanding of how bats cope with the summer challenges at high latitudes.

## Introduction

The dilemma of small-birds-in-winter needing to balance high energetic demands with limited foraging opportunities during short winter days is a well-known paradigm in behavioural ecology research (e.g. Bednekoff & Houston 1994; McNamara *et al*. 1994; Brodin 2007; Brodin *et al*. 2017). A similar but less well known dilemma is the one small bats face in summer. At northern latitudes the summer seasons are short, with only a few months available for the bats to reproduce and build up sufficient fat reserves to survive the subsequent long and cold winter in hibernation. The northern summer season is characterised by short nights consisting largely of periods of twilight, which limit the time nocturnal creatures like bats have available to forage in safety from aerial predators (Speakman *et al*. 2000). Furthermore, most temperate zone bat species are insectivorous, and therefore dependent upon a food source that constantly fluctuates with environmental conditions, such as temperature and precipitation (see Taylor 1963; Speakman *et al*. 2000). Being the only mammals in the world capable of powered flight, bats also have a high energy expenditure during each foraging trip (Winter & Von Helversen 1998). Consequently, when summer nights are short and weather conditions are poor, bats may face a mismatch between the energy expenditure needed for flight and the amount of available prey when foraging, which potentially limits the northern range of many bat species (Parker *et al*. 1997).

One way for insectivorous bats to overcome these energetic challenges of summer foraging is to use the energy-saving state of torpor during day-time or on nights when foraging prospects are poor (Ruf & Geiser 2015). Torpor is characterised by a controlled and reversible reduction in heart rate and oxygen consumption while temporarily abandoning the defence of a high and stable body temperature (Geiser 2021). Due to its inherent potential for saving large amounts of energy, temporal heterothermy is an important evolutionary trait that in the short-term may enhance individual survival and in the long-term reduce the risk of species extinctions (Geiser & Turbill 2009; Liow *et al*. 2009; Stawski *et al*. 2014). However, despite the apparent benefits of torpor (and hibernation) as an energy-saving state during inclement conditions, there are some inherent costs that will influence any adaptive state- and weather-dependent use of torpor. Firstly, employing torpor may help save energy, but rewarming from torpor brings associated energetic and physiological costs that may limit the energetic benefit (Currie *et al*. 2015; Landes *et al*. 2020). Detrimental physiological costs of prolonged hibernation periods have also been identified (Humphries *et al*. 2003; Boyles *et al*. 2020), but these were found to be reversible following frequent arousals (Humphries *et al*. 2003). Such costs should therefore be less likely to affect the expression of daily torpor in non-reproductive bats outside of the hibernation season. Secondly, and perhaps more importantly for the individual bat during summer, are the potential benefits of being awake. Most bat species are highly social and dependent upon frequent interactions with roost mates to obtain important benefits like social grooming, cooperative offspring care, sharing of information or attracting mates (Wilkinson *et al*. 2016; Chaverri *et al*. 2018). Bats strategically choose warm roosts during summer to facilitate lower thermoregulatory costs of being awake, which is important particularly for pregnant or lactating females as they face added costs to their reproduction if they employ frequent deep torpor bouts (Speakman & Rowland 1999; Lausen & Barclay 2003; Lourenço & Palmeirim 2004). Non-reproductive and post-lactating individuals may also spend considerable periods of time awake during day- and night-time in summer, although the expression of torpor is found to vary greatly with weather and roost type (Lausen & Barclay 2003; Bergeson *et al*. 2021; Fjelldal *et al*. 2021). Even in non-reproductive bats, there is likely to be a complex relationship between the optimal energetic decision to use torpor versus being awake in the roost (henceforth referred to as *resting*) or foraging outside. Environmental temperature has been identified as a strong driver of general bat behaviour (e.g. see Ruf & Geiser 2015), but body mass have also been found to impact individual strategies such as summer night-time torpor use (Fjelldal *et al*. 2021) and microclimate selection during winter hibernation (Boyles *et al*. 2007). Indeed, the effect of within-individual variation in body mass (i.e. fat reserves and stomach contents) on daily decision making in bats is largely unstudied due to the challenges of continuous monitoring of free-ranging individuals, making this an appropriate moment to fully explore our theoretical expectations.

This study therefore presents one of the first theoretical models for understanding individual optimal state-dependent decisions in daily heterotherms like bats. Our stochastic dynamic optimization model is developed using detailed parameterisation from natural systems in order to quantitatively predict optimal behaviour as accurately as possible. By incorporating quantified values such as dynamic photoperiods, light- and state-dependent predation risk, representative daily temperature conditions for the roost and outside air, temperature-dependent prey availabilities and physiological costs of different activities, our model includes all of the variables we consider likely to be important drivers of behaviour and thermal physiology in non-reproductive bats. The model organism here is based upon physiological data collected from the literature on various species of insectivorous small-bodied bats (see Supplementary Materials 1). Physiological energetics in bats depend upon body mass and temperature, but show no apparent latitudinal or climate zone effects of local adaptation (Speakman & Thomas 2003; Skåra *et al*. 2021; Fjelldal *et al*. 2022). Our model organism should therefore be representative of any small-bodied insectivorous bat species inhabiting seasonal environments. To test the applicability of our model predictions, we briefly compare the model results with empirical data collected on northern bats (*Eptesicus nilssonii*) at two latitudes in Norway (Nittedal, 60.1°N and Trondheim, 63.4°N). Our study aims are three-fold. Firstly, we seek to discover how bats at the northern limits of their species distributional range overcome the associated challenges of summer survival through the strategic use of torpor. Secondly, we want to explore the magnitude of state-dependency on this behavioural and thermal physiological decision-making employed across a range of climatic conditions. Finally, by running model simulations of one month summer survival across a large latitudinal gradient, we attempt to determine the main limiting causes of bat species distributional ranges at their northern limits. Through our findings, we aim to provide a deeper understanding of the diurnal nature of strategic decision-making by bats, and thus their scope for adaptation to changing environments.

## Methods

### Stochastic dynamic programming

We used stochastic dynamic programming (see Houston *et al*. 1988; Clark & Mangel 2000) to investigate the optimal sequence of state-dependent decisions made by small bats in summer. The model time horizon (*T*) consisted of 72 daily timesteps (*t*) repeated for 30 days (*D*), thus focusing on the life of small bats through the lightest summer month of the year. As we wanted to investigate the impact of behavioural and physiological decisions on individual energetics for non-breeding bats in summer, we allowed the three following behaviours for the bats to choose from:

1. Torpor – bat in the roost, torpid
2. Resting – bat in the roost, awake
3. Foraging – bat outside of the roost, foraging

#### Parametrization

The parametrization was developed by collecting biologically relevant values from the literature for the physiological parameters (see Supplementary Materials 1). For the different environmental scenarios, we used temperature data obtained through the Norwegian Centre for Climate Services and our own light-measurements at two high-latitude field sites: Nittedal, Norway (60.1°N, 10.8°E) and Trondheim, Norway (63.4°N, 10.4°E). The light-measurements were used to estimate a light-dependent predation threat variable and a light-dependent energetic competition cost (i.e. we assume variation in fitness costs like predation threat, interspecific competition for food and mobbing by non-competitor species to be determined by light conditions and energy reserves, see below). Baseline parameter values for our stochastic dynamic models are shown in Table 1a and Table 1b, comparing two locations that differ in latitude and therefore in light-conditions and the associated predation threats and competition costs. The values represent average conditions for the month of July in Nittedal and Trondheim, thus representing one of the months with the Shortest nights and the lightest twilight at these high-latitude locations.

**Table 1a:** General parameter definitions and model baseline values. Details and justifications for the parameterization can be found in Supplementary Materials 1, including equations for the air temperature cycles across day types. Text in bold indicate when values differ between locations.

| Symbol | Parameter | Values | Main reference |
| --- | --- | --- | --- |
| D | Number of days in month | 30 | Model structure |
| T | Number of timesteps in month | 2160 (72 × 30) | Model structure |
| X<sub>0</sub> | Mass of bat with no fat reserves | 7 g | Model assumption |
| X<sub>max</sub> | Max body fat deposit | 3 g (40 discrete steps) | Model assumption |
| X<sub>min</sub> | Fat reserve threshold for survival | 0.05 g | Model assumption |
| X<sub>dec</sub> | Threshold for decreasing survival | 2 g | Model assumption |
| Y<sub>state</sub> | Temperature state | *Torpid* or *Non-torpid* | Model structure |
| tnz | Thermal neutral zone | ≥ 29°C | Geiser and Brigham (2000) |
| w | Weather type across 24 hours | Very warm, dynamic warm, stable warm, dynamic cold, stable cold, very cold | Model structure |
| air<sub>very_warm</sub>(*t*) | Temperature outside, very warm day | Equation S1.5 | Norwegian Centre for Climate Services (NCCS) |
| air<sub>dynamic_warm</sub>(*t*) | Temperature outside, dynamic warm day | Equation S1.6 | NCCS |
| air<sub>stable_warm</sub>(*t*) | Temperature outside, stable warm day | Equation S1.7 | NCCS |
| air<sub>dynamic_cold</sub>(*t*) | Temperature outside, dynamic cold day | Equation S1.8 | NCCS |
| air<sub>stable_cold</sub>(*t*) | Temperature outside, stable cold day | Equation S1.9 | NCCS |
| air<sub>very_cold</sub>(*t*) | Temperature outside, very cold day | Equation S1.10 | NCCS |
| roost<sub>warm</sub>(air(*w*, *t*), *t*) | Temperature roost, warm day types | air(*w*, *t*) + (2.34 + 0.27 × *t* - 2.75 × 10<sup>-4</sup> × *t*<sup>2</sup> - 4.88 × 10<sup>-5</sup> × *t*<sup>3</sup>) | Own data |
| roost<sub>cold</sub>(air(*w*, *t*)) | Temperature roost, cold day types | air(*w*, *t*) + 2°C | Own data |
| p<sub>w</sub> | Probability of very warm days, dynamic warm days, stable warm days, dynamic cold days, stable cold days, and very cold days, respectively | **For Nittedal:** 0.24, 0.48, 0.07, 0.20, 0.01, 0<br>**For Trondheim:** 0.13, 0.30, 0.04, 0.47, 0.04, 0.02 | NCCS |

**Table 1b:** Parameter definitions and model baseline values for each behaviour. Equations for the temporal predation threats and energetic competition costs can be found in Supplementary Materials 1.

| Symbol | Parameter | Values | Main reference |
| --- | --- | --- | --- |
| *Behaviour 1 (torpor)* |  |  |  |
| $\lambda(\text{air}(w,t))$ | Food availability | 0 | Model assumption |
| $\mu(t)$ | Predation threat baseline | 0 | Model assumption |
| $C_{\text{TMR}}(\text{roost}(w,t))$ | Hourly cost of TMR | $0.0008 \times \exp(0.086 \times \text{roost}(w,t) \times X_0)$ if $C_{\text{TMR}} < C_{\text{BMR}} \times 0.9$;<br>$C_{\text{BMR}} \times 0.9$ if $C_{\text{TMR}} \geq C_{\text{BMR}} \times 0.9$ | Fjelldal *et al.* (2022)* |
| $C_{\text{RW}}(\text{roost}(w,t))$ | Rewarming cost | $0.018 - 0.00046 \times \text{roost}(w,t) \times X_0$ | Turbill (2008)* |
| *Behaviour 2 (resting)* |  |  |  |
| $\lambda(\text{air}(w,t))$ | Food availability | 0 | Model assumption |
| $\mu(t)$ | Predation threat baseline | 0 | Model assumption |
| $C_{\text{BMR}}$ | Hourly BMR cost (for roost $\geq \text{tnz}$) | $0.0052 \text{ g} \times X_0$ | Geiser and Brigham (2000)* |
| $C_{\text{RMR}}(\text{roost}(w,t))$ | Hourly RMR cost (for roost $< \text{tnz}$) | $0.036 \text{ g} \times X_0 - 0.0011 \times \text{roost}(w,t) \times X_0$ | Geiser and Brigham (2000)* |
| $\theta$ | Resting fitness benefit | 0.0015 | Model assumption |
| *Behaviour 3 (foraging)* |  |  |  |
| $\alpha$ | Potential energy gain per hour | 1.3 g | Sørås *et al.* (2022)* |
| $\lambda(\text{air}(w,t))$ | Food availability | $\alpha / (1 + \exp(-0.52 \times (\text{air}(w,t) - 8)))$ | Speakman *et al.* (2000)* |
| $\kappa$ | Coefficient for energy reserve effect on foraging success | -0.03 | Model assumption |
| $p_{\text{food}}(x)$ | Probability of finding available food | $0.9 \times \exp(\kappa(x))$ | Model assumption |
| $\mu(t)$ | Predation threat baseline | 0.0001 if $\mu < 0.0001$ & 0.2 if $\mu > 0.2$;<br>**For Nittedal:** Equation S1.13 if $0.0001 \leq \mu \leq 0.2$<br>**For Trondheim:** Equation S1.14 if $0.0001 \leq \mu \leq 0.2$ | Own data |
| $r$ | Coefficient for energy reserve effect on predation threat | 0.05 | Model assumption |
| $C_{\text{competition}}(t)$ | Energetic competition cost | 0.0001 if $C_{\text{competition}} < 0.0001$ & 0.2 if $C_{\text{competition}} > 0.2$;<br>**For Nittedal:** Equation S1.19 if $0.0001 \leq C_{\text{competition}} \leq 0.2$<br>**For Trondheim:** Equation S1.20 if $0.0001 \leq C_{\text{competition}} \leq 0.2$ | Own data |
| $C_{\text{flight}}$ | Hourly flight cost | 0.615 g | Kurta *et al.* (1989)* |

\* Converted from original values, see Supplementary Materials 1

Environmental conditions like temperature and perceived predation threat (based on sunlight illumination-levels) are non-static across the daily cycle in the natural systems we base our model upon. therefore, we implemented time- or temperature-dependent variables when appropriate in order to capture the dynamic diurnal environments experienced by individuals at specific latitudes and times of year. Figure 1 shows the dynamic values of the baseline parametrizations, where time-dependent variables show fluctuations between sunrise (*t* = 1) and the following sunrise (*t* = 1 next day). Based upon meteorological data from Norwegian Centre for Climate Services, we specified six daily weather types as shown in Figure 1a: ‘very warm days’ (mean daily temperatures > 20°C and daily maximum temperatures > 26°C), ‘dynamic warm days’ (mean daily temperatures > 14°C and daily temperature ranges ≥ 6°C), ‘stable warm days’ (mean daily temperatures > 14°C and daily temperature ranges < 6°C), ‘dynamic cold days’ (mean daily temperatures ≤ 14°C and daily temperature ranges ≥ 3°C), ‘stable cold days’ (mean daily temperatures ≤ 14°C and daily temperature ranges < 3°C), and ‘very cold days’ (mean daily temperatures < 9°C, minimum daily temperature < 6°C and maximum daily temperature < 13°C), with a probability (*p*_*w*_) for the occurrence of each weather condition event. The different daily temperature cycles in the roost (Fig. 1a, dashed lines) were estimated using our own collected field-data (see Supplementary Materials 1), which further impact metabolic costs (Fig. 1b), while outside air temperatures (Fig. 1a, solid lines) affect prey availabilities (Fig. 1c). For simplicity, the estimated predation threat and competition cost in the model did not differentiate between day types (Fig. 1d), but we added a slight increase in predation threat with level of energy reserves in line with published estimates of mass-dependent flight costs (Anthony & Kunz 1977; Aldridge 1987; Witter & Cuthill 1993), which broadly tally with observations of heavy individual bats being more light avoiding (Speakman 1991). We thus specify a simple pattern of light-dependent predation threat from avian diurnal raptors and an interspecific competition cost to test if this is sufficient to generate realistic diurnal activity patterns in our model bat. We included two types of costs since the origins of the evolution of nocturnal activity in bats are currently unclear, potentially being related to niche-differentiation, mobbing or risk of hyperthermia, as well as diurnal predation threat (Rydell & Speakman 1995; Speakman 1995; Speakman *et al*. 2000).

**Figure 1:**
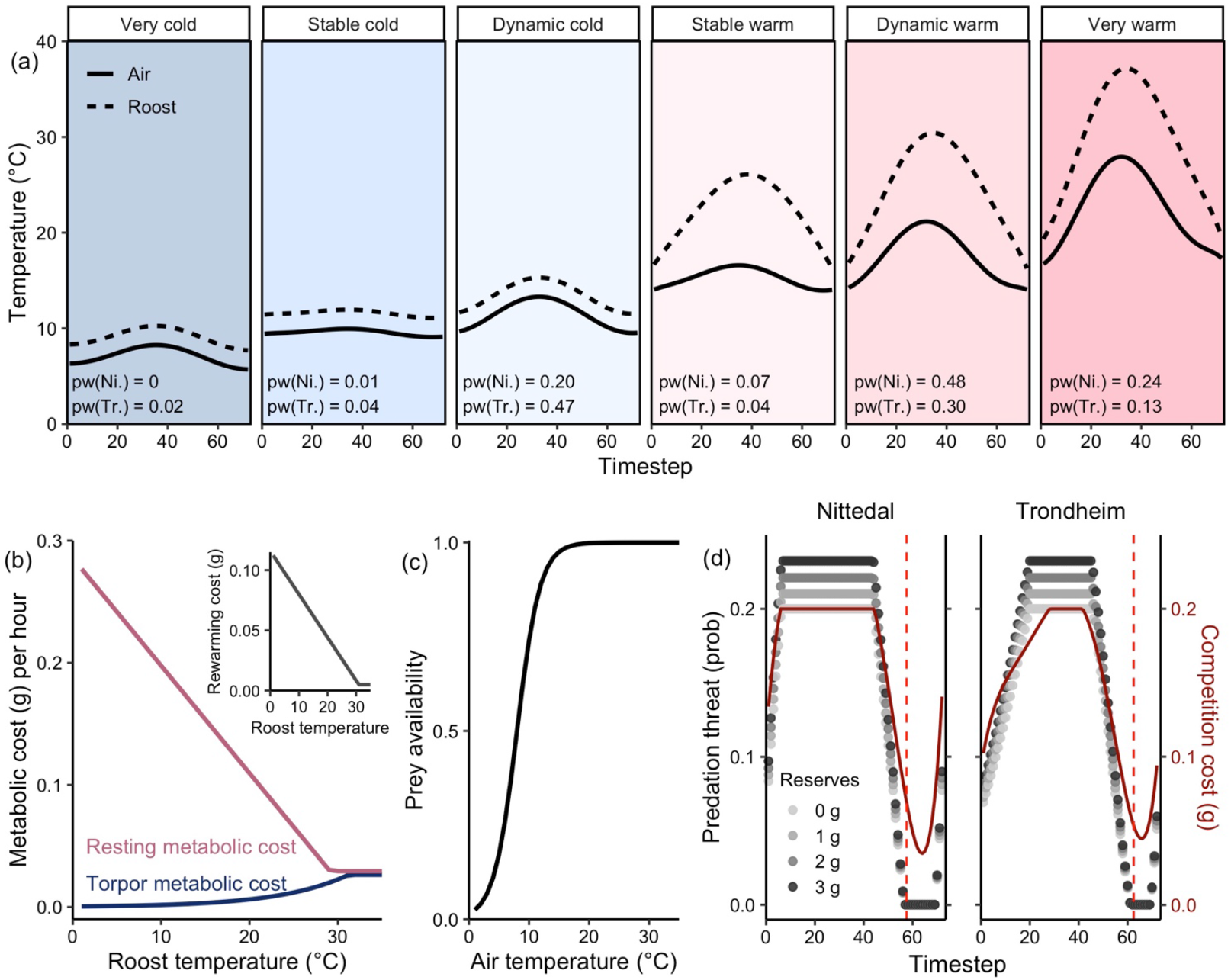
The dynamic parameter values used in the model (for the non-dynamic parameter values, see Table 1a and Table 1b). **(a)** Diurnal temperature fluctuations across six different day types for the outside air (solid lines, which impacts the insect abundance), and inside the roost (dashed lines, which affects the metabolic costs of resting, employing torpor and rewarming from torpor). p_w_ indicates the occurrence probability of each day type in Nittedal (denoted with Ni.) and Trondheim (denoted with Tr.) **(b)** Hourly metabolic costs shown as grams of energy reserves for behaviour 1 (torpor; dark blue line) and 2 (resting; dark pink line) as a function of roost-temperature. The smaller inset plot shows the linear function of the total rewarming cost against roost-temperature when transitioning from the state ‘torpid’ to the state ‘non-torpid’. **(c)** Prey availability as a function of the outside air temperature. **(d)** Diurnal variation in predation threat (grey-scale points) and energetic competition cost (dark red solid lines) calculated from light-measurements at the two locations Nittedal and Trondheim (see Supplementary Materials 1). The grey-scale indicates state-dependent (i.e. body mass) effects on predation threat, with individual energy reserves increasing from 0 to 3, and heavier individuals experiencing increased threat of predation. Dashed vertical red lines indicate timing of sunset at each of the two locations.

#### Model construction

The models were developed in R (version 4.0.3) and consist of two parts: a ‘backwards’ calculation of the optimal behaviour in each instance, in which we calculate the fitness of a hypothetical individual at a given time-of-day and physiological state across a wide range of possible scenarios; followed by a ‘forward’ iteration that simulates the best individual sets of diurnal sequences of these optimal decisions (see Clark & Mangel 2000). R-codes for the model are included as r-scripts in the Supplementary Materials. For our model framework, we used survival probability as the fitness proxy that the bats would attempt to maximize across the time horizon. The dynamic programming functions for our small bat in summer model were thus based on optimizing survival, *V*(*x, y, w, t, d*) (see equations 1 to 7 below), this being the maximum probability for a bat to survive from time period *t* until the last time period of the summer (*T, D*) plus their expected future fitness after summer S(*x*). Here, *X* represents the energetic state in terms of potential energy reserves with the current value *x, Y* is the body temperature state with the current state denoted as *y*, and *W* is the weather conditions with the current condition *w*. Therefore, *X* and *Y* function as the two individual state variables that are impacted by, as well as driving, individual decisions, while *W* represents the stochastic element of the model that affects costs and benefits associated with the different decisions.

In the model, bats can optimise their fitness by choosing between three activities (*a*): (1) employing torpor in the roost; (2) resting in the roost; or (3) foraging outside of the roost. The first two options have an associated predation threat of 0, but no food intake (Table 1b). The foraging option included a temperature-dependent food availability (λ(air(*w,t*)), Table 1b, Fig. 1c), but this activity also involved a predation threat and an energetic competition cost that varied across the daily cycle (Table 1b, Fig. 1d). We implemented an accelerating mass-dependent predation risk (*µ*(*t*)^*rx*^), as well as declining foraging success with increasing individual fat deposits (*p*_*food*_*(x)*), because we expect increasing flight costs and less agility in bats as individual body mass increases (Anthony & Kunz 1977; Aldridge 1987).

The two non-foraging activities, torpor and resting, differed in their respective temperature-dependent physiological costs (Fig. 1b). In our model, torpid individuals decreased their energy expenditure dramatically at low temperatures compared to resting individuals. However, the difference between the energetic costs decreased with increasing temperatures, until the curves flattened at a parallel level of torpor metabolic rate (TMR) being 90% of basal metabolic rate (BMR). Thus, a bat would at any temperature spend less energy employing torpor than resting, but the energetic benefit of torpor would decrease with increasing temperatures. In addition, we implemented a temperature-dependent cost of arousing from torpor (Fig. 1b), which was paid when an individual transitioned from the state ‘torpid’ to the state ‘non-torpid’. We expect there to be other potential associated physiological or ecological costs of prolonged periods spent in torpor (Humphries *et al*. 2003), but potential benefits of being awake might be more important to the individual bat during summer. We therefore implemented a fitness-benefit (*θ*) per time interval for resting activity versus being in torpor, which from our findings below we believe is an important aspect of the model and probably the natural world alike, although it has yet to be quantified in empirical studies.

The programming function is defined as:

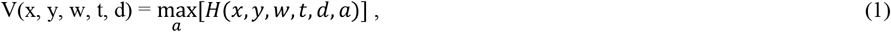

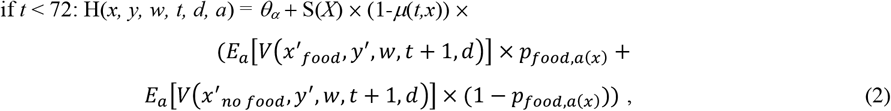

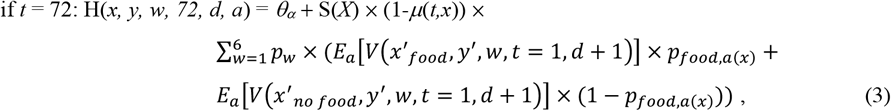

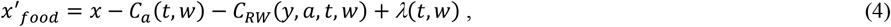

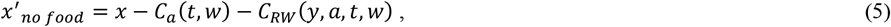

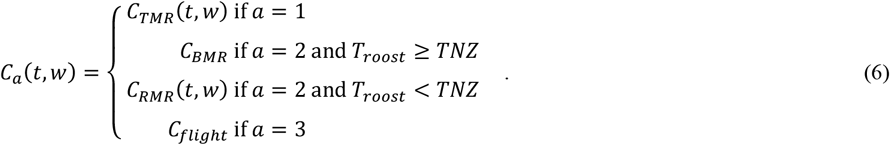

Each day (*d*) with its 72 timesteps (each *t* corresponding to 20 minutes) represented a daily cycle starting at sunrise and ending at the following sunrise. This daily cycle was chosen with regard to the nocturnal lifestyle of most bats, modelling a continuous timeline across 24-hours as we were interested in individual-level decisions during both day-time and night-time. We implemented a lower threshold value for energy reserves (*X*_*min*_), where individuals were treated as ‘dead’ and received no fitness if their energy reserves fell below the threshold at any point. We also applied a linearly decreasing survival probability below a given level of fat deposits (*X*_*dec*_). We thus defined the general survival probability (*S*(*x*)) of bats as:

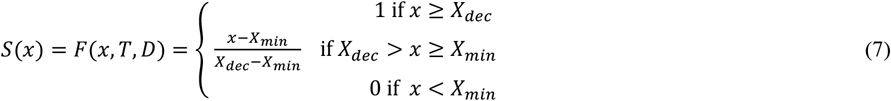

This was true for every *t* and therefore also functioned as the terminal reward (*F*(*x,T,D*)) in the model. As the end of July is still a long time before the onset of the hibernation season, we did not specify any additional requirement (i.e. above that needed for immediate survival) for over-winter energy reserves to be reached by the end of the time horizon. When energy reserves were between the pre-calculated discrete values, we used linear interpolation to calculate the corresponding fitness value as described in Clark and Mangel (2000) (see details in Supplementary Materials 2).

The optimal decision by the bat of which of the three activities (*a*) to carry out was calculated for every possible fat state (*x*), thermal state (*y*), weather (*w*) and timestep (*t*) within day (*d*), by going ‘backwards’ from the last time step modelled (*T*). These were saved and used for the ‘forward’ simulation of optimal sequence of individual behavioural routines for the whole month. For the forward simulations, we simulated 200 bats each starting in the non-torpid state with 1.5 grams of energy reserves at the first timestep in the model. The sequence of day types was randomised across the 30 days with the probability *p*_*w*_, with a new sequence generated for each simulated individual. Predation threat was included as a stochastic risk in the forward simulations, where individuals that experienced being predated upon were given a new and final energy state of −1, marking them as ‘dead’. Another stochastic elements in the forward simulation was the probability of finding food (*p*_*food*_), with a current foraging success of 0 if not successful.

### Empirical data comparisons

To test the qualitative and quantitative accuracy of our model predictions, we compared the model forward simulation results with our own data collected on northern bats at two locations in Norway: Nittedal and Trondheim. We used data collected from 7 non-reproductive northern bats in Trondheim (June 2020: 2; June 2021: 2; July 2021: 3) and on 2 in Nittedal (June 2019: 1, June 2021: 1). Permits to conduct the research were granted by the Norwegian Food Safety Authority (FOTS ID 23284) and the Norwegian Environment Agency (ref. 2018/4899).

Bats at each location were captured using mist-nets set up along tree-corridors and forest openings. Upon capture, individuals were weighed and fitted with a small transmitter (∼ 0.5 g, PIP31, Lotek Wireless Inc., Dorset, U.K.) that had been calibrated in a water bath (0°C to 45°C with stepwise increases of 5°C) prior to capture. We attached the tag by trimming a patch of fur from the dorsal region and applying a skin adhesive (B-530 Adhere Adhesive) on the transmitter before attaching it on the bat. The bats were thereafter released and tracked to their day-roosts using radiotelemetry. At the roosts, we put up remote loggers to record pulse-intervals from the transmitters every 10 minutes, which could afterwards be converted to skin temperatures (T_skin_) based upon the transmitter calibration.

To identify torpor bouts, we applied the method described in Fjelldal et al. (*under review*). This method consists of first determining a torpor onset (T_onset_) temperature value before differentiating each torpor bout into the three different phases of ‘torpor entry’, ‘stable torpor’ periods and ‘rewarming’. We therefore first calculated a species-specific T_onset_ value using the following equation (8), which was introduced by Willis (2007):

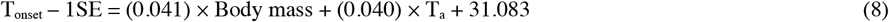

We used the mean values for body mass (mean capture weight: 8.7g ± 0.9) and environmental temperature (T_a_; collected by placing temperature-sensitive dataloggers outside each day roost) to calculate a species-specific T_onset_ value. Equation 4 is based upon true body temperature (T_b_) recordings with internal sensors, and as T_b_ – T_skin_ measurements usually is < 2ºC for small mammals (Audet & Thomas 1996; Barclay *et al*. 1996) we extracted 2 degrees from our T_onset_ value to get a torpor T_skin_ threshold value of 30.1ºC. We then extended the torpor bouts to include the full torpor entry and rewarming based upon the criteria described in Fjelldal et al. (*under review*). Bats with T_skin_ < 30.1ºC or in the torpor entry or rewarming phase were considered to be *torpid*. Bats in their roost with T_skin_ ≥ 30.1ºC and not in the torpor entry or rewarming phase were considered to be *resting*, and bats that were away from the roost (detected as a period of time with loss of the transmitter-signal) were considered to be *foraging*. These simplifications of behaviour were used to compare empirical results more easily with the simulated individual model sequences. Using the criteria for each of the six day types we categorized the days and nights (using the whole 24-hour temperature cycle also when determining nights) of the collected data. Sample sizes for each location were: N_night_ = 31 (7 ‘very warm’, 7 ‘dynamic warm’, 9 ‘stable warm’, 5 ‘dynamic cold’ and 3 ‘stable cold’) and N_day_ = 32 (12 ‘very warm’, 13 ‘dynamic warm’, 3 ‘stable warm’, 4 ‘dynamic cold’) in Trondheim, and N_night_ = 7 (5 ‘dynamic warm’, 1 ‘dynamic cold’ and 1 ‘stable cold’) and N_day_ = 7 (5 ‘dynamic warm’, 1 ‘dynamic cold’ and 1 ‘stable cold’) in Nittedal.

Based upon the sample sizes from each day type at the two locations, we ran simulations for each location with the same number of simulated individuals for each day type as listed above to compare with the empirical data. We then calculated the percentage of time spent on each of the three behaviours during the day (between sunrise and sunset) and night (between sunset and sunrise) for both simulated individuals and from the field data. The means of day-time or night-time percentages of expressed behaviour for empirical data and model simulations were then compared for each of the two locations using Welch two sample t-tests.

### Model predictions across a latitudinal gradient

To investigate limitations to bat species distributions at high latitudes, we calculated optimal decisions and simulated survival for 200 individuals throughout one summer month (30 days) at five locations across a latitudinal gradient: Nittedal (60.1°N), Trondheim (63.4°N), Bodø (67.3°N), Tromsø (69.6°N) and Gamvik (70.9°N). There are no observations of bats above 69.73°N in Norway (GBIF.org), and the northernmost location was therefore included to explore how the model predicted behaviour and survival at an extreme latitude. The only differences between the different location scenarios were probabilities of different day types, daily predation threat cycles and energetic competition cost cycles (see Supplementary Materials 1). For the predation-threat and energetic competition costs for Bodø, Tromsø and Gamvik, where we did not have our own collected light-levels, we used sun altitude obtained from the webpage of SunCalc to estimate light-levels based upon our collected data from Trondheim and Nittedal.

## Results

### General model simulation results

The final values in the baseline models for Nittedal and Trondheim were a mix of *a priori* estimates from the literature (metabolic costs, flight cost and the cost of arousing from torpor) and systematically tested values for parameters that have not been properly quantified (resting benefit, predation threat, energetic competition cost). Figures S3.1 to S3.11 in Supplementary Materials 3 show the outcome from the various model runs and subsequent simulations of individual decision-making (torpor, resting or foraging) across the daily cycle from sunrise to sunrise for each day type. This shows the results of systematic adjustments in resting benefits (*θ*), predation threats (*µ*) and competition costs (*C*_*competition*_) around their most likely values. Summarising across all individual iterations for each day-type, the middle panes in the figures thus represent the simulation outcome of baseline values from the model (Table 1a and Table 1b). Increasing or decreasing these values produce less biologically realistic diurnal patterns, as shown in the other panes of each figure. More details are given in Supplementary Materials 3.

The naturalistic diurnal scenario generated by the baseline parameter values was used to model the forward iterations, which reassuringly showed clear diurnal patterns in torpor, resting and foraging for small bats in summer across day types (Fig. 2). We modelled two main scenarios: one representing a low latitudinal (in this comparison) bat population from Nittedal and one representing a high latitudinal bat population from Trondheim. Before sunset on warm day types (*t* < 58 in Nittedal and *t* < 62 in Trondheim), most bats employed torpor in the morning (except bats in Nittedal on ‘very warm’ days, which most were awake and resting in the morning) and rewarmed to spend various amounts of time resting throughout the day, depending on day type and location (Fig. 2). In Trondheim on ‘stable warm’ and ‘dynamic warm’ days, most bats spent the middle of the day resting in their roost before re-entering torpor again in the afternoon. On ‘very warm’ days in Trondheim and on ‘stable warm’ and ‘dynamic warm’ days in Nittedal, most bats stayed awake and resting throughout the afternoon without re-entering torpor before leaving the roost to forage. These two decision patterns (employing torpor in the morning and afternoon with a resting period in the middle, or resting throughout the afternoon after a morning torpor bout) are the two most common daily torpor cycles found in free-ranging bats outside of the hibernation season, and are observed in bat species across climate zones (Fjelldal *et al*. 2022). During cold day types, the simulated bats spent the whole day torpid at both locations, and most of the bats even spent the whole night in the torpid state (Fig. 2). This daily pattern of spending the full day and following night torpid has been observed in several studies on various bat species, and is the second most frequently observed pattern in high-latitude northern bats (Fjelldal *et al*. 2022). Between sunset and the following sunrise during warm day types, however, simulated bats spent varying amounts of time foraging, resting and employing torpor, depending on day type and location (Fig. 2).

**Figure 2:**
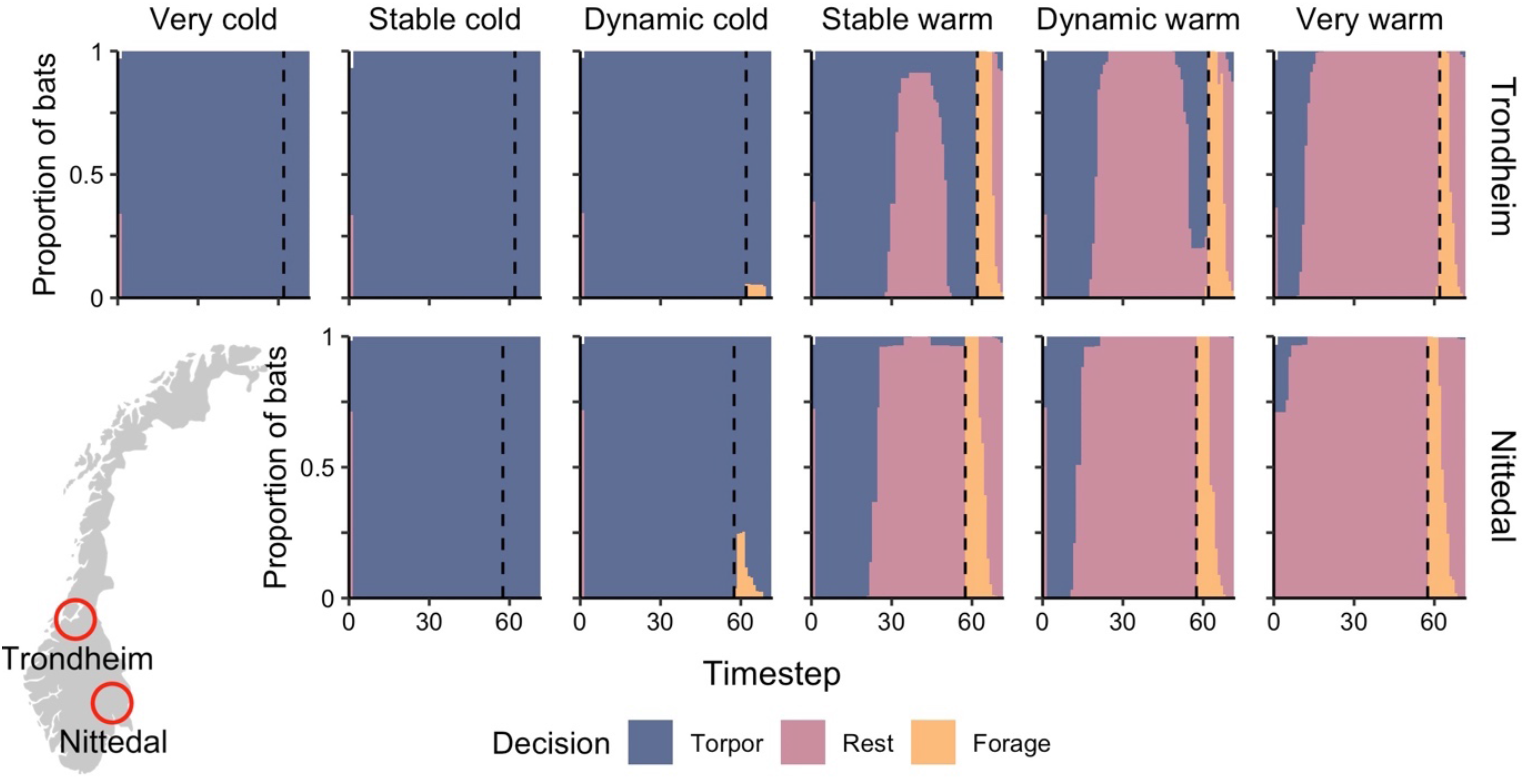
Frequency plots of the individual activities across the daily cycle (from sunrise to following sunrise) for different day types at the two locations Nittedal (lower panes) and Trondheim (upper panes). The plots summarise the decisions at each timestep made by 200 simulation forward model runs (individuals) across 30 days for each of the two locations. The bats could choose between employing torpor (blue), resting (pink) and foraging (yellow) at each timestep. Vertical dashed lines indicate the timing of sunset (at *t* = 58 in Nittedal and *t* = 62 in Trondheim). ‘Very cold’ day types are not shown for the Nittedal location because such days were never recorded during July in Nittedal and thus had an occurrence probability of 0 in the model.

The fluctuations in energy reserves across various day types (Fig. S4.1 in Supplementary Materials 4) demonstrate the amount of energy bats can save by spending a full day torpid during colder conditions, although the lack of foraging means that their reserves will keep depleting until a warmer day arrives. The average decline in individual energy reserves on ‘very warm’ days from timestep 1 to the timestep before sunset (*t* = 57 for Nittedal and *t* = 61 for Trondheim) was 0.76g ± 0.06g in Nittedal and 0.56g ± 0.06g in Trondheim. On ‘very cold’ days in Trondheim and ‘stable cold’ days in Nittedal (‘very cold’ days were never recorded in July at this location) the bats lost respectively 0.03g and 0.04g (with little to no variation) of their energy reserves between the same two timesteps, thus using a mere 5.7% and 5.3% of these reserves on the warmer days.

### Effects of environmental conditions and individual state

The combination of effects of photoperiod and state-dependency on the optimal diurnal routines for bats revealed strong responses in terms of the adaptive use of day-time torpor across day types (Fig. 3). On warm day types the percentage day-time torpor use decreased with increasing energy reserves (at the beginning of the day, *t* = 1) and with warmer weather conditions. On colder day types and in response to lower energy reserves on ‘stable warm’ days, bats expressed 100% torpor during day-time at each of the two locations. Photoperiod affected the day-time torpor expression on the warmer day types except for on ‘very warm’ days, with bats facing the lighter conditions of the Trondheim scenario generally expressing more torpor than conspecifics facing the darker Nittedal photoperiod, although the effect of energy reserves on torpor use were similar between locations (Fig. 3).

**Figure 3:**
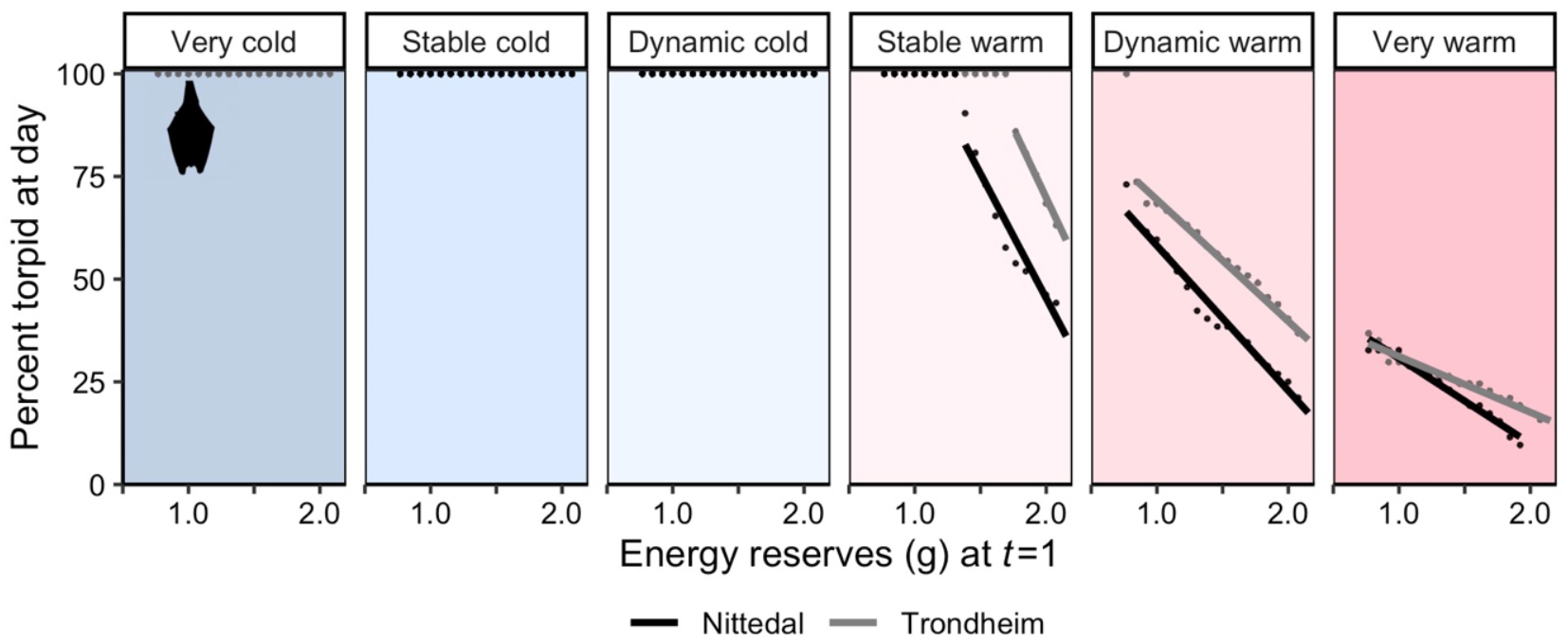
Model results for the percentage of expressed activity in ‘torpor’ in relation to individual fat reserves at the first timestep of the day (x-axis), temperature conditions (vertical panels left to right) and light photoperiod scenarios (dots per model run with best-fit lines for Nittedal in black and Trondheim in grey). Results are shown as separate dots for 100 one-day model runs for each location and fat reserve level scenario (only including fat reserves between the natural range of the initial simulation outputs, being 0.8 to 2.5g). Only percent of torpor expression is shown in this figure as the bats otherwise only rested during day-time. There are no datapoints from the Nittedal bat population on ‘very cold’ days because this day type had an occurrence probability = 0 for this scenario.

The optimal night-time decision to use torpor, resting or to forage was more complex in response to individual state and environmental conditions. Increasing individual energy reserves before sunset (i.e. at *t* = 58 for the Trondheim scenario, and *t* = 55 for the Nittedal scenario) had a generally negative effect on nightly foraging across all day types and locations (Fig. 4, top row), except for the two coldest day types on which all bats spent the whole night torpid. On ‘dynamic cold’ days, all bats used torpor at some point at night when they were not out foraging (Fig. 4, middle row), and individuals with greater energy reserves before sunset spent less time foraging and more time torpid than individuals with lower levels of energy reserves. For this day type, there was also no apparent effect of photoperiod (i.e. the Nittedal versus Trondheim scenarios) on either foraging or torpor (Fig. 4).

**Figure 4:**
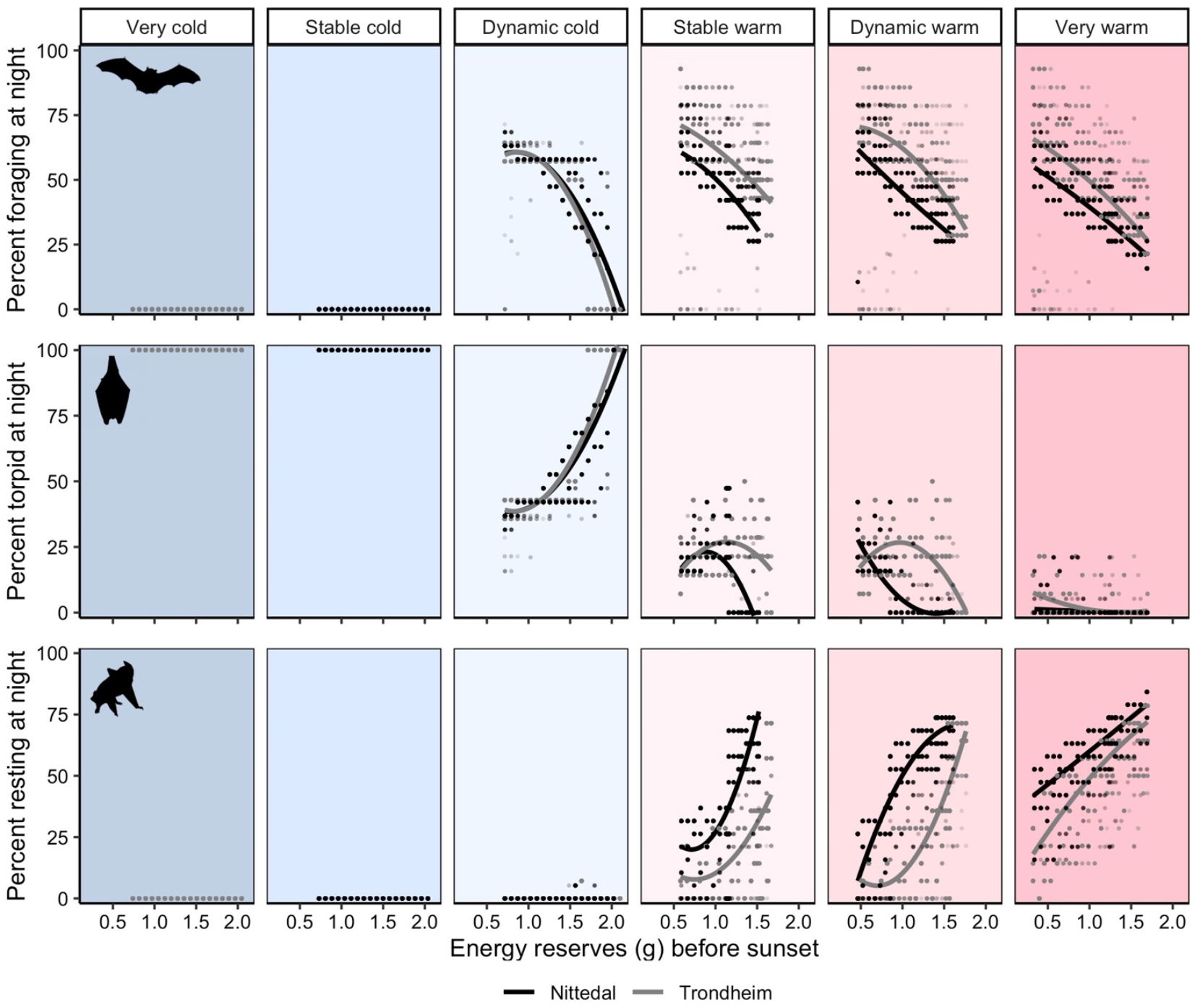
Model results for the percentage of expressed activity in ‘foraging’ (top panels), ‘torpor’ (middle panels) and ‘resting’ (lower panels) during night-time in relation to individual fat reserves before sunset (x-axes), temperature conditions (vertical panels left to right) and light photoperiod scenarios (dots per model run with best-fit polynomial lines for Nittedal in black and Trondheim in grey). Results are shown as separate dots for 100 one-day model runs for each location and fat reserve level scenario (only including fat reserves between the natural range of the initial simulation outputs for timestep 1, being 0.8 to 2.1g). Different fat reserves were implemented at timestep 1 and decreased across the day, depending on individual torpor vs. resting activities before night-fall. The large scattering of datapoints is due to implemented stochasticity in foraging success per timestep, affecting the continuous fat levels and thus the optimal decisions. There are no datapoints from the Nittedal bat population on ‘very cold’ days because this day type had an occurrence probability = 0 for this scenario.

Nightly foraging was not markedly different across light- and weather conditions on the warmer day types, although bats in Trondheim spent slightly more of the night foraging than conspecifics in Nittedal (Fig. 4, top row), while torpor use (Fig. 4, middle row) and resting (Fig. 4, bottom row) varied with both environmental conditions and location. The state-dependent effect of nightly torpor use changed direction from positive to negative with increasingly warmer day types in combination with higher energy reserves, but this shift happened at higher energy reserves and during warmer conditions for the Trondheim scenario than for the darker Nittedal scenario (Fig. 4, middle row). This shift was caused by the gradual increase in time spent resting at night (Fig. 4, bottom row), because individuals would decrease foraging time with increasing energy reserves, but at higher temperatures and energy reserves bats would exchange the remaining time spent torpid with time spent resting in the roost instead. However, while heavier bats facing the darker Nittedal light conditions would exchange all their torpor use for time resting, this was only the case for the very heavy bats during warmer day types when experiencing the lighter night-time conditions of Trondheim, except during nights of ‘very warm’ days. For ‘very warm’ day types, the effect of the photoperiod on the three different activities was more similar, indicating that at very high or very low temperatures the photoperiod is less important for nightly behavioural responses than at intermediate temperatures.

### Comparison of model predictions with empirical data

Overall, there was very good correspondence between our field data and the model predictions when using only the *a priori* parameter values from the literature and our own studies (Table 1a and Table 1b), plus the already established non *a priori* parameter values from the base model (Supplementary Materials 3). To provide more realistic quantitative comparisons, we matched sample sizes by using only the same number of model runs as we have samples from the field, considering the sample size for each day type in the model as in the empirical data.

#### Day-time torpor expression

Comparing the empirical data and model simulations of day-time torpor use at each location revealed strong similarities between observations and model results (Table S5.1 and Fig. S5.1 in Supplementary Materials 5). We performed Welch two sample t-tests between the field data and the model data for each location but found no significant difference between the means (for Nittedal: p-value = 0.86; for Trondheim: p-value = 0.38). However, although the sample sizes of bats at each day type at each location is similar between simulated model runs and empirical data, temperature cycles within day types still vary in the field data while it does not vary within day types in the model scenarios. Both the empirical data and the model simulation results revealed negative effects of increasing mean day-time air temperatures on day-time torpor expression (Fig. S5.1b), and although this effect was similar between the empirical data and model simulations for the Trondheim scenario, the model scenario for the Nittedal population showed an apparently stronger negative effect of the mean temperature than what was recorded in the empirical data. However, whether this is caused by discrepancies between the model and the field conditions, or if it is coincidental due to a small sample size at this particular location (N_day_ = 7), is not possible to determine here.

#### Night-time foraging, torpor and resting

For the nightly time allocation on foraging, torpor use and resting, the model simulations were again very similar to the empirical field data at the Trondheim location (Table S5.1 and Fig. 5a-c), revealing no significant differences between the means when tested with Welch two sample t-tests (foraging: p-value = 0.67; torpor use: p-value = 0.59; resting: p-value = 0.46). Larger differences were seen between the empirical data and the model simulations at the Nittedal location (Table S5.1 and Fig. 5a-c), although the means for torpor use and resting were not significantly different (torpor use: p-value = 0.78; resting: p-value = 0.11), while the means for time spent foraging revealed significantly more time spent on foraging in the empirical data than in the model simulations (p-value < 0.05). The temperature effect on the nightly behaviour showed positive impacts on the foraging and resting decisions and negative effects on the nightly torpor use; however, all these effects were apparently stronger in the model simulations than in the empirical data for the Nittedal scenario (Fig. 5d-f), which again is not possible to determine the causation of due to lack of field data from this location.

**Figure 5:**
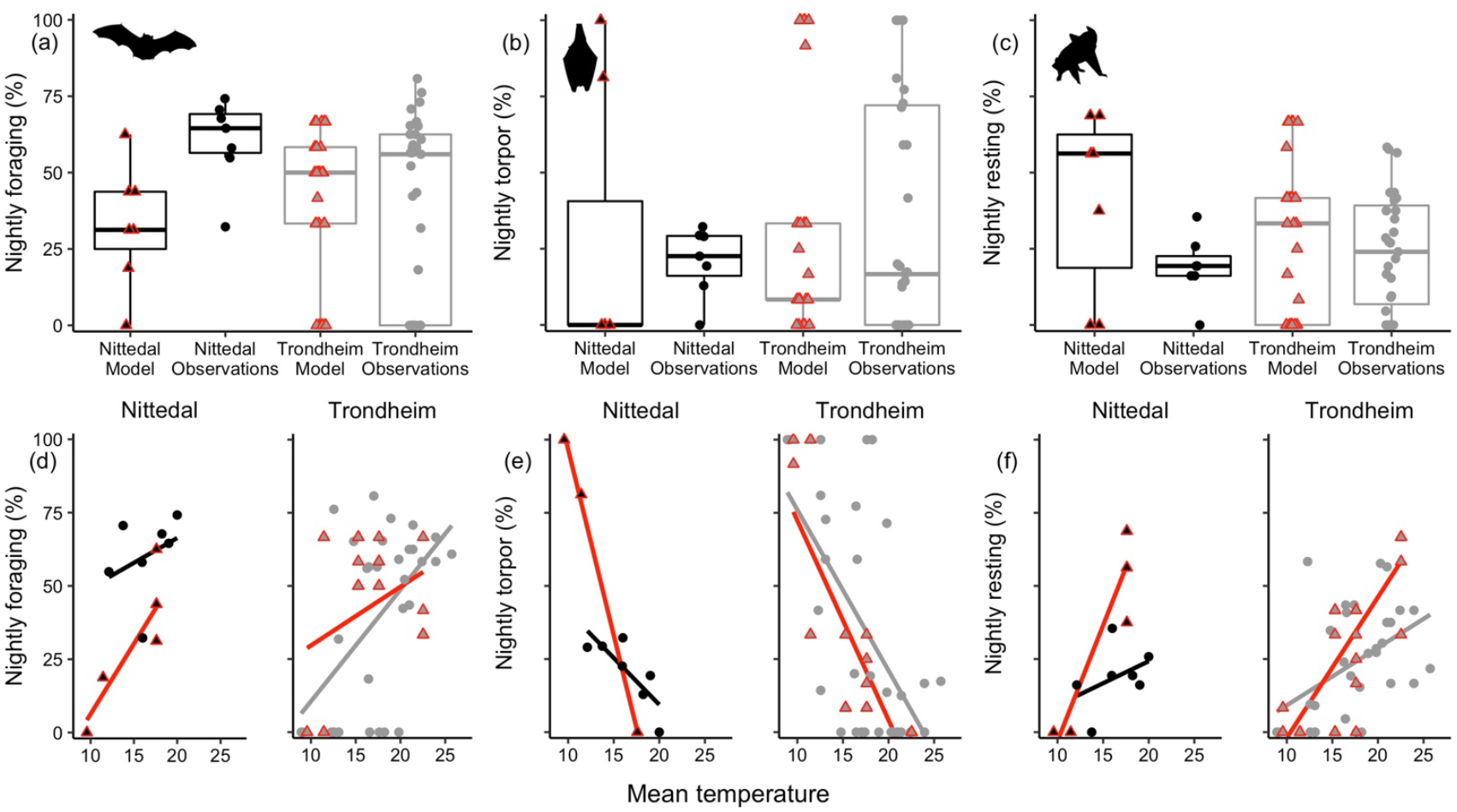
Comparisons of night-time ‘foraging’ (%), ‘torpor use’ (%) and ‘resting’ (%) from field derived data (round points) and from model simulations (triangle points with red outline) for the two locations of Nittedal (black points and boxes) and Trondheim (grey points and boxes). **(a)** Boxplots of nightly foraging between locations and data origin with boxes marking the data median and the 25^th^ and 75^th^ percentiles. **(b)** Boxplots of nightly torpor use between locations and data origin. **(c)** Boxplots of nightly resting between locations and data origin. **(d)** Positive effects of increasing mean daily temperatures on nightly foraging, both in the empirical data (round points and black or grey lines) and in data generated from matched simulated model runs (triangle points and red lines) for the two locations. **€** Negative effects of increasing mean daily temperatures on nightly torpor expression, both in the empirical data and in data generated from matched simulated model runs for the two locations. **(f)** Positive effects of increasing mean daily temperatures on nightly resting, both in the empirical data and in data generated from matched simulated model runs for the two locations.

#### Comparing model predictions across a latitudinal gradient

By first running ‘backwards’ calculations of optimal decisions followed by ‘forward’ simulating 200 individuals across one summer month at five different latitudes (see Fig. 6), we were able to reveal location-specific differences in both diurnal routines and optimal levels of each behaviour, as well as in survival probabilities. We explored potential limitations to location-specific survival by increasing and decreasing the intercept for daily temperature cycles by 2°C, simplifying scenarios for potential climate change effects at each location. These results are shown in panels in Figure 6 with the survival probability across 30 days from the baseline scenario for each location shown in the middle row of each set of panels. The results indicate that our simulated populations from Nittedal up to Tromsø were buffered against changes in temperatures by adjustments in their behavioural routines, although model bats in Tromsø showed signs of being at the very limit of their distributional range, with only slight effects on the summer survival if the mean temperature increased or decreased. Model bats in Gamvik were well beyond the distribution limit for survival given our model parameterization, and none of the temperature scenarios allowed them to survive past the 30 modelled days.

**Figure 6:**
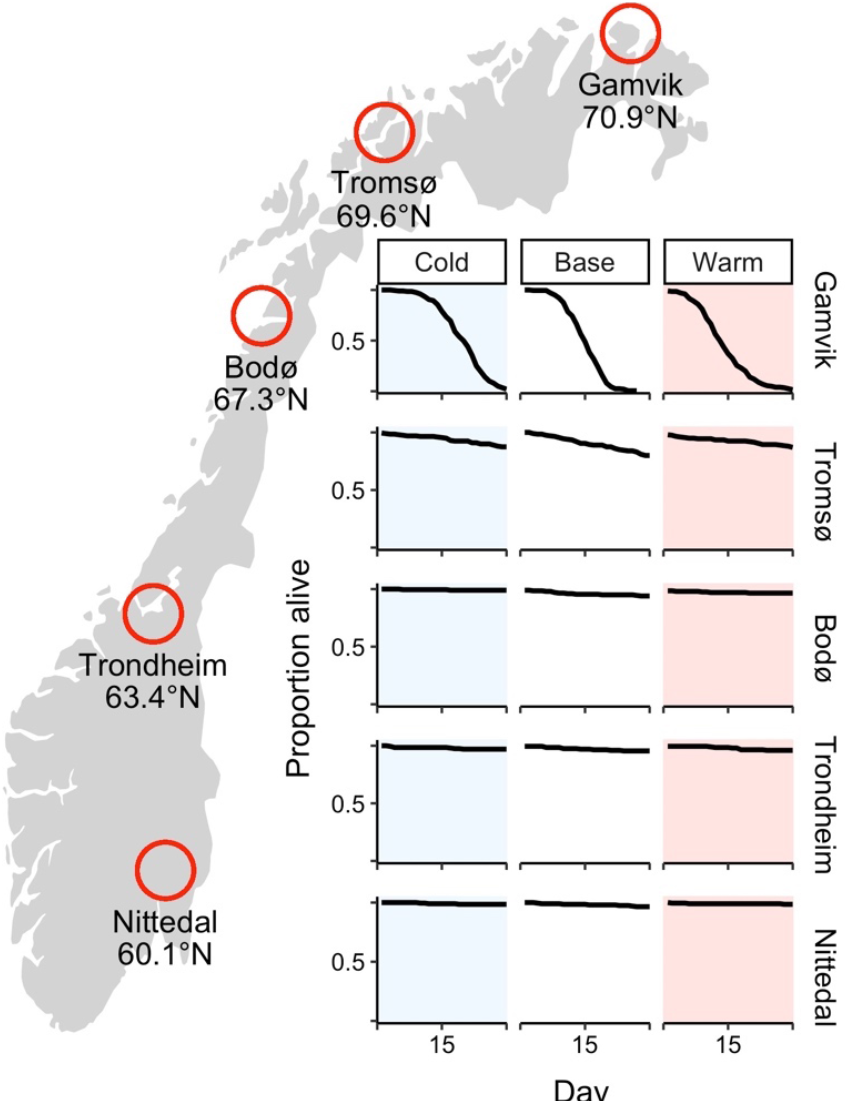
Survival probabilities from forward simulations of 200 individuals across one summer month (30 days) at five locations across a latitudinal gradient in Norway for different intercepts of the diurnal temperature cycle. Middle panes (white background) show scenarios with baseline values for each location, left panes show survival scenarios given a ‘cold’ temperature scenario (blue background), while right panes show scenarios given a ‘warm’ temperature scenario (pink background). The ‘cold’ temperature scenario corresponds to a decrease in the baseline temperature intercept of 2°C, whilst the ‘warm’ temperature scenario corresponds to an increase of 2°C.

When summarizing all individual simulations across the time horizon for all day types, the model bats in Nittedal showed the highest levels of resting during day-time on warmer day-types (Fig. 7). Model bats showed decreasing levels of day-time resting behaviour and increasing levels of torpor expression with increasing latitude on the same day-types. Given that the temperature cycles within day-types were the same across latitudes, the differences in daily torpor-strategies in Figure 7 must mostly be the result of variation in summer light levels, and hence foraging possibilities and individual energy reserves.

**Figure 7:**
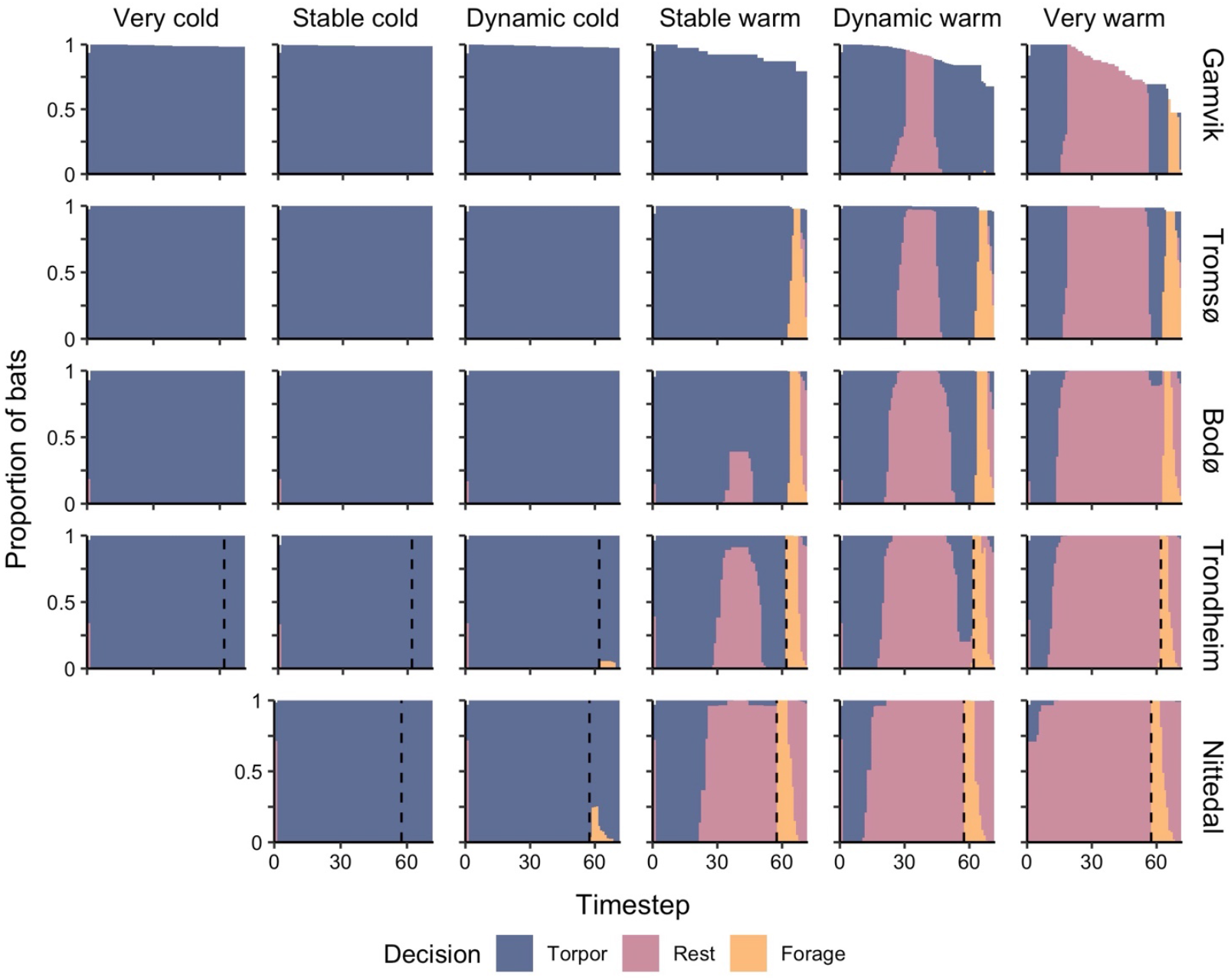
Summary plots across all day-types from forward simulations of 200 individuals at each of the five latitudes, illustrating the general behavioural patterns across the diurnal cycle at each location. Dark blue signifies when bats were expressing ‘torpor’, pink signifies ‘resting’, and yellow signifies ‘foraging’. Dashed vertical lines indicate sunset in Nittedal and Trondheim, while the sun in Bodø, Tromsø and Gamvik does not go down below the horizon during this summer month. ‘Very cold’ day types are not shown for the Nittedal location because such days were never recorded during July in Nittedal and thus had an occurrence probability of 0 in the model.

Diurnal fluctuations in energy reserves across warm day-types for each of the five locations showed that model bats in Nittedal spent more energy during day-time because they spent more time awake and resting, but they compensated for this mass loss by gaining markedly more foraging reserves at night than bats at any other location (Fig. S4.1 in Supplementary Materials 4). The average mass gain for foraging bats when comparing the body mass of timestep 57 to the maximum body mass of the following night on ‘very warm’ days was 1.1g ± 0.11 in Nittedal, 0.85g ± 0.15 in Trondheim, 0.81g ± 0.19 in Bodø, 0.84g ± 0.32 in Tromsø and 0.54g ± 0.17 in Gamvik. The greater foraging gain in Nittedal was due to the earlier sunset in the south of Norway, which allows bats to exit the roost to forage before the ambient temperature, and consequently the insect density, declines to its lowest levels due to lower overnight temperatures. Lighter nights further north also led the model bats at higher latitudes to stop their mass gain at lower levels as the predation risk increased with mass. Therefore, the bats in Nittedal were able to spend more time awake during the day, as they often had the chance to fully replenish their reserves during the following night. In contrast, model bats in Tromsø spent the whole day-time on even ‘stable warm’ days in torpor (Fig. 7), and thus lost less day-time body mass than the bats further south, but this was mostly because their foraging prospects were less good with limited time to forage and fewer insects available. For the northernmost location in Gamvik, most model bats did not forage until the occasional ‘very warm’ day, although no sequence of behaviours allowed them to survive throughout the whole summer month given the baseline parameterization values.

## Discussion

Our study describes the state-dependent optimal behaviour of small bats facing environmental and physiological challenges of high-latitude living in summer. By exploring the diurnal routines in the use of three activities (‘foraging’, ‘resting’, and ‘torpor’), we are able to show the various possible effects on individual behavioural strategies maximizing energy gain versus expenditure, and ultimately survival and limits to species northern geographical ranges. We can confirm that temperature cycles, and the strategic use of energy reserves, are important drivers of diurnal patterns in behavioural and physiological decision-making in bats, and specifically the strategic use of torpor. Such strong state-dependency is perhaps expected from the behavioural ecology literature (mostly on birds, Clark & Mangel 2000), but is not well explored in heterothermic endotherms and should contribute to our future understanding of strategic decisions in the order of Chiroptera.

A few empirical studies have demonstrated state-dependency regarding torpor expression in heterotherms, where animals with more energy reserves spent less time torpid, for example woodchucks (Marmota monax: Zervanos *et al*. 2014), dormice (Glis glis: Bieber *et al*. 2014), and bats (Myotis myotis: Wojciechowski *et al*. 2007; Myotis lucifugus: Matheson *et al*. 2010; Plecotus auritus: Sørås *et al*. 2022). Our model predictions of day-time torpor use are consistent with these findings, showing that individuals with lower energy reserves benefit to a greater extent from torpor energy savings, whilst individuals with greater energy reserves benefit from instead spending more time awake and resting. However, these responses are temperature dependent and vary most during night-time and the bats’ active period. This suggests that both the environmental conditions and any strategic diurnal activity patterns need to be taken into account when testing for state-dependent effects on behavioural and physiological decision-making.

The implementation of realistic and dynamic time- and/or temperature-dependent parameters in our model framework should result in quantitative predictions that match the relative costs and benefits for foraging, torpor and resting in our natural bat populations. As hoped, comparisons of day-time and night-time behavioural decisions in our empirical data on northern bats versus data generated from the model simulations showed strong similarities. However, the model underestimated the proportion spent foraging at the lower latitude location (Nittedal) compared to the field observations (Table S5.1, Fig. 5), but due to a small sample size for this population we were unable to determine if this was caused by actual differences between field observations and model parameters or just by chance in the sampling. The data collected at the higher latitude location (Trondheim) involved a much larger sample size and showed greater levels of variation in behavioural decisions across mean daily temperatures (Fig. S5.1b and 5d-f), which was also reflected in the model results, presumably caused by (stochastic) variation in individual energy reserves. The model comparisons for this location matched very well with the field observations, indicating that our model incorporates important drivers for general strategies in the behaviour and physiology of a range of individual bats in the studied species.

The forward simulation patterns of daily torpor use across the full Norwegian latitudinal gradient showed that, except for on ‘very warm’ days in the Nittedal scenario, all model bats employed torpor during the morning. On any of the warm day types in Nittedal, or on ‘very warm’ days in Trondheim and Bodø, the bats would mostly rewarm early and stay awake and resting until the evening, whilst bats on ‘stable warm’ or ‘dynamic warm’ days in Trondheim and Bodø, or on ‘dynamic warm’ and ‘very warm’ days in Tromsø or Gamvik, would re-enter torpor again in the afternoon and only rewarm again around sunset to forage (see Fig. 7). This closely matches northern bat behaviour recorded during the summer in Nittedal and Trondheim, corresponding to the two most commonly observed patterns in daily torpor use across bat species and climate zones (Fjelldal *et al*. 2022). These patterns involve either the ‘one-bout’ pattern (one torpor bout expressed daily) or the ‘W-shaped’ pattern (two torpor bouts expressed, one in the morning and one in the afternoon). Our model confirms that a main driver of these patterns is likely the diurnal temperature cycles within the roost (Turbill *et al*. 2008; this study), as well as state-dependency and prospect of successful foraging (Wojciechowski *et al*. 2007; this study). Bats in Nittedal were able to leave the roost to forage earlier than populations further north, and therefore profited from longer nights and higher prey availabilities, and this allowed them to spend more time awake during the day as their energy reserves could be fully replenished at night. Bats at higher latitudes adjusted their energy expenditure according to their more limited foraging prospects, and thus expressed more torpor across the daily cycle the further north they were. These strategic behavioural and physiological decisions regarding energy saving and foraging acquisitions are therefore driven by complex interactions between both current and prospective environmental conditions and individual energetic state (Wojciechowski *et al*. 2007; Fjelldal *et al*. 2021; this study).

Field-studies investigating latitudinal effects on torpor expression in free-ranging heterotherms have already identified patterns of increased daily torpor use with increasing latitudes (Fenn *et al*. 2009; Stawski 2012; Czenze *et al*. 2017). However, disentangling latitudinal effects from correlated temperature effects is challenging in field studies, although Czenze *et al*. (2017) found less torpor in a lower-latitude population of lesser short-tailed bats (*Mystacina tuberculata*) compared with higher-latitude living individuals across similar summer temperature conditions. This is in line with our model predictions, although Boyles *et al*. (2016) suggested that shifts in diet and foraging behaviour also could be used in response to environmental conditions at higher latitudes.

In our model framework, we only considered behaviours and trade-offs relevant for non-reproductive bats. Reproductive females face additional challenges during the summer as they go through highly energetically expensive periods of gestation and lactation while risking delays in the foetus development or reductions in milk production if they enter deep torpor to save energy (Racey & Swift 1981; Wilde *et al*. 1999). Pregnant and lactating bats therefore still use torpor, but only for short periods of time, and they maintain higher body temperatures in torpor than non-reproductive or post-lactating individuals (Dzal & Brigham 2013). A potential extension of our non-reproductive small-bat-in-summer model into one customised for reproductive female bats would thus also need to include not only the costs and benefits specifically related to reproduction and a modified measure of reproductive fitness plus survival, but also a multi-level body temperature state variable (e.g. see Brodin *et al*. 2017) and all of its consequences for foetal growth and survival, as opposed to our current ‘torpid’ versus ‘non-torpid’ two-level state variable. As our model does not include female reproduction, the results presented here regarding bat distributional ranges in Norway need to be interpreted as suitable environmental limits for general bat survival probabilities. Viable breeding colonies of bats have to navigate an even finer line between foraging, torpor and resting that encompasses the energetic requirements and costs of pregnancy and lactation (Kunz 1974; Kurta *et al*. 1989), and this will further affect any predictions of suitable environmental conditions for different species ranges across any latitudinal gradient.

Our non-reproductive model bat species is based upon quantified physiological estimates from several small-bodied bat species (see Supplementary Materials 1) and the distribution ranges predicted by our model correspond to the northernmost species in Norway, the northern bat. However, other bat species in Norway like the brown long-eared bat (*P. auritus*) or *Myotis spp*. are not found much further north than Trondheim (63.4°N). This suggests strong interspecific differences in distribution limits that require explanation. European insectivorous bat species show considerable interspecific variation in diet (Rydell 1989; Vaughan 1997), foraging behaviour (Norberg & Rayner 1987; Jones & Rydell 1994) and light sensitivity (Rydell *et al*. 1996; Duvergé *et al*. 2000), which will affect any theoretical assumptions regarding the factors limiting the northern distributional ranges of different species. However, the inclusion of quantitative estimates concerning species-specific diets and energetic requirements is perhaps something that could be investigated through further development of our model. Our general framework of optimal decision making of small-bats-in-summer should therefore contribute to the understanding of behavioural adaptations to high-latitude living in the order of Chiroptera and offer opportunities for further explorations of species- or location-specific behavioural decision-making in bats during summer.

## Conclusions

By developing a state-dependent stochastic dynamic programming model, we aimed to understand the strategic use of torpor in non-breeding bats facing the particular challenge of summer at high latitudes. Our simulations of general bat behaviour from our baseline model runs suggest that bats express the common adaptive diurnal routine in foraging and torpor versus resting observed in empirical studies as a response to particular combinations of temperature conditions, photoperiods (included as predation threat and energetic competition cost), individual energetic reserves and the anticipation of foraging profitability. Further simulating populations across a latitudinal gradient showed that the survival of bats inhabiting locations up to a certain latitude in Norway (≥ 67.3°N) was more or less buffered against environmental variation by such strategic use of torpor during the lightest summer month. Further north (69.6°N), the simulated bat populations seemed to be at the very edge of their distributional range limit as regards to summer survival, even with all their behavioural adjustments in torpor and foraging employed, which again aligns with the observed distributional range of northern bats in Norway.

Our model currently only considers the general survival challenges faced by non-reproductive small-bats-in-summer, but female reproductive bats face additional energetic challenges will affect the optimal use of torpor according to variation in environmental conditions. Distributional ranges of bats in Norway and elsewhere can only be properly investigated and understood by including the potential for breeding colony persistence, which should be possible via specific extensions of our model. As we demonstrate here, the powerful combination of state-dependent modelling and detailed empirical data collection can provide meaningful insight into environmental and physiological factors that drive behavioural decision-making of systems such as the small bat in summer, and a more complete understanding of the ecological limits of such species.

## Supporting information

Supplementary Materials (with R-code)

## Acknowledgements

We thank Thorbjørn Lundin and Myranda Murray for help with the initial coding of our earlier stochastic dynamic models, and Rune Sørås for the help with the empirical data collection in the field. MAF was supported by PhD funding (awarded to CS and JW) from the Department of Biology at the Norwegian University of Science and Technology (NTNU). JW and IIR were supported by the Norwegian Research Council Grant 223257 to the Centre for Biodiversity Dynamics (CBD) at NTNU.

## Author contributions (CRediT roles)

**Mari Aas Fjelldal**:Conceptualization, Software, Data curation, Formal analysis, Investigation, Methodology, Visualization, Writing - original draft. **Amandine Sophie Muller**: Software, Data curation, Methodology, Writing - Reviewing and Editing. **Irja Ida Ratikainen**: Software, Data curation, Methodology, Writing - Reviewing and Editing. **Clare Stawski**: Conceptualization, Funding acquisition, Supervision, Writing - Reviewing and Editing. **Jonathan Wright**: Conceptualization, Investigation, Methodology, Supervision, Writing - Reviewing and Editing.

## Data availability statement

R-codes for our model framework are included as r-scripts in the Supplementary Materials. The dataset used to compare model predictions and field observations are available from the corresponding author on reasonable request.

## References

Aldridge, H. (1987). Turning flight of bats. Journal of Experimental Biology, 128, 419–425.

Anthony, E.L. & Kunz, T.H. (1977). Feeding strategies of the little brown bat, Myotis lucifugus, in southern New Hampshire. Ecology, 58, 775–786.

Audet, D. & Thomas, D.W. (1996). Evaluation of the accuracy of body temperature measurement using external radio transmitters. Canadian Journal of Zoology, 74, 1778–1781. https://doi.org/10.1139/z96-196

Barclay, R.M., Kalcounis, M.C., Crampton, L.H., Stefan, C., Vonhof, M.J., Wilkinson, L. & Brigham, R.M. (1996). Can external radiotransmitters be used to assess body temperature and torpor in bats? Journal of Mammalogy, 77, 1102–1106. https://doi.org/10.2307/1382791

Bednekoff, P.A. & Houston, A.I. (1994). Optimizing fat reserves over the entire winter: a dynamic model. Oikos, 408–415.

Bergeson, S.M., Brigham, R.M. & O’Keefe, J.M. (2021). Free-ranging bats alter thermoregulatory behavior in response to reproductive stage, roost type, and weather. Journal of Mammalogy, 102, 705–717.

Bieber, C., Lebl, K., Stalder, G., Geiser, F. & Ruf, T. (2014). Body mass dependent use of hibernation: why not prolong the active season, if they can? Functional Ecology, 28, 167–177.

Boyles, J.G., Dunbar, M.B., Storm, J.J. & Brack, V. (2007). Energy availability influences microclimate selection of hibernating bats. Journal of Experimental Biology, 210, 4345–4350.

Boyles, J.G., Johnson, J.S., Blomberg, A. & Lilley, T.M. (2020). Optimal hibernation theory. Mammal Review, 50, 91–100. https://doi.org/10.1111/mam.12181

Boyles, J.G., McGuire, L.P., Boyles, E., Reimer, J.P., Brooks, C.A., Rutherford, R.W., Rutherford, T.A., Whitaker Jr, J.O. & McCracken, G.F. (2016). Physiological and behavioral adaptations in bats living at high latitudes. Physiology & Behavior, 165, 322–327.

Brodin, A. (2007). Theoretical models of adaptive energy management in small wintering birds. Philosophical Transactions of the Royal Society B: Biological Sciences, 362, 1857–1871.

Brodin, A., Nilsson, J.-Å. & Nord, A. (2017). Adaptive temperature regulation in the little bird in winter: predictions from a stochastic dynamic programming model. Oecologia, 185, 43–54.

Chaverri, G., Ancillotto, L. & Russo, D. (2018). Social communication in bats. Biological Reviews, 93, 1938–1954.

Clark, C.W. & Mangel, M. (2000). Dynamic state variable models in ecology: methods and applications. Oxford University Press.

Currie, S.E., Noy, K. & Geiser, F. (2015). Passive rewarming from torpor in hibernating bats: minimizing metabolic costs and cardiac demands. American Journal of Physiology-Regulatory, Integrative and Comparative Physiology, 308, 34–41. https://doi.org/10.1152/ajpregu.00341.2014

Czenze, Z.J., Brigham, R.M., Hickey, A.J. & Parsons, S. (2017). Stressful summers? Torpor expression differs between high-and low-latitude populations of bats. Journal of Mammalogy, 98, 1249–1255.

Duvergé, P.L., Jones, G., Rydell, J. & Ransome, R.D. (2000). Functional significance of emergence timing in bats. Ecography, 23, 32–40.

Dzal, Y.A. & Brigham, R.M. (2013). The tradeoff between torpor use and reproduction in little brown bats (Myotis lucifugus). Journal of Comparative Physiology B, 183, 279-288. 10.1007/s00360-012-0705-4

Fenn, A., Zervanos, S. & Florant, G. (2009). Energetic relationships between field and laboratory woodchucks (Marmota monax) along a latitudinal gradient. Ethology Ecology & Evolution, 21, 299–315.

Fjelldal, M.A., Sørås, R. & Stawski, C. (2022). Universality of Torpor Expression in Bats. Physiological and Biochemical Zoology, 95, 326–339.

Fjelldal, M.A., Wright, J. & Stawski, C. (2021). Nightly torpor use in response to weather conditions and individual state in an insectivorous bat. Oecologia, 197, 129–142.

GBIF.org (04 August 2022) GBIF Occurrence Download https://doi.org/10.15468/dl.g8m5nf.

Geiser, F. (2021). Ecological Physiology of Daily Torpor and Hibernation. Springer.

Geiser, F. & Brigham, R.M. (2000). Torpor, thermal biology, and energetics in Australian long-eared bats (Nyctophilus). Journal of Comparative Physiology B, 170, 153–162. https://doi.org/10.1007/s003600050270

Geiser, F. & Turbill, C. (2009). Hibernation and daily torpor minimize mammalian extinctions. Naturwissenschaften, 96, 1235-1240. 10.1007/s00114-009-0583-0

Houston, A., Clark, C., McNamara, J. & Mangel, M. (1988). Dynamic models in behavioural and evolutionary ecology. Nature, 332, 29–34.

Humphries, M.M., Thomas, D.W. & Kramer, D.L. (2003). The role of energy availability in mammalian hibernation: a cost-benefit approach. Physiological and Biochemical Zoology, 76, 165–179. https://doi.org/10.1086/367950

Jones, G. & Rydell, J. (1994). Foraging strategy and predation risk as factors influencing emergence time in echolocating bats. Philosophical Transactions of the Royal Society of London. Series B: Biological Sciences, 346, 445–455.

Kunz, T.H. (1974). Feeding ecology of a temperate insectivorous bat (Myotis velifer). Ecology, 55, 693–711.

Kurta, A., Bell, G.P., Nagy, K.A. & Kunz, T.H. (1989). Energetics of pregnancy and lactation in freeranging little brown bats (Myotis lucifugus). Physiological Zoology, 62, 804–818.

Landes, J., Pavard, S., Henry, P.-Y. & Terrien, J. (2020). Flexibility is costly: hidden physiological damage from seasonal phenotypic transitions in heterothermic species. Frontiers in Physiology, 11, 985. https://doi.org/10.3389/fphys.2020.00985

Lausen, C.L. & Barclay, R.M. (2003). Thermoregulation and roost selection by reproductive female big brown bats (Eptesicus fuscus) roosting in rock crevices. Journal of Zoology, 260, 235–244.

Liow, L.H., Fortelius, M., Lintulaakso, K., Mannila, H. & Stenseth, N.C. (2009). Lower extinction risk in sleep-or-hide mammals. The American Naturalist, 173, 264-272. 10.1086/595756

Lourenço, S.I. & Palmeirim, J.M. (2004). Influence of temperature in roost selection by Pipistrellus pygmaeus (Chiroptera): relevance for the design of bat boxes. Biological Conservation, 119, 237–243.

Matheson, A.L., Campbell, K.L. & Willis, C.K. (2010). Feasting, fasting and freezing: energetic effects of meal size and temperature on torpor expression by little brown bats Myotis lucifugus. Journal of Experimental Biology, 213, 2165-2173. 10.1242/jeb.040188

McNamara, J.M., Houston, A.I. & Lima, S.L. (1994). Foraging routines of small birds in winter: a theoretical investigation. Journal of Avian Biology, 287–302.

Norberg, U.M. & Rayner, J.M. (1987). Ecological morphology and flight in bats (Mammalia; Chiroptera): wing adaptations, flight performance, foraging strategy and echolocation. Philosophical Transactions of the Royal Society of London. B, Biological Sciences, 316, 335–427.

Parker, D.I., Lawhead, B.E. & Cook, J.A. (1997). Distributional limits of bats in Alaska. Arctic, 256–265.

Racey, P. & Swift, S.M. (1981). Variations in gestation length in a colony of pipistrelle bats (Pipistrellus pipistrellus) from year to year. Reproduction, 61, 123–129.

Ruf, T. & Geiser, F. (2015). Daily torpor and hibernation in birds and mammals. Biological Reviews, 90, 891-926. 10.1111/brv.12137

Rydell, J. (1989). Food habits of northern (Eptesicus nilssoni) and brown long-eared (Plecotus auritus) bats in Sweden. Ecography, 12, 16–20.

Rydell, J., Entwistle, A. & Racey, P.A. (1996). Timing of foraging flights of three species of bats in relation to insect activity and predation risk. Oikos, 243–252.

Rydell, J. & Speakman, J. (1995). Evolution of nocturnality in bats: potential competitors and predators during their early history. Biological Journal of the Linnean Society, 54, 183–191.

Skåra, K.H., Bech, C., Fjelldal, M.A., van Der Kooij, J., Sørås, R. & Stawski, C. (2021). Energetics of whiskered bats in comparison to other bats of the family Vespertilionidae. Biology Open, 10, bio058640.

Sørås, R., Fjelldal, M.A., Bech, C., van der Kooij, J., Skåra, K.H., Eldegard, K. & Stawski, C. (2022). State dependence of arousal from torpor in brown long-eared bats (Plecotus auritus). Journal of Comparative Physiology B, 1–13. 10.1007/s00360-022-01451-8

Speakman, J. (1991). Why do insectivorous bats in Britain not fly in daylight more frequently? Functional Ecology, 518–524.

Speakman, J. (1995). Chiropteran nocturnality. Symposia of the zoological society of London, pp. 187–201. London: The Society, 1960-1999.

Speakman, J. & Rowland, A. (1999). Preparing for inactivity: how insectivorous bats deposit a fat store for hibernation. Proceedings of the Nutrition Society, 58, 123–131.

Speakman, J., Rydell, J., Webb, P., Hayes, J., Hays, G., Hulbert, I. & McDevitt, R. (2000). Activity patterns of insectivorous bats and birds in northern Scandinavia (69 N), during continuous midsummer daylight. Oikos, 88, 75–86.

Speakman, J. & Thomas, M.D. (2003). Physiological ecology and energetics of bats. Bat Biology (eds T.H. Kunz & M.B. Fenton), pp. 430–492. University of Chicago Press.

Stawski, C. (2012). Comparison of variables of torpor between populations of a hibernating subtropical/tropical bat at different latitudes. Living in a seasonal world (eds T. Ruf, C. Bieber, W. Arnold & E. Millesi), pp. 99–108. Springer. 10.1007/978-3-642-28678-0_9

Stawski, C., Willis, C. & Geiser, F. (2014). The importance of temporal heterothermy in bats. Journal of Zoology, 292, 86–100.

Taylor, L. (1963). Analysis of the effect of temperature on insects in flight. The Journal of Animal Ecology, 99–117.

Turbill, C. (2008). Winter activity of Australian tree-roosting bats: Influence of temperature and climatic patterns. Journal of Zoology, 276, 285–290. 10.1111/j.1469-7998.2008.00487.x

Turbill, C., Kortner, G. & Geiser, F. (2008). Timing of the daily temperature cycle affects the critical arousal temperature and energy expenditure of lesser long-eared bats. Journal of Experimental Biology, 211, 3871–3878. 10.1242/jeb.023101

Vaughan, N. (1997). The diets of British bats (Chiroptera). Mammal Review, 27, 77–94.

Wilde, C.J., Knight, C.H. & Racey, P.A. (1999). Influence of torpor on milk protein composition and secretion in lactating bats. Journal of Experimental Zoology, 284, 35–41.

Wilkinson, G.S., Carter, G.G., Bohn, K.M. & Adams, D.M. (2016). Non-kin cooperation in bats. Philosophical Transactions of the Royal Society B: Biological Sciences, 371, 20150095.

Willis, C.K. (2007). An energy-based body temperature threshold between torpor and normothermia for small mammals. Physiological and Biochemical Zoology, 80, 643–651. https://doi.org/10.1086/521085

Winter, Y. & Von Helversen, O. (1998). The energy cost of flight: do small bats fly more cheaply than birds? Journal of Comparative Physiology B, 168, 105–111. 10.1007/s003600050126

Witter, M.S. & Cuthill, I.C. (1993). The ecological costs of avian fat storage. Philosophical Transactions of the Royal Society of London. Series B: Biological Sciences, 340, 73–92.

Wojciechowski, M.S., Jefimow, M. & Tęgowska, E. (2007). Environmental conditions, rather than season, determine torpor use and temperature selection in large mouse-eared bats (Myotis myotis). Comparative Biochemistry and Physiology, Part A: Molecular & Integrative Physiology, 147, 828–840. 10.1016/j.cbpa.2006.06.039

Zervanos, S.M., Maher, C.R. & Florant, G.L. (2014). Effect of body mass on hibernation strategies of woodchucks (Marmota monax). Oxford University Press.

