## Supplementary Materials (with R-code) for "The small-bat-in-summer paradigm: energetics and adaptive behavioural routines of bats investigated through a stochastic dynamic model"

### Supplementary Materials 1: Parameterization details and justifications

We constructed a stochastic dynamic model with a parameterization as close as possible to values from natural study systems. Explanations and justifications for all of these parameterization values can be found below.

#### 1.1. Quantified physiology

1.1.1 Cost of resting MR ($C_{RMR}$), basal MR ($C_{BMR}$) and torpor MR ($C_{TMR}$)

For the temperature-dependent resting metabolic rate (RMR) and basal metabolic rate (BMR), we used data from the small-bodied (~7 grams) Australian bat species *Nyctophilus geoffroyi* published in a study by Geiser and Brigham (2000). No latitudinal nor climate zone effects have been found to affect metabolic rates in the order of Chiroptera after correcting for body size and temperature (Speakman & Thomas, 2003; Cruz-Neto & Jones, 2006; Skåra et al., 2021; Fjelldal et al., 2022), which should result in these metabolic rates being representative for small bats. Furthermore, the thermoregulatory curve (Fig. 3b in Geiser and Brigham (2000)) and BMR ($1.36 \pm 0.17$ mL $O_2$ $g^{-1}$ $h^{-1}$) of *N. geoffroyi* closely resembles what we find in the northern bats (*unpublished data*), which is the focal species in our empirical data comparisons (see main text).

Equation S1.1 (from Geiser and Brigham (2000)) estimates the mass specific oxygen consumption (mL $O_2$ $g^{-1}$ $h^{-1}$) for temperatures below the lower critical temperature ($T_{LC}$) of 29.0 degrees:

$$\mathbf{C_{RMR}}: \quad \text{mL } O_2 \text{ g}^{-1} \text{ h}^{-1} = 11.69 - 0.35 \times \text{air temperature} \quad . \qquad \text{(S1.1)}$$

Temperatures above the $T_{LC}$ belong to the species' thermal neutral zone (TNZ; °C > 29°C) with a temperature-independent BMR as described in equation S1.2 (below). Given the high latitude and generally mild summers found at the study locations, we assume that the roosts in our model are not heating up to temperatures that lead to further increases in the MR, the highest modelled roost temperature being 35.4° (see section 1.2.2).

$$\mathbf{C_{BMR}}: \quad 1.36 \pm 0.17 \text{ mL } O_2 \text{ g}^{-1} \text{ h}^{-1} \quad . \qquad \text{(S1.2)}$$

For the torpor metabolic rate (TMR), we implemented the equation derived from measuring torpid Norwegian brown long-eared bats, presented in Fjelldal et al. (2022). We chose not to use the TMR curve presented in Geiser and Brigham (2000), because it lacked TMR values > 0.4 mL $O_2$ $g^{-1}$ $h^{-1}$, which meant that there was a large range between the highest measured TMR and the BMR that was unaccounted for. The TMR of Norwegian brown long-eared bats showed a similar curve to that of the

Australian eastern long-eared bat *Nyctophilus bifax* (Fjelldal et al., 2022), and could constitute a general representation for TMR in small bats. The TMR was found to increase exponentially with body temperature ($T_b$), following equation S1.3 (Fig. 1 in Fjelldal et al. (2022)). However, small bats have been found capable of saving large amounts of energy through torpor expressions even at high temperatures (Reher & Dausmann, 2021), and may stay torpid far into their TNZ (Sørås et al., 2022). Therefore, to avoid the exponential TMR curve increasing to unrealistically high values at high temperatures, we added a condition that TMR never increases above a value of $C_{BMR} \times 0.9$. This meant that a bat would always have the choice of saving energy by expressing torpor even at high temperatures:

$$\mathbf{C_{TMR}}: \quad mL\ O_2\ g^{-1}\ h^{-1} = \begin{cases} 0.0198 \times 1.137^{T_b} & \text{if } C_{TMR} < C_{BMR} \times 0.9; \\ C_{BMR} \times 0.9 & \text{if } C_{TMR} \geq C_{BMR} \times 0.9 \end{cases} \quad . \qquad (S1.3)$$

Using the values from Geiser and Brigham (2000) and Fjelldal et al. (2022), we estimated energy consumption and eventually the weight loss at different temperatures (see section 1.1.4 below for converting the MR into weight loss).

1.1.2 Rewarming cost ($C_{RW}$)

Heterotherms may save large amounts of energy by employing torpor, but face a temperature-dependent cost of active rewarming when choosing to exit this state (Turbill et al., 2008; Currie et al., 2015). The total cost of active arousals was quantified in *N. geoffroyi* at different environmental temperatures by Turbill et al. (2008), and described with the following equation:

$$\mathbf{C_{RW}}: \quad kJ = 0.84 - 0.026 \times \text{air temperature} \quad . \qquad (S1.4)$$

A second equation was presented in the original study where the net passive rewarming energy expenditure was included (see Fig. 5 in Turbill et al. (2008)); however, as each timestep in our model consists of a temperature-dependent TMR value this is already accounted for. We therefore chose to implement equation S1.4 as the quantified cost of arousal in our model, after converting it into weight loss (see section 1.1.4 below).

1.1.3 Flight cost ($C_{flight}$)

The energetic cost of flying was calculated for free-ranging little brown bats (*Myotis lucifugus*) by Kurta et al. (1989). The study calculated that the cost of foraging flight of a 9 g *M. lucifugus* was 4.46 kJ $h^{-1}$, a value which we implemented in our model. Another option would be to rearrange the equation provided by Thomas (1975) to estimate energy consumption during flight; however, this equation should be used with caution, particularly for species with small body masses (Kunz, 1980). In the study by Kurta et al. (1989), the calculated flight requirements for female brown bats were 13% lower than what Thomas' equation predicted. As the northern bats are approximately the same in size as the little brown bats, we used the calculation of their flight energy requirements in our model after converting the value into weight loss (see section 1.1.4 below).

##### 1.1.4 Converting physiological measurements into weight loss (g)

As our model represents unreproductive bats during summer, we assume that the main source of daily weight fluctuations is caused by stomach contents rather than by changes in fat reserves. According to the diet composition and conversion factors described in Kurta et al. (1989), one gram of fresh insects corresponds to 7.25 kJ of ingested energy, 6.12 kJ of assimilated energy, or 5.51 kJ of metabolized energy. We chose to use the estimates of ingested energy, where the energy density (7.25 kJ $g^{-1}$ wet mass) resemble the results found when performing a bomb calorimetry (Kunz, 1988). Knowing this, we converted the values obtained from respiratory studies into kJ and further into grams.

First, oxygen consumption was converted into $CO_2$ production: a typical insect diet consists of 70% water, 17.8 % protein, 4.6 % fat and 2.2 % carbohydrates (Kurta et al., 1989). Disregarding the water content, the diet consists of 72.4 % protein, 18.7 % fat and 8.9 % carbohydrates. In order to convert $O_2$ consumption to $CO_2$ production, the nutrient mixture in the diet should be considered due to its effect on the metabolism. We considered the following respiratory exchange ratios (RER = $VCO_2$ / $VO_2$) for the different components: 0.82 for protein; 0.70 for fat; and 1 for carbohydrates. Applying this to the mixed typical insect diet, we arrived at a RER = $(0.82 \times 0.724) + (0.70 \times 0.187) + (1 \times 0.089) = 0.81358$. We then utilized this RER value in order to estimate $VCO_2$ produced from $VO_2$ consumed.

Second, after estimating $VCO_2$, the $VCO_2$ was further converted into kJ. Each liter of $CO_2$ produced represents 27.2 kJ of assimilated energy (Kurta et al., 1989), and the estimates of $VCO_2$ were therefore first converted from mL to L and then further into kJ.

Finally, the estimated kJ was converted into loss of grams based on a conversion factor of 7.25 kJ = 1 gram wet mass (Kurta et al., 1989). The cost of RMR, BMR, TMR and the flight cost were calculated as hourly weight losses and was further converted into a cost per timestep in the model, while the cost of rewarming was kept as an overall total weight loss. Table S1.1 shows the conversion of each of the values from the original measurement to the estimated weight loss in grams.

**Table S1.1**: The step-wise conversion of each of the physiological measurements into weight loss in grams. Note that the units are different for $C_{RW}$ and $C_{flight}$ than what is described in the column header.

| Parameter | mL $O_2$ $g_{bat}^{-1}$ $h^{-1}$ | mL $CO_2$ $g_{bat}^{-1}$ $h^{-1}$ | kJ $g_{bat}^{-1}$ $h^{-1}$ | g ($g_{bat}^{-1}$ $h^{-1}$) |
|---|---|---|---|---|
| $C_{RMR}$ | $11.69 - 0.35 \times T_a$ | $9.51 - 0.285 \times T_a$ | $0.259 - 0.0078 \times T_a$ | $0.0357 - 0.0011 \times T_a$ |
| $C_{BMR}$ | 1.36 | 1.106 | 0.0301 | 0.0042 |
| $C_{TMR}$ | $0.0198 \times 1.137^{Tb}$ | $0.0161^{0.1284 \times Tb}$ | $0.0004^{0.1284 \times Tb}$ | $0.00006^{0.1284 \times Tb}$ |
| $C_{RW}$ | | | $0.84 - 0.026 \times T_a$ (kJ) | $0.11586 - 0.003586$ (g) |
| $C_{flight}$ | | | 4.46 (kJ $h^{-1}$) | 0.615 (g $h^{-1}$) |

#### 1.2. Quantified environmental conditions

1.2.1 Air temperature cycles across different day-type scenarios

In our model framework, we implemented dynamic time-dependent temperature cycles, modelling realistic environmental conditions bats may face across days and nights. We were interested in six general day types to catch variability in mean temperatures and daily temperature fluctuations alike: ‘very warm’ days; ‘dynamic warm’ days; ‘stable warm’ days; ‘dynamic cold’ days; ‘stable cold’ days; and ‘very cold’ days. To quantify the daily temperature cycles and the probability of each day type, we downloaded temperature data recorded every 10 minutes (if available, otherwise 1-hour recordings were used) from the Norwegian Centre for Climate Services webpage for the month of July across 9-10 years at each of the five locations (see table S1.2 for details). Not all stations or locations had temperature data recorded for July each year, which is why stations vary within location and years vary between locations.

We categorized each day (defining one day as the time between sunrise to the following sunrise) as one of six day types based on the following criteria:

1. ‘Very warm’ day: mean daily temperatures > 20°C and daily maximum temperatures > 26°C;
2. ‘Dynamic warm’ day: mean daily temperatures > 14°C and daily temperature ranges ≥ 6°C;
3. ‘Stable warm’ day: mean daily temperatures > 14°C and daily temperature ranges < 6°C;
4. ‘Dynamic cold’ day: mean daily temperatures ≤ 14°C and daily temperature ranges ≥ 3°C;
5. ‘Stable cold’ day: mean daily temperatures ≤ 14°C and daily temperature ranges < 3°C;
6. ‘Very cold’ day: mean daily temperatures < 9°C, minimum daily temperature < 6°C and maximum daily temperature < 13°C.

We converted the time of day into 72 timesteps between sunrise and the following sunset, corresponding to 20 minutes per timestep (following the timeframe used in our model framework – see main text), and fitted sixth order polynomial models for each day type to quantify the daily temperature cycles (Fig. S1.1). The polynomial models were fitted based on data from all five locations. We finally calculated the occurrence probability of each day type at each location based on the number of days in each category in relation to the total number of days (Fig. S1.2). The probabilities for each day type were accounted for when computing the optimal decisions in the backwards iterations by calculating the expected energy reserves and fitness values for each day type at timestep 1 for the next day, if the current timestep was the last timestep of the current day ($t = 72$). A weighted mean (based on the day type occurrence probabilities) for the expected fitness values for each decision was then calculated across day types for each energy reserve state level. In the forward simulation, the day types were included as stochastic variables by drawing day type sequences based on the occurrence probabilities.

**Table S1.2**: Details regarding the temperature data obtained for the month of July from the Norwegian Centre for Climate Services webpage.

| Location | Year | Station | Data measurement intervals |
|---|---|---|---|
| Nittedal | | | |
| | 2012 | SN18280 | 10 minutes |
| | 2013 | SN4460 | 1 hour |
| | 2014 | SN18280 | 10 minutes |
| | 2016 | SN18280 | 10 minutes |
| | 2017 | SN18280 | 10 minutes |
| | 2018 | SN18280 | 10 minutes |
| | 2019 | SN18280 | 10 minutes |
| | 2020 | SN18280 | 10 minutes |
| | 2021 | SN18280 | 10 minutes |
| Trondheim | | | |
| | 2012 | SN68860 | 1 hour |
| | 2013 | SN68860 | 1 hour |
| | 2014 | SN68175 | 10 minutes |
| | 2015 | SN68860 | 1 hour |
| | 2016 | SN68175 | 10 minutes |
| | 2017 | SN69035 | 10 minutes |
| | 2018 | SN68175 | 10 minutes |
| | 2019 | SN68175 | 10 minutes |
| | 2020 | SN68175 | 10 minutes |
| | 2021 | SN68175 | 10 minutes |
| Bodø | | | |
| | 2011 | SN82290 | 10 minutes |
| | 2012 | SN82290 | 10 minutes |
| | 2013 | SN82290 | 10 minutes |
| | 2014 | SN82290 | 1 hour |
| | 2015 | SN82290 | 10 minutes |
| | 2016 | SN82220 | 10 minutes |
| | 2017 | SN82220 | 10 minutes |
| | 2018 | SN82220 | 10 minutes |
| | 2019 | SN82220 | 10 minutes |
| | 2020 | SN82220 | 10 minutes |
| Tromsø | | | |
| | 2011 | SN90490 | 1 hour |
| | 2012 | SN90490 | 1 hour |
| | 2013 | SN90490 | 1 hour |
| | 2014 | SN90490 | 1 hour |
| | 2015 | SN90490 | 1 hour |
| | 2016 | SN90490 | 1 hour |
| | 2017 | SN91180 | 10 minutes |
| | 2018 | SN91180 | 10 minutes |
| | 2019 | SN91180 | 10 minutes |
| | 2020 | SN91180 | 10 minutes |
| Gamvik | | | |
| | 2012 | SN96310 | 1 hour |
| | 2013 | SN96310 | 1 hour |
| | 2014 | SN96310 | 1 hour |
| | 2015 | SN96310 | 1 hour |
| | 2016 | SN96310 | 1 hour |
| | 2017 | SN98265 | 10 minutes |
| | 2018 | SN98265 | 10 minutes |
| | 2019 | SN96310 | 1 hour |
| | 2020 | SN98265 | 10 minutes |
| | 2021 | SN96310 | 1 hour |

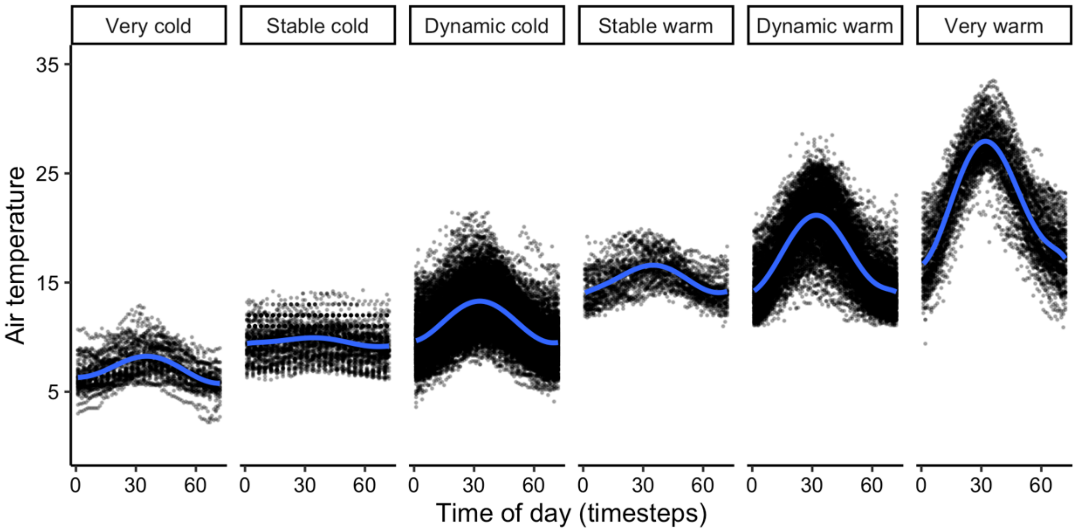


**Figure S1.1**: Daily temperature cycles for each day-type from sunrise (timestep = 1) to the last timestep before the following sunrise (timestep = 72). Black dots are the recorded temperatures across locations and years during the full month of July, while blue lines show fitted six-degree polynomial models for each day-type.

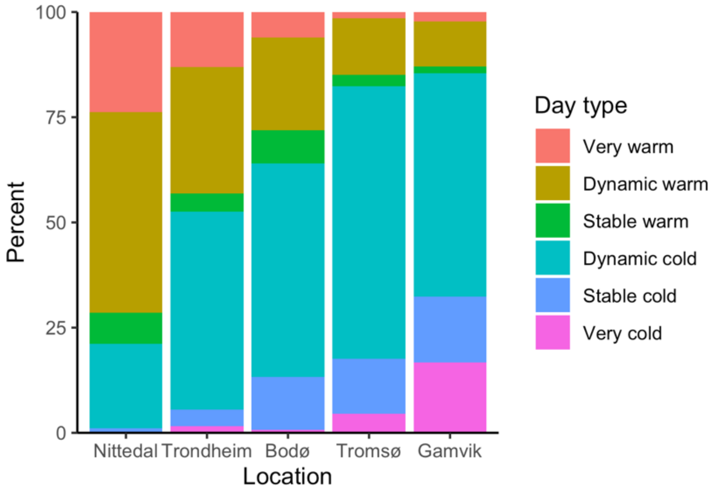


**Figure S1.2:** Occurrence probabilities of each day-type during July across locations. The exact probabilities for, respectively, 'very warm' days, 'dynamic warm' days, 'stable warm' days, 'dynamic cold' days, 'stable cold' days and 'very cold' days at each location were: 0.0, 0.01, 0.20, 0.07, 0.48 and 0.24 in Nittedal; 0.02, 0.04, 0.47, 0.04, 0.30 and 0.13 in Trondheim; 0.01, 0.12, 0.51, 0.08, 0.22 and 0.06 in Bodø; 0.04, 0.13, 0.65, 0.03, 0.13 and 0.02 in Tromsø; 0.17, 0.16, 0.53, 0.01, 0.11 and 0.02 in Gamvik.

Equations S1.5 to S1.10 below describes the sixth-degree polynomial model equations for the temporal (dependent on timestep (*t*)) air temperature cycles for each day type:

'Very warm' days: $16.58°C + 0.12 \times t + 0.024 \times t\text{^}2 - 8.99 \times 10^{-5} \times t\text{^}3 - 2.95 \times 10^{-5} \times t\text{^}4 + 6.01 \times 10^{-7} \times t\text{^}5 - 3.38 \times 10^{-9} \times t\text{^}6$ (S1.5)

| | | |
|---|---|---|
| 'Dynamic warm' days: | $14.11°C + 0.11 \times t + 0.009 \times t^2 + 2.53 \times 10^{-4} \times t^3 - 2.56 \times 10^{-5} \times t^4 + 4.47 \times 10^{-7} \times t^5 - 2.36 \times 10^{-9} \times t^6$ | (S1.6) |
| 'Stable warm' days: | $13.95°C + 0.11 \times t – 0.007 \times t^2 + 5.62 \times 10^{-4} \times t^3 - 1.82 \times 10^{-5} \times t^4 + 2.30 \times 10^{-7} \times t^5 - 1.00 \times 10^{-9} \times t^6$ | (S1.7) |
| 'Dynamic cold' days: | $9.62°C + 0.04 \times t + 0.008 \times t^2 - 8.43 \times 10^{-5} \times t^3 - 6.03 \times 10^{-6} \times t^4 + 1.24 \times 10^{-7} \times t^5 - 6.43 \times 10^{-10} \times t^6$ | (S1.8) |
| 'Stable cold' days: | $9.39°C + 0.03 \times t – 0.004 \times t^2 + 2.89 \times 10^{-4} \times t^3 - 8.41 \times 10^{-6} \times t^4 + 1.03 \times 10^{-7} \times t^5 - 4.43 \times 10^{-10} \times t^6$ | (S1.9) |
| 'Very cold' days: | $6.31°C + 0.005 \times t + 0.002 \times t^2 + 1.80 \times 10^{-4} \times t^3 - 9.02 \times 10^{-6} \times t^4 + 1.27 \times 10^{-7} \times t^5 - 5.68 \times 10^{-10} \times t^6$ | (S1.10) |

1.2.2 Roost temperature cycles across different day-type scenarios

Roost temperatures can be markedly different from outside air temperatures, depending on roost characteristics (e.g. Kerth et al., 2001; Lausen & Barclay, 2003; Lourenço & Palmeirim, 2004; Stawski et al., 2008; Michaelsen et al., 2014). Our model framework incorporates temperature-dependent physiological energetics (i.e. metabolic rates and rewarming costs), and given our assumptions that bats could only employ torpor or the resting behaviour while inside their respective roosts we attempted to implement realistic roost temperature cycles that depended on, but that were not necessarily identical to the outside air temperature. For this model framework, we used our own estimates of roost temperature, as described below.

All our radio-tagged northern bats from the field (see details in the main text Methods section) roosted under the roofs of houses or sheds during our data collection period. Upon losing their transmitters, three of these were dropped while the bats were still in their roosts and kept on recording temperature data for a prolonged period of time, logging the roost temperature. Although we could not verify the exact location of the tags within the roosts, we assumed they were reasonably close to where the bats had been roosting, and thus recorded temperature conditions representing the environment experienced within the roost. The daily temperatures recorded in the roosts showed large fluctuations on most days, with the total temperature range recorded being 9.6°C to 42.1°C (Fig. S1.3).

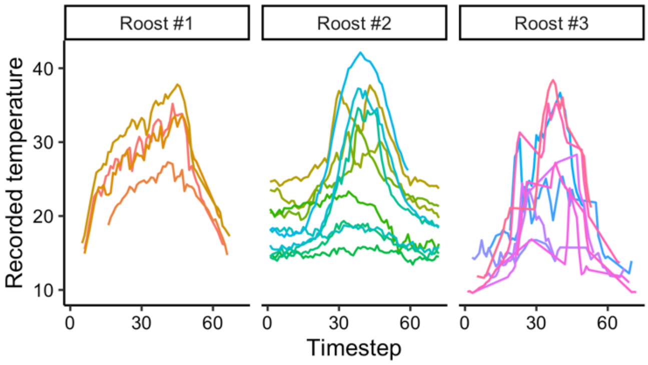


**Figure S1.3**: Recorded roost temperature across the daily cycle (between sunrise and the following sunrise, one timestep corresponding to 20 minutes) for three different roosts, all located under the roofs in houses. Each coloured line represents measurements across one day.

A total of 23 days of roost temperature data were recorded across the three transmitters lost in the roosts. Simultaneous outside air temperature data were recorded every 10 minutes using small heat-sensitive data loggers (0.5°C, DS 1921G Thermochron iButtons, Maxim Integrated Products, Inc., Sunnyvale, CA, USA) placed inside cardboard cups and hung from tree branches ~1.5 meters above the ground outside of each roost. By subtracting the measured air temperature from the roost temperature, we obtained a measure of the difference between air temperature and roost temperature across the 23 days of data.

Based on the outside air temperature, we defined each day as one out of six day types (as explained in section 1.2.1 above). Because we did not have enough data to determine the different roost temperature trajectories for all six day types, we made a simplified assumption of categorizing 'cold' day types ('very cold', 'stable cold' and 'dynamic cold') and 'warm' day types ('very warm', 'stable warm' and 'dynamic warm'), resulting in 6 days in the 'cold days' category and 17 days in the 'warm days' category. The daily trajectories of temperature differences between roost temperature and air temperature are shown in figure S1.4, illustrating how the roosts warmed up markedly more than the outside air temperature on 'warm' days (likely due to the sun warming up the roofs under which the bats were roosting), while on 'cold' days the roosts had a slightly higher but stable temperature compared to the outside air temperature. For simplicity (and due to the low sample size for 'cold' days) we defined roost temperature on any of the 'cold' day types as a constant 2 degrees higher than the outside air temperature in our model, while the roost temperature on 'warm' day types were defined with a 3$^{rd}$ degree polynomial model (as shown in Fig. S1.4; equation specified in Table 1a in Methods).

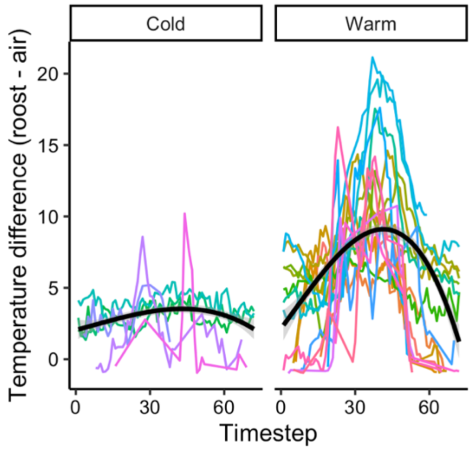


**Figure S1.4**: Temperature differences between the recorded roost temperature and the outside air temperature across the daily cycle (between sunrise and the following sunrise, one timestep corresponding to 20 minutes) for 'cold' and 'warm' day types. Black lines show fitted three-degree polynomial models for each day-type category.

<u>1.2.3 Food availability</u>

For our study system, we used insectivorous bats as model organisms. About 70% of all bat species feed on insects (Jones & Rydell, 2005), a food source that is heavily affected by seasonal and environmental fluctuations, and particularly air temperature (Anthony & Kunz, 1977; Speakman et al., 2000; Stawski, 2012). For our model, we considered the findings by Speakman et al. (2000) where variation in bat activity and prey abundance were correlated with air temperature, linking the insect abundance to daily temperature cycles (see Fig. 2 in Speakman et al. (2000)). Using this figure as our reference point, we created the following equation (S1.11) regarding temperature-dependent insect abundance:

$$\text{Food abundance (proportion):} \quad 1 / (1 + \exp( -0.524 \times (T_a - 8)) \qquad \text{(S1.11)}$$

This equation produces a logistic curve for the temperature-dependent proportion of insect abundance (from 0-1; see Fig. 1c in the main text Methods). For our model, we further needed to include a net foraging intake to provide the energetic value the bats could obtain at each time-step when foraging. In the study by Sørås et al. (2022), brown long-eared bats were found to increase their body mass at capture with 0.009 grams per minute after sunset, indicating increased gut fill with increased foraging time. Converting this into an hourly value resulted in an expected increase in body mass of 0.54 g $h^{-1}$. For the purpose of our model, we were interested in including a net foraging intake, which meant accounting for flight costs and potential temperature effects on the foraging success. We therefore added the presumed $C_{flight}$ of 0.615 g $h^{-1}$ to the hourly body mass increase and, after calculating the mean nightly $T_a$ for the capture nights at the study location (*unpublished data*), used the overall average nightly temperature (12.18°C) to account for expected food abundance using equation S1.11. With an estimated food abundance of 0.9 at 12.18°C, we arrived at a net foraging intake of ~1.3 g $h^{-1}$. The potential energy gain (α in the model) was tested in early models with three different values (1 g, 1.3 g, 1.6 g) to explore model sensitivity and how slightly higher and lower values altered the decision-pattern of bats in summer, and this verified that the calculated baseline value of 1.3 g $h^{-1}$ generated realistic scenarios.

##### 1.2.4 Light-dependent predation threat

Predation threat represents one of the main explanations for why bats are nocturnal creatures, although bat-catches by vertebrate predators seem to be mainly opportunistic (Lima & O'Keefe, 2013). Nevertheless, the risk of being preyed upon by avian raptors has been estimated to be 100 to 1000 times higher during daylight compared with nocturnal hours (Speakman, 1991a; 1991b; 1995). We therefore assumed a light-dependent predation threat, using our own collected light measurements (Fig. S1.5a) to estimate predation threats (Fig. S1.5b) across the daily cycle in Nittedal and Trondheim. Light conditions were measured in lux using an illuminance UV recorder (TR-74Ui data logger). These light recordings were then log-scaled (due to the massive range in measured lux), before they were converted to fit as a realistic predation risk ranging between 0 and up towards 1. The equation below (S1.12) shows the conversion from measured lux to an estimated predation risk:

$$\text{Predation threat:} \quad = \frac{(\log(\text{lux}))^3}{270} \quad . \qquad \text{(S1.12)}$$

We fitted fifth order polynomial models to represent the fluctuations in light-dependent predation threats throughout the day, but specified a minimum threshold-value of 0.0001 as some of the estimated values from the fitted models were negative during night-time, and an upper threshold-value of 0.2 because of assumptions that the predation threat would be light-independent above certain light-levels. For simplicity, the estimated predation threat in the model did not differentiate between the six different day types.

As we did not have light-measurements from higher latitudes than Trondheim, we downloaded sun altitude data per 20 minutes for July at each of the five locations (Nittedal, Trondheim, Bodø, Tromsø and Gamvik), obtained through the webpage SunCalc. We then used the light-measurements from Nittedal and Trondheim to estimate light-levels at the three remaining locations, averaging light-levels measured in Nittedal and Trondheim for similar sun altitude levels across the daily cycle for each of the three other locations. We then converted the estimated light-level cycles into predation threat values for Bodø, Tromsø and Gamvik, as described above. Figure S1.6 shows the estimated predation threat cycles for each of the five locations (not showing the added effect of energy reserves on predation threat, see Methods and Fig. 1d). Fifth degree polynomial model equations describing the temporal predation threat for each location are defined with equations S1.13 to S1.17 below.

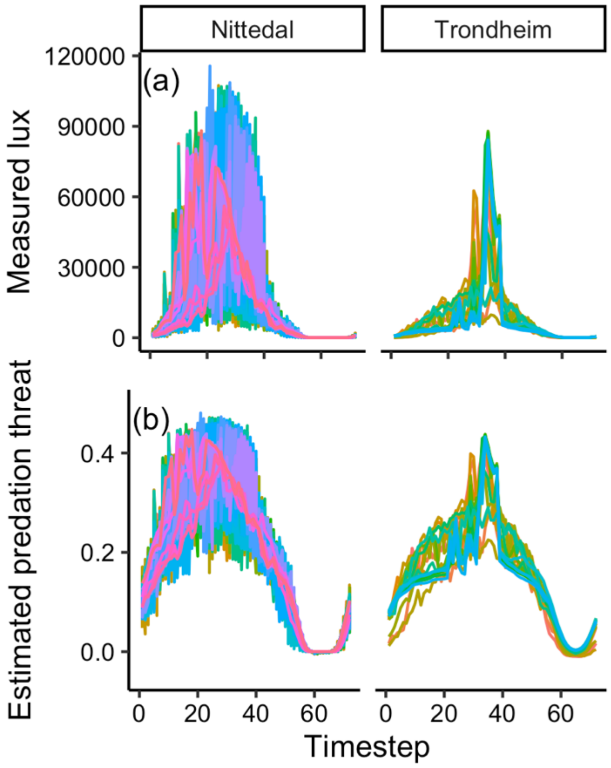


**Figure S1.5**: Daily cycles of light or light-dependent predation threats between sunrise and the following sunrise (one timestep corresponds to 20 minutes) for Nittedal and Trondheim in July. **(a)** The measured light levels (lux) at each location across the day and night. **(b)** Estimated predation threat at the two locations, converted from the measured light-levels as described in equation S1.6.

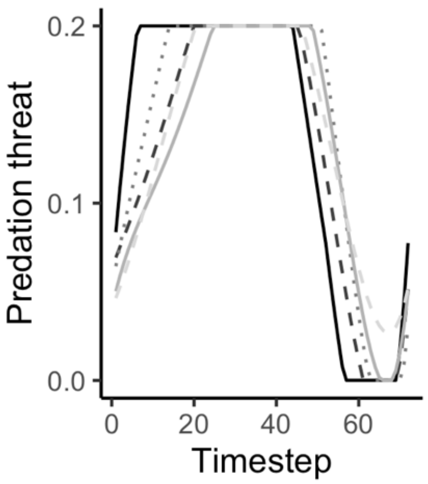


**Figure S1.6**: Estimated predation threat cycles for each of the five locations based upon light-measurements and sun altitudes, here shown with conditions of maximum values of 0.2 and minimum values of 0.0001. Cycles for each location is shown as listed: Nittedal as black solid line; Trondheim as dark grey dashed line; Bodø as grey dotted line; Tromsø as light grey solid line; Gamvik as light grey dashed line.

Nittedal: $0.06 + 0.03 \times t - 8.09 \times 10^{-4} \times t^2 + 1.29 \times 10^{-5} \times t^3 - 2.59 \times 10^{-7} \times t^4 + 2.26 \times 10^{-9} \times t^5$ if $t < 52$; $0.28 + 0.026 \times t - 8.83 \times 10^{-4} \times t^2 + 1.28 \times 10^{-5} \times t^3 - 2.56 \times 10^{-7} \times t^4 + 2.38 \times 10^{-9} \times t^5$ if $t \geq 52$ (S1.13)

Trondheim: $0.06 + 0.006 \times t - 4.95 \times 10^{-5} \times t^2 + 1.18 \times 10^{-5} \times t^3 - 4.35 \times 10^{-7} \times t^4 + 3.67 \times 10^{-9} \times t^5$ (S1.14)

Bodø $0.052 + 0.012 \times t - 3.3 \times 10^{-4} \times t^2 + 2.25 \times 10^{-5} \times t^3 - 6.36 \times 10^{-7} \times t^4 + 4.91 \times 10^{-9} \times t^5$ (S1.15)

Tromsø $0.040 + 0.011 \times t - 6.92 \times 10^{-4} \times t^2 + 3.89 \times 10^{-5} \times t^3 - 8.52 \times 10^{-7} \times t^4 + 5.79 \times 10^{-9} \times t^5$ (S1.16)

Gamvik $0.041 + 0.006 \times t + 2.13 \times 10^{-4} \times t^2 - 3.84 \times 10^{-6} \times t^3 - 1.06 \times 10^{-7} \times t^4 + 1.44 \times 10^{-9} \times t^5$ (S1.17)

1.2.5 Light-dependent energetic competition cost

Predation threat is not the single potential explanation of the evolved nocturnality of insectivorous bats, and three main alternative hypotheses have been presented and debated, these being overheating, competition from insectivorous birds and mobbing by non-competitors. Although strong support for these hypotheses have proven difficult to obtain through field-observations and lab-experiments (Speakman, 1991b; 1995; Speakman et al., 2000), there are likely complex relationships regarding daylight activity costs for insectivorous bats that does not solely involve the risk of being predated upon. We therefore added a second light-dependent cost to our model framework, this being an energetic cost rather than a direct fitness cost, to account for potential direct or indirect energy expenditure caused by interspecific competition or mobbing by birds. We used the measured or estimated daylight cycles as described in section 1.2.4 above and converted the values as described in equation S1.18 (below) to generate what we considered to be realistic value-ranges (see Supplementary Materials 2), although with an implemented higher threshold of 0.2 due to assumptions of the energetic competition cost becoming light-independent above certain values (Fig. S1.7).

Energetic competition cost: $= \frac{(\log(\text{lux}))^{1.8}}{70}$ . (S1.18)

Fifth degree polynomial model equations describing the temporal energetic competition cost for each location are defined with equations S1.19 to S1.23 below.

| | | |
|---|---|---|
| Nittedal: | $0.11\ g + 0.02 \times t - 0.001 \times t^2 + 4.72 \times 10^{-5} \times t^3 - 8.34 \times 10^{-7} \times t^4 + 5.31 \times 10^{-9} \times t^5$ | (S1.19) |
| Trondheim: | $0.09\ g + 0.01 \times t - 7.48 \times 10^{-4} \times t^2 + 3.24 \times 10^{-5} \times t^3 - 6.29 \times 10^{-7} \times t^4 + 4.10 \times 10^{-9} \times t^5$ | (S1.20) |
| Bodø | $0.11 + 0.012 \times t - 7.35 \times 10^{-4} \times t^2 + 3.13 \times 10^{-5} \times t^3 - 6.13 \times 10^{-7} \times t^4 + 3.98 \times 10^{-9} \times t^5$ | (S1.21) |
| Tromsø | $0.10 + 0.01 \times t - 7.12 \times 10^{-4} \times t^2 + 3.06 \times 10^{-5} \times t^3 - 5.73 \times 10^{-7} \times t^4 + 3.59 \times 10^{-9} \times t^5$ | (S1.22) |
| Gamvik | $0.11 + 0.006 \times t - 1.99 \times 10^{-4} \times t^2 + 8.12 \times 10^{-6} \times t^3 - 1.92 \times 10^{-7} \times t^4 + 1.4 \times 10^{-9} \times t^5$ | (S1.23) |

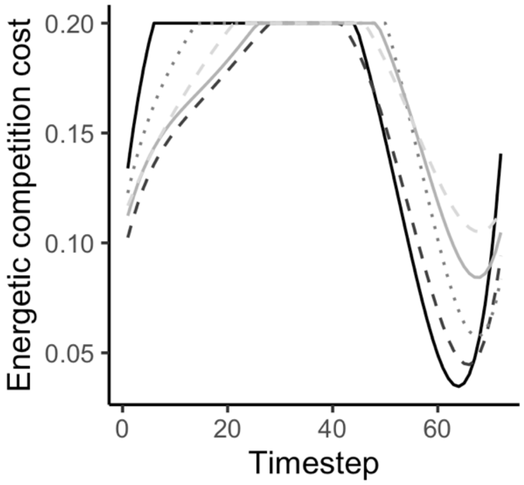


**Figure S1.7**: Estimated competition cost cycles for each of the five locations based upon light-measurements and sun altitudes, here shown with conditions of maximum values of 0.2. Cycles for each location is shown as listed: Nittedal as black solid line; Trondheim as dark grey dashed line; Bodø as grey dotted line; Tromsø as light grey solid line; Gamvik as light grey dashed line.

### Supplementary Materials 2: Linear interpolation

Because the energetic state variable ($x$) used for calculating fitness values is a discrete rather than a continuous variable, the model needs to specify how the model calculates corresponding fitness values from energetic values that fall in-between discrete steps. For this framework we have implemented the interpolation method described in Clark and Mangel (2000). Here, the model assumes that there is a linear increase in the fitness values from one discrete energetic reserve value to the next. For a current energetic reserve value ($x$) that does not correspond to one of the specified discrete integers ($j$), we can specify the following linear interpolation functions (as described in Clark & Mangel, 2000):

Fit($x_c$, y, w, t, d) = V(x, y, w, t, d) . (2.1)

If $j \leq x_c < j + 1$ , (2.2)

then $\Delta x = x - j$ , (2.3)

then V(x, y, w, t, d) = Fit($x_c$, y, w, t, d) ≅

$$\begin{cases} (1 - \Delta x_c) Fit(j, y, w, t, d) + \Delta x_c Fit(j + 1, y, w, t, d) & \text{if } j < j_{max} \\ Fit(j_{max}, y, w, t, d) & \text{if } j = j_{max} \end{cases} . \quad (2.4)$$

### Supplementary Materials 3: Parameter adjustments

Some parameter values were not possible to obtain or verify from the literature, because they lack any previous quantification. This included the parameters for predation threat, energetic competition cost and resting benefit. However, we judged these parameters to be important for our model system based upon assumptions regarding drivers of general bat behaviour (see Discussion in main text), and therefore felt we needed to attempt to include more-or-less realistic values followed by sensitivity tests where we tested the consequences in our model system if the values were increased or decreased (Fig. S3.1-11). Both the predation threat and the competition costs were based on light-measurements (see Supplementary Materials 1), and thus follow the established daily cycle but further testing was required to arrive at the most reasonable and hopefully realistic values. The resting fitness benefit was assumed to be time and temperature independent and thus kept as a simple non-static value.

Figures S3.1-S3.11 show the summary plots of daily decisions from 'forward' simulations of 200 individuals across 30 days for Nittedal and Trondheim, revealing the results of different parameter values (within figures) across day types (between figures). We carried out evaluations of each of the three parameters as described below.

Figures S3.1-S3.11 show that the removal of the benefit of resting over torpor ($\theta = 0$) resulted in no resting behaviour being expressed across any day type or location. Because bats have been found capable of using heat generated from initiating flight to finalise their rewarming process from torpor (Willis & Brigham, 2003), bats in our model were not required to enter a resting state between being torpid and going out to forage, and could thus go directly from being torpid to foraging, although they still had to pay the temperature-dependent rewarming cost when carrying out this transition. With no benefits of being awake, resting behaviour was thus never expressed in our model even on warm days. An enlarged resting benefit ($\theta = 0.003$), however, led to excessive amounts of time spent resting during day and night on any of the warm day types (Fig. S3.1-3 and S3.6-9). Given the other known baseline values (Table 1 in the Methods main text), a resting benefit of 0.0015 was thus considered most representative. This is because patterns where the bats spend most of the day awake and resting are usually associated to unusual and excessively warm days (corresponding to 'very warm' day types in our model framework) in non-reproductive individuals (see Turbill et al., 2003).

Alterations of either side of baseline levels of predation threat also led to less realistic outcomes. A decrease in predation threat ($\mu$ - 0.03) resulted in bats leaving the roost to forage long before sunset, while an increase ($\mu$ + 0.03) resulted in delaying the onset of foraging until long after sunset while experiencing high mortality from predation, especially in the Trondheim scenario.

Adjusting the energetic competition cost (assuming this is caused by interspecific competition from diurnal insectivorous birds or energy expenditure from avoiding mobbing by crows or other birds) led to alterations in the amount of time spent resting and foraging on days and nights across day types.

With no competition cost implemented in the model (*C* = 0), the bats spent excessive amounts of time resting and foraging, even on some of the colder day types. An increase in the energetic competition cost (*C* + 0.05), on the other hand, led to a drastic reduction in time spent foraging and resting on any of the warm day types at either location. Even on 'very warm' days did the bats spend a considerable part of the morning torpid with this adjusted scenario.

Like any other parameter from the baseline scenarios (Table 1 in Methods), the values used in our model may not be representative for any specific bat population at any particular habitat or latitude, but as a general model framework for high-latitude living bats we believe these previously non-quantified relative values are very likely to be within realistic ranges.

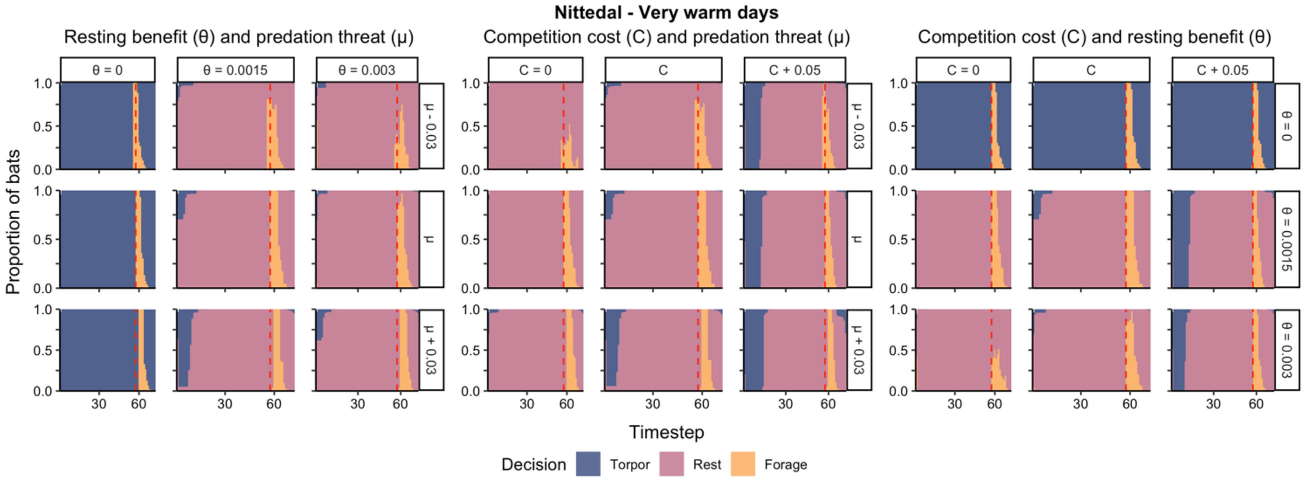


**Fig. S3.1**: Summary plots of all individual activity decisions across the daily cycle (from sunrise to following sunrise) for 'very warm' days in Nittedal from the forward iterations. Each pane represents one summary plot for given values of resting benefits (*θ*), predation threats (*μ*) and competition costs (*C*). The three parameters are tested pairwise, with the third parameter kept at baseline levels. The plots illustrate how an increase (right and bottom panes) or a decrease (left and top panes) in these parameter values affect daily activity patterns in bats. Red dashed vertical lines indicate the timing of sunset.

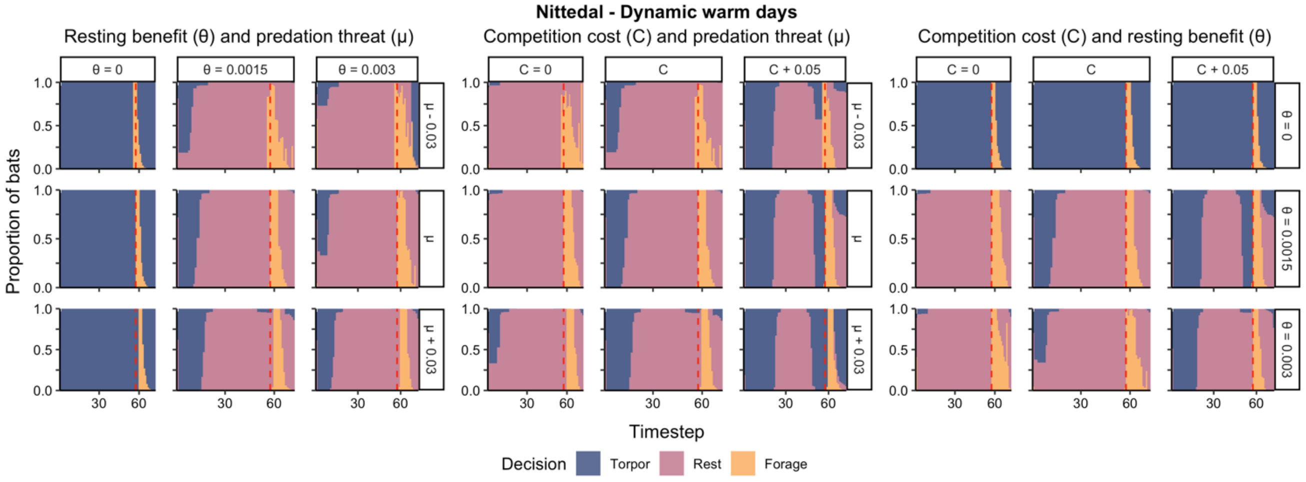


**Fig. S3.2**: Summary plots of all individual activity decisions across the daily cycle for 'dynamic warm' days in Nittedal from the forward iterations.

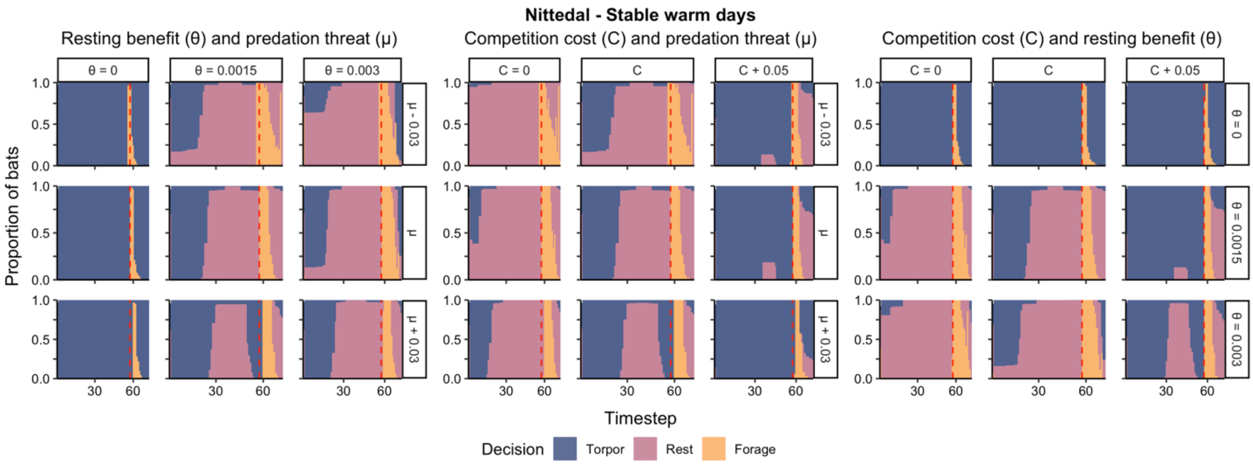


**Fig. S3.3**: Summary plots of all individual activity decisions across the daily cycle for 'stable warm' days in Nittedal from the forward iterations.

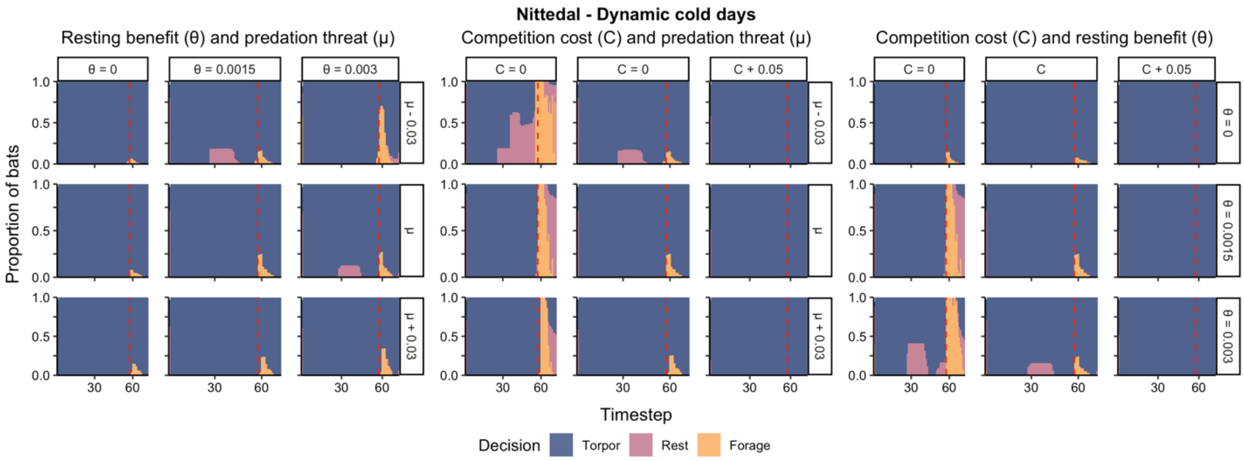


**Fig. S3.4**: Summary plots of all individual activity decisions across the daily cycle for 'dynamic cold' days in Nittedal from the forward iterations.

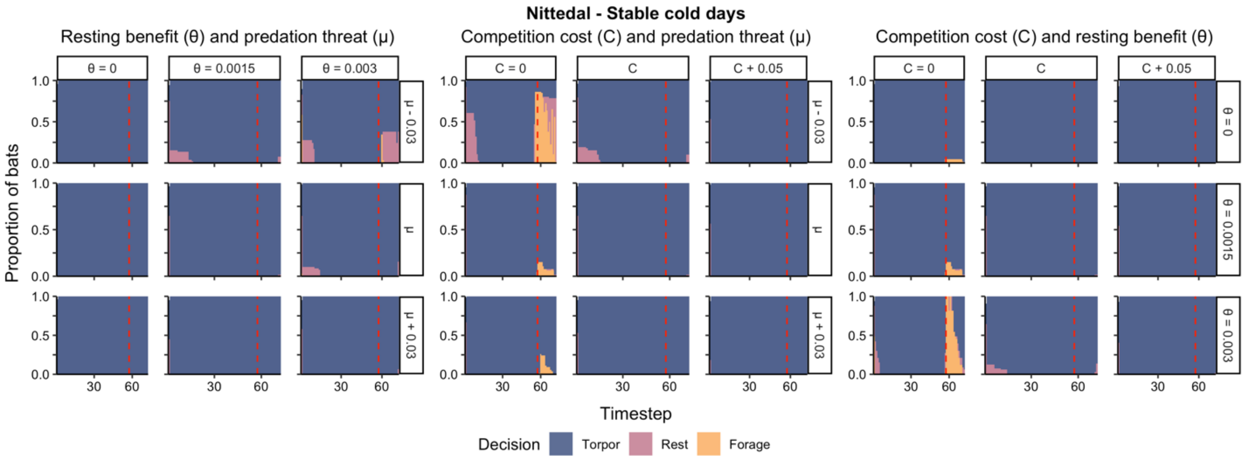


**Fig. S3.5**: Summary plots of all individual activity decisions across the daily cycle for 'stable cold' days in Nittedal from the forward iterations.

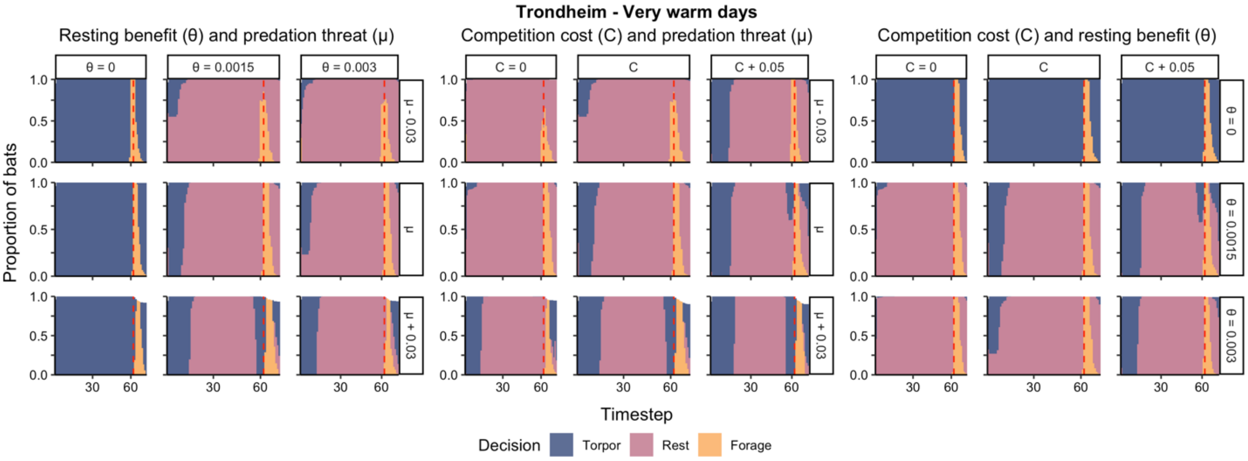


**Fig. S3.6**: Summary plots of all individual activity decisions across the daily cycle for ‘very warm’ days in Trondheim from the forward iterations.

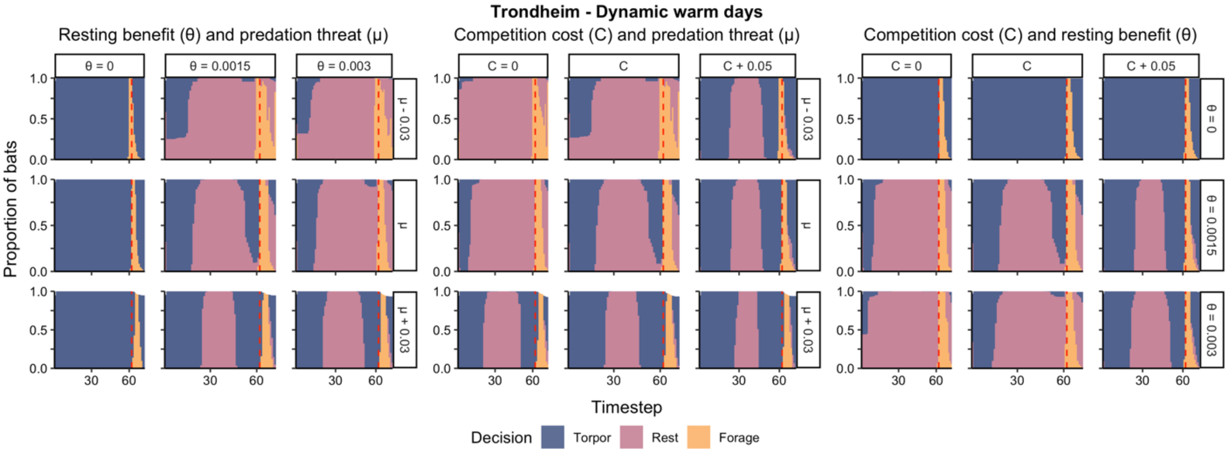


**Fig. S3.7**: Summary plots of all individual activity decisions across the daily cycle for ‘dynamic warm’ days in Trondheim from the forward iterations.

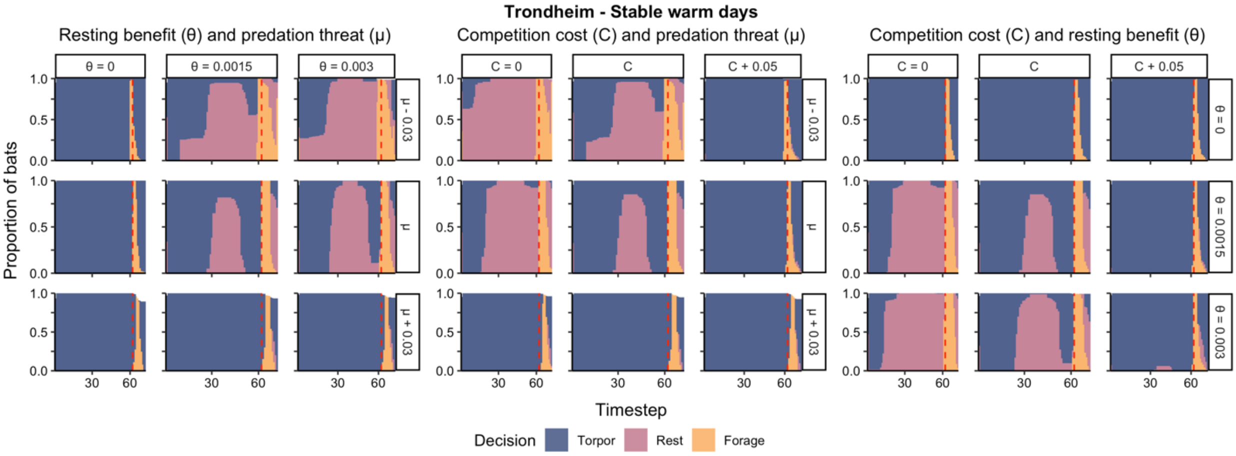


**Fig. S3.8**: Summary plots of all individual activity decisions across the daily cycle for ‘stable warm’ days in Trondheim from the forward iterations.

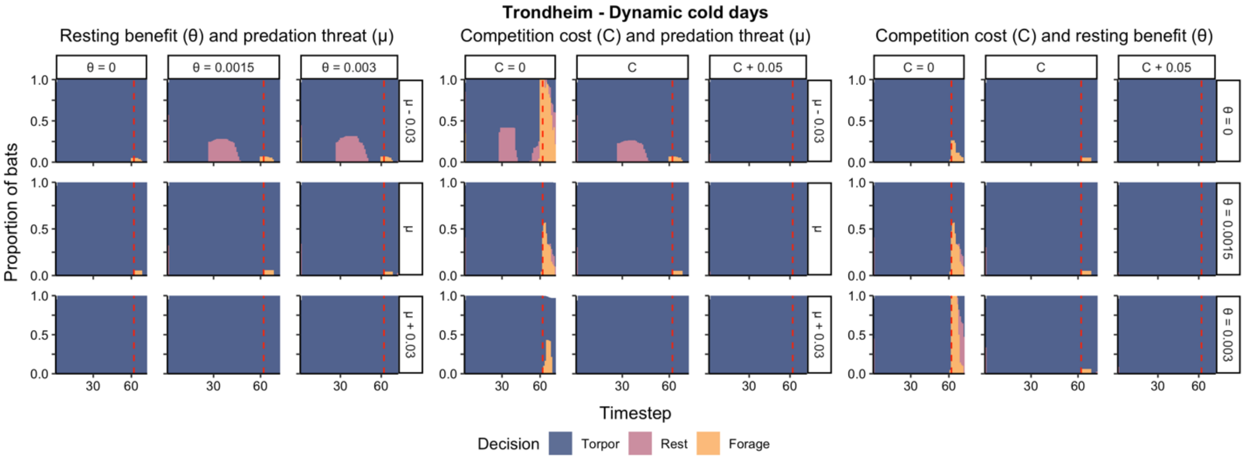


**Fig. S3.9**: Summary plots of all individual activity decisions across the daily cycle for ‘dynamic cold’ days in Trondheim from the forward iterations.

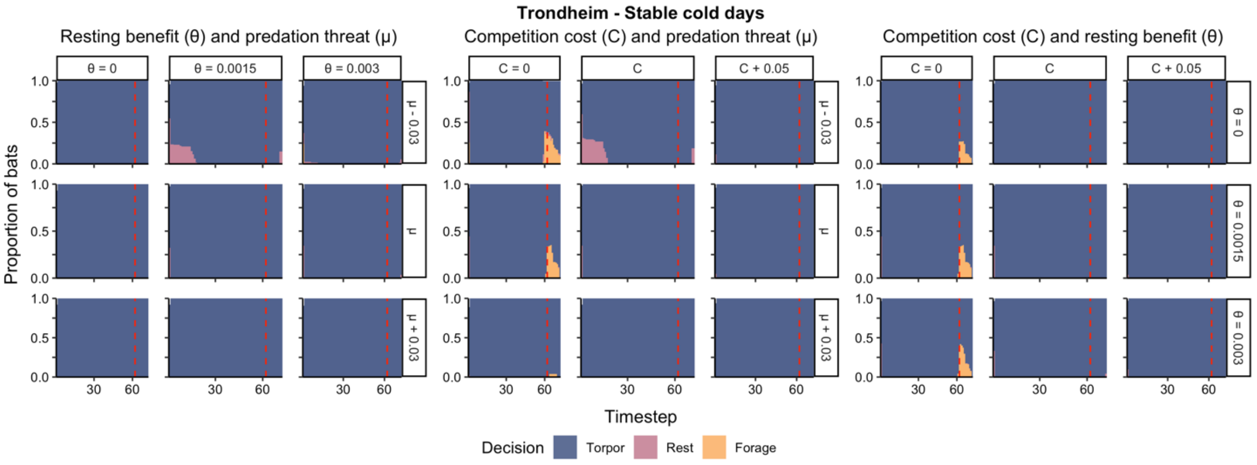


**Fig. S3.10**: Summary plots of all individual activity decisions across the daily cycle for ‘stable cold’ days in Trondheim from the forward iterations.

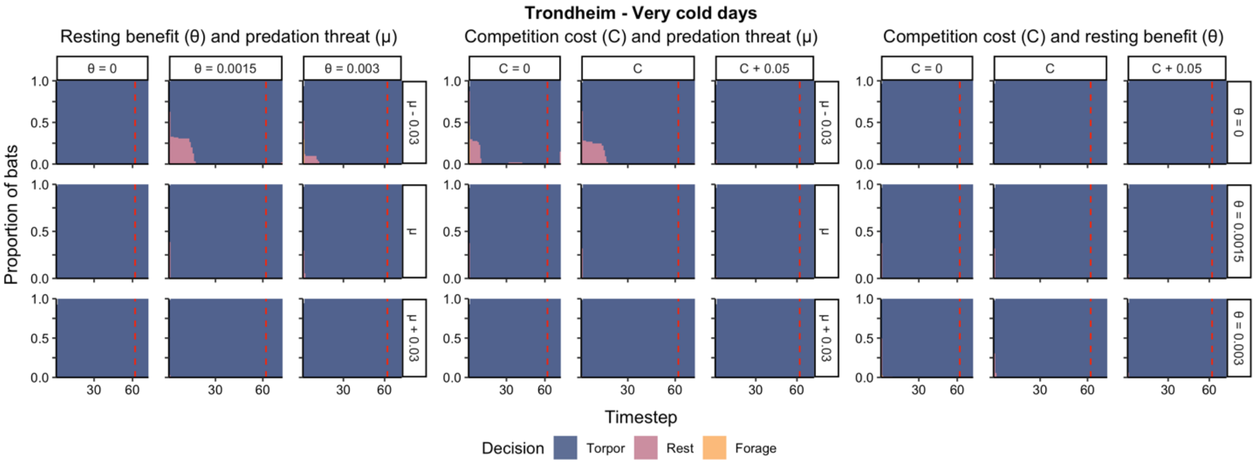


**Fig. S3.11**: Summary plots of all individual activity decisions across the daily cycle for ‘very cold’ days in Trondheim from the forward iterations.

### Supplementary Materials 4: Fat reserves across the daily cycle

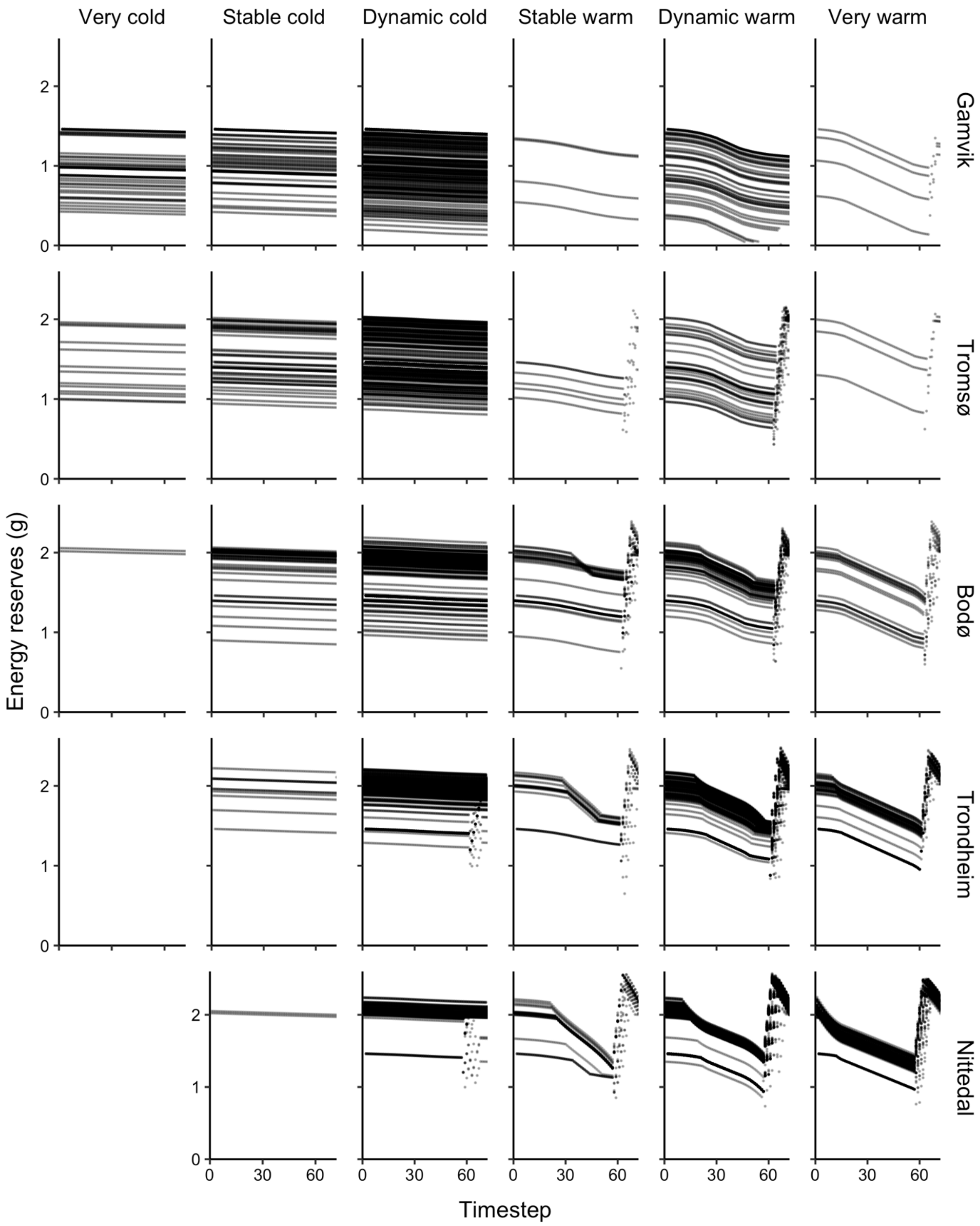


**Figure S4.1**: Summary plots of all individual fat reserves across the daily cycle (from sunrise to following sunrise) extracted from the forward iterations with baseline values across day types in the Nittedal, Trondheim, Bodø, Tromsø and Gamvik scenarios. Given the different probabilities for each day type at each location, the number of individual trajectories vary between day types and locations (the day type 'very cold' days had an occurrence probability = 0 in the Nittedal scenario and is thus not shown in the figure).

### Supplementary Materials 5: Empirical data comparisons

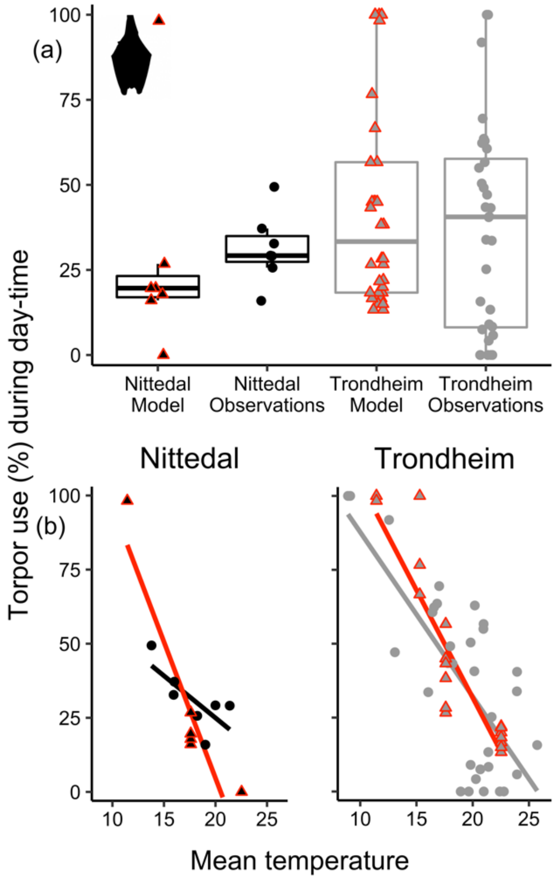


**Figure S5.1**: Comparisons of day-time torpor use (%) from field derived data (round points) and from model simulations (triangle points with red outline) for the two locations of Nittedal (black points and boxes) and Trondheim (grey points and boxes). **(a)** Boxplots of daily torpor use between locations and data origin with boxes marking the data median and the 25th and 75th percentiles. **(b)** Negative effects of increasing mean daily temperatures on day-time torpor expression, both in the empirical data (round points and black or grey lines) and in data generated from matched simulated model runs (triangle points and red lines).

**Table S5.1**: Means and ranges of the percentages of day-time and night-time behaviour in the empirical data and from the model simulations at the two locations Nittedal and Trondheim.

| | Nittedal | | Trondheim | |
|---|---|---|---|---|
| | Empirical data | Model simulations | Empirical data | Model simulations |
| *Daily torpor use* | | | | |
| Mean time (± sd) | 31.3% (±10.3) | 29.1% (±31.6) | 37.3% (±30.3) | 44.0% (±29.7) |
| Range time | 15.9% to 49.4% | 0% to 98.2% | 0% to 100% | 13.3% to 100% |
| *Nightly foraging* | | | | |
| Mean time (± sd) | 60.3% (±14.1) | 33.0% (±20.0) | 39.3% (±30.0) | 42.2% (±23.7) |
| Range time | 32.3% to 74.2% | 0% to 62.5% | 0% to 80.8% | 0% to 66.7% |
| *Nightly torpor use* | | | | |
| Mean time (± sd) | 20.8% (±11.4) | 25.9% (±44.6) | 34.7% (±39.2) | 29.6% (±36.3) |
| Range time | 0% to 32.3% | 0% to 100% | 0% to 100% | 0% to 100% |
| *Nightly resting* | | | | |
| Mean time (± sd) | 18.9% (±10.8) | 41.1% (±29.9) | 24.0% (±18.8) | 28.2% (25.2%) |
| Range time | 0% to 35.5% | 0% to 68.8% | 0% to 58.3% | 0% to 66.7% |

**References**


Anthony, E. L., & Kunz, T. H. (1977). Feeding strategies of the little brown bat, Myotis lucifugus, in southern New Hampshire. *Ecology, 58*(4), 775-786.

Clark, C. W., & Mangel, M. (2000). *Dynamic state variable models in ecology: methods and applications*. Oxford: Oxford University Press.

Cruz-Neto, A., & Jones, K. (2006). Exploring the evolution of basal metabolic rate in bats. In *Functional morphology and ecology of bats* (pp. 58-69). New York: Oxford University Press.

Currie, S. E., Noy, K., & Geiser, F. (2015). Passive rewarming from torpor in hibernating bats: minimizing metabolic costs and cardiac demands. *American Journal of Physiology-Regulatory, Integrative and Comparative Physiology, 308*(1), 34-41. doi:https://doi.org/10.1152/ajpregu.00341.2014

Fjelldal, M. A., Sørås, R., & Stawski, C. (2022). Universality of Torpor Expression in Bats. *Physiological and Biochemical Zoology, 95*(4), 326-339.

Geiser, F., & Brigham, R. M. (2000). Torpor, thermal biology, and energetics in Australian long-eared bats (*Nyctophilus*). *Journal of Comparative Physiology B, 170*(2), 153-162. doi:https://doi.org/10.1007/s003600050270

Jones, G., & Rydell, J. (2005). Attack and defense: interactions between echolocating bats and their insect prey. In T. H. Kunz & M. B. Fenton (Eds.), *Bat Ecology* (pp. 301-345). Chicago: Chicago University Press.

Kerth, G., Weissmann, K., & König, B. (2001). Day roost selection in female Bechstein's bats (Myotis bechsteinii): a field experiment to determine the influence of roost temperature. *Oecologia, 126*(1), 1-9.

Kunz, T. H. (1980). *Daily energy budgets of free-living bats.* Paper presented at the Proceedings of the fifth international bat research conference (DE Wilson and AL Gardner, eds.). Texas Tech Press, Lubbock.

Kunz, T. H. (1988). Methods of assessing the availability of prey to insectivorous bats. *Ecological and behavioral methods for the study of bats*, 191-210.

Kurta, A., Bell, G. P., Nagy, K. A., & Kunz, T. H. (1989). Energetics of pregnancy and lactation in freeranging little brown bats (Myotis lucifugus). *Physiological Zoology, 62*(3), 804-818.

Lausen, C. L., & Barclay, R. M. (2003). Thermoregulation and roost selection by reproductive female big brown bats (*Eptesicus fuscus*) roosting in rock crevices. *Journal of Zoology, 260*(3), 235-244.

Lima, S. L., & O'Keefe, J. M. (2013). Do predators influence the behaviour of bats? *Biological Reviews, 88*(3), 626-644. doi:10.1111/brv.12021

Lourenço, S. I., & Palmeirim, J. M. (2004). Influence of temperature in roost selection by *Pipistrellus pygmaeus* (Chiroptera): relevance for the design of bat boxes. *Biological Conservation, 119*(2), 237-243.

Michaelsen, T. C., Jensen, K. H., & Högstedt, G. (2014). Roost site selection in pregnant and lactating soprano pipistrelles (Pipistrellus pygmaeus Leach, 1825) at the species northern extreme: the importance of warm and safe roosts. *Acta Chiropterologica, 16*(2), 349-357.

Reher, S., & Dausmann, K. H. (2021). Tropical bats counter heat by combining torpor with adaptive hyperthermia. *Proceedings of the Royal Society B: Biological Sciences, 288*(1942), 20202059. doi:https://doi.org/10.1098/rspb.2020.2059

Skåra, K. H., Bech, C., Fjelldal, M. A., van Der Kooij, J., Sørås, R., & Stawski, C. (2021). Energetics of whiskered bats in comparison to other bats of the family Vespertilionidae. *Biology Open, 10*(8), bio058640.

Sørås, R., Fjelldal, M. A., Bech, C., van der Kooij, J., Skåra, K. H., Eldegard, K., & Stawski, C. (2022). State dependence of arousal from torpor in brown long-eared bats (Plecotus auritus). *Journal of Comparative Physiology B*, 1-13. doi:10.1007/s00360-022-01451-8

Speakman, J. (1991a). The impact of predation by birds on bat populations in the British Isles. *Mammal Review, 21*(3), 123-142.

Speakman, J. (1991b). Why do insectivorous bats in Britain not fly in daylight more frequently? *Functional Ecology*, 518-524.

Speakman, J. (1995). *Chiropteran nocturnality.* Paper presented at the Symposia of the zoological society of London.

Speakman, J., Rydell, J., Webb, P., Hayes, J., Hays, G., Hulbert, I., & McDevitt, R. (2000). Activity patterns of insectivorous bats and birds in northern Scandinavia (69 N), during continuous midsummer daylight. *Oikos, 88*(1), 75-86.

Speakman, J., & Thomas, M. D. (2003). Physiological ecology and energetics of bats. In T. H. Kunz & M. B. Fenton (Eds.), *Bat Biology* (pp. 430–492). Chicago: University of Chicago Press.

Stawski, C. (2012). Capture and care of northern long-eared bats (*Nyctophilus bifax*) and seasonal changes in insect abundance. *Australian mammalogy, 34*(2), 245-250. doi:10.1071/AM11043

Stawski, C., Turbill, C., & Geiser, F. (2008). *Prolonged torpor use during winter by a free-ranging bat in subtropical Australia.* Paper presented at the Hypometabolism in animals: Hibernation, torpor and cryobiology. 13th International Hibernation Symposium. University of KwaZulu-Natal, Pietermaritzburg, South Africa.

Thomas, S. P. (1975). Metabolism during flight in two species of bats, Phyllostomus hastatus and Pteropus gouldii. *Journal of Experimental Biology, 63*(1), 273-293.

Turbill, C., Kortner, G., & Geiser, F. (2008). Timing of the daily temperature cycle affects the critical arousal temperature and energy expenditure of lesser long-eared bats. *Journal of Experimental Biology, 211*(Pt 24), 3871-3878. doi:10.1242/jeb.023101

Turbill, C., Körtner, G., & Geiser, F. (2003). Natural use of heterothermy by a small, tree-roosting bat during summer. *Physiological and Biochemical Zoology, 76*(6), 868-876.

Willis, C. K., & Brigham, R. (2003). Defining torpor in free-ranging bats: experimental evaluation of external temperature-sensitive radiotransmitters and the concept of active temperature. *Journal of Comparative Physiology B, 173*(5), 379-389. doi:https://doi.org/10.1007/s00360-003-0343-y

### R-codes for stochastic dynamic programming model

###### SCRIPT WITH PREPARATIONS FOR BACKWARDS ITERATION ####

###### Parameter values (CAN BE CHANGED) ####

###### State (fat reserves) ####
mass_zero_fat <- 7 # Mass of bat with zero fat reserves (g)
fat_max <- 3 # Maximum fat reserves (g)
fat_discretized <- seq(from = 0, to = fat_max, length.out=40) # Discretized values of fat_state
predation_risk_increase <- 0.05 # Increase in predation risk due to body mass (/g fat reserves)

###### State (torpor-use) ####
temp_state <- c(1,2) # state 1 = torpid, state 2 = not torpid

###### Fitness (time, probability of good or bad night) ####

### A season is divided in days, themselves divided in timesteps.
nb_timesteps <- 72 # Number of time periods per day
nb_days <- 30 # Number of days in summer month
nb_hours <- 24 # Number of hours in a day, used later to calculate metabolic cost per timestep instead of per hour.

### Environmental stochasticity and foraging
prob_finding_food <- 0.9 # Stochasticity in foraging success
foraging_success_decline <- -0.03 # Decrease in foraging success with increasing fat reserves (fat bats are less agile)
weather <- c(1:6) # weather on that day (very warm (1), dynamic warm (2), stable warm (3), dynamic cold (4), stable cold (5) or very cold (6)); affects the output of the get_temperature_XXX functions

###### Metabolism and physiology ####

tnz <- 29 # thermoneutral zone (TNZ) in degrees celsius, used to define when to apply resting metabolic rate (RMR) or basal metabolic rate (BMR)
bmr <- 0.0042*mass_zero_fat # BMR, used to calculate metabolic cost of resting in the TNZ (using results from Geiser and Turbill 2000)
cost_flight_hourly <- 0.615 # Hourly cost of flight in g for a little bat
energy_gain <- 1.3
fat_survival_threshold <- 0.05 # If bats reach fat reserve levels below this threshold they die (fitness = 0)
fat_survival_decrease <- 2 # Fat level threshold for a linear decrease in fitness until the fat_survival_threshold is reached and fitness = 0

###### Scenarios ####
Temp_scenario <- "Nittedal" #Can be one of the following 5: Trondheim, Nittedal, Bodø, Tromsø or Gamvik (affects the temperature cycles and the prob of good days)

```
Light_scenario <- "Nittedal" #Can be one of the following 5: Trondheim, Nittedal, Bodø,
Tromsø or Gamvik (affects the predation threat cycles)

weather_prob <- if (Temp_scenario == "Nittedal") {
  weather_prob <- c(0.238, 0.476, 0.0741, 0.201, 0.0106, 0)
}  else if (Temp_scenario == "Trondheim")  {
  weather_prob <- c(0.13, 0.30, 0.0435, 0.47, 0.0395, 0.0158)
}  else if (Temp_scenario == "Bodø")  {
  weather_prob <- c(0.0612, 0.219, 0.0791, 0.507, 0.126, 0.00719)
}  else if (Temp_scenario == "Tromsø")  {
  weather_prob <- c(0.015, 0.135, 0.0262, 0.648, 0.131, 0.0449)
}  else if (Temp_scenario == "Gamvik")  {
  weather_prob <- c(0.0223, 0.108, 0.0149, 0.532, 0.156, 0.167)
}

# Predation threath scenarios (affects the intercept of the predation threath daily cycle)
pred_low <- -0.03
pred_base <- 0
pred_high <- 0.03

# Energetic competition cost scenarios (affects the intercept of the competition cost)
comp_cost_low <- -0.05
comp_cost_base <- 0
comp_cost_high <- 0.05

# Resting benefit (theta) scenarios (a per-timestep fitness benefit when choosing to rest)
theta_low <- 0
theta_base <- 0.0015
theta_high <- 0.003

# Temperature scenario (for the last part of the results)
Ta_low <- -2
Ta_base <- 0
Ta_high <- 2

# Choose the scenario to run

pred_scenario <- pred_base
resting_fitness_benefit <- theta_base
comp_cost_scenario <- comp_cost_base
Ta_scenario <- Ta_base

#External temperature on different day types in degrees celsius (affects prey availability)

get_temperature_ext_very_warm <- function(time_current){
  temperature_ext_current <- Ta_scenario + 1.658e+01 + (1.176e-01*time_current) +
(2.438e-02*time_current^2) - (8.987e-05*time_current^3) - (2.954e-05*time_current^4) +
(6.013e-07*time_current^5) - (3.375e-09*time_current^6)
  return(temperature_ext_current)
}
```

```
get_temperature_ext_dynamic_warm <- function(time_current){
  temperature_ext_current <- Ta_scenario + 1.411e+01 + (1.147e-01*time_current) +
(9.198e-03*time_current^2) + (2.528e-04*time_current^3) - (2.555e-05*time_current^4) +
(4.470e-07*time_current^5) - (2.359e-09*time_current^6)
  return(temperature_ext_current)
}


get_temperature_ext_stable_warm <- function(time_current){
  temperature_ext_current <- Ta_scenario + 1.395e+01 + (1.082e-01*time_current) - (6.717e-
03*time_current^2) + (5.616e-04*time_current^3) - (1.815e-05*time_current^4) + (2.299e-
07*time_current^5) - (1.002e-09*time_current^6)
  return(temperature_ext_current)
}


get_temperature_ext_dynamic_cold <- function(time_current){
  temperature_ext_current <- Ta_scenario + 9.622e+00 + (3.908e-02*time_current) +
(7.858e-03*time_current^2) - (8.429e-05*time_current^3) - (6.033e-06*time_current^4) +
(1.238e-07*time_current^5) - (6.431e-10*time_current^6)
  return(temperature_ext_current)
}


get_temperature_ext_stable_cold <- function(time_current){
  temperature_ext_current <- Ta_scenario + 9.390e+00 + (3.525e-02*time_current) - (4.099e-
03*time_current^2) + (2.890e-04*time_current^3) - (8.411e-06*time_current^4) + (1.025e-
07*time_current^5) - (4.431e-10*time_current^6)
  return(temperature_ext_current)
}


get_temperature_ext_very_cold <- function(time_current){
  temperature_ext_current <- Ta_scenario + 6.305e+00 + (4.960e-03*time_current) +
(1.621e-03*time_current^2) + (1.799e-04*time_current^3) - (9.024e-06*time_current^4) +
(1.267e-07*time_current^5) - (5.682e-10*time_current^6)
  return(temperature_ext_current)
}




# Roost temperature for the different types of days in degrees celsius (affects metabolism
when resting or using torpor)
```

```
# Warm days follow the same dynamic pattern in relation to air temperature, while cold days all are 2 degrees warmer than the air temperature

get_temperature_roost_very_warm <- function(time_current){
  temperature_roost_current <- Ta_scenario + 1.658e+01 + (1.176e-01*time_current) +
(2.438e-02*time_current^2) - (8.987e-05*time_current^3) - (2.954e-05*time_current^4) +
(6.013e-07*time_current^5) - (3.375e-09*time_current^6) +
    (2.342e+00 + (2.708e-01*time_current) - (2.754e-04*time_current^2) - (4.875e-
05*time_current^3))
  return(temperature_roost_current)
}

get_temperature_roost_dynamic_warm <- function(time_current){
  temperature_roost_current <- Ta_scenario + 1.411e+01 + (1.147e-01*time_current) +
(9.198e-03*time_current^2) + (2.528e-04*time_current^3) - (2.555e-05*time_current^4) +
(4.470e-07*time_current^5) - (2.359e-09*time_current^6) +
    (2.342e+00 + (2.708e-01*time_current) - (2.754e-04*time_current^2) - (4.875e-
05*time_current^3))
  return(temperature_roost_current)
}

get_temperature_roost_stable_warm <- function(time_current){
  temperature_roost_current <- Ta_scenario + 1.395e+01 + (1.082e-01*time_current) -
(6.717e-03*time_current^2) + (5.616e-04*time_current^3) - (1.815e-05*time_current^4) +
(2.299e-07*time_current^5) - (1.002e-09*time_current^6) +
    (2.342e+00 + (2.708e-01*time_current) - (2.754e-04*time_current^2) - (4.875e-
05*time_current^3))
  return(temperature_roost_current)
}


get_temperature_roost_dynamic_cold <- function(time_current){
  temperature_roost_current <- Ta_scenario + 9.622e+00 + (3.908e-02*time_current) +
(7.858e-03*time_current^2) - (8.429e-05*time_current^3) - (6.033e-06*time_current^4) +
(1.238e-07*time_current^5) - (6.431e-10*time_current^6) + 2
  return(temperature_roost_current)
}

get_temperature_roost_stable_cold <- function(time_current){
  temperature_roost_current <- Ta_scenario + 9.390e+00 + (3.525e-02*time_current) -
(4.099e-03*time_current^2) + (2.890e-04*time_current^3) - (8.411e-06*time_current^4) +
(1.025e-07*time_current^5) - (4.431e-10*time_current^6) + 2
  return(temperature_roost_current)
}

get_temperature_roost_very_cold <- function(time_current){
  temperature_roost_current <- Ta_scenario + 6.305e+00 + (4.960e-03*time_current) +
(1.621e-03*time_current^2) + (1.799e-04*time_current^3) - (9.024e-06*time_current^4) +
(1.267e-07*time_current^5) - (5.682e-10*time_current^6) + 2
  return(temperature_roost_current)
```

```
}





# Prey availability (depends on current outside temperature)
get_prey <- function(temperature_ext_current){
  reward_prey_current <- 1 / (1 + exp( -0.524* (temperature_ext_current - 8)))
  return(reward_prey_current)
}


# Predation threat cycle (no difference between types of days)

risk_predation_baseline <- function(time_current){
  if (Light_scenario == "Trondheim") {
    pred_current <- pred_scenario + 0.06333 + (0.006028*time_current) -
(0.00004952*time_current^2) + (0.00001183*time_current^3) -
(0.0000004353*time_current^4) + (0.000000003666*time_current^5)
  }  else if (Light_scenario == "Nittedal") {
    pred_current <- ifelse(time_current < 52, pred_scenario + 0.057 + (0.02739*time_current)
- (0.0008094*time_current^2) + (0.00001293*time_current^3) -
(0.0000002586*time_current^4) + (0.000000002255*time_current^5),
                           pred_scenario + 0.28 + 0.026*time_current - 0.000883*time_current^2 +
0.0000128*time_current^3 - 0.000000256*time_current^4 +
0.00000000238*time_current^5)
  }  else if (Light_scenario == "Bodø") {
    pred_current <- pred_scenario + 0.05231 + 0.01239*time_current -
0.0003302*time_current^2 + 0.00002246*time_current^3 - 0.0000006359*time_current^4 +
0.000000004912*time_current^5
  }  else if (Light_scenario == "Tromsø") {
    pred_current <- pred_scenario + 0.04045 + 0.0105*time_current -
0.0006918*time_current^2 + 0.00003891*time_current^3 - 0.0000008519*time_current^4 +
0.000000005794*time_current^5
  }  else if (Light_scenario == "Gamvik") {
    pred_current <- pred_scenario + 0.04051 + 0.005727*time_current +
0.0002128*time_current^2 - 0.000003837*time_current^3 - 0.0000001062*time_current^4 +
0.000000001438*time_current^5
  }

  return(pred_current)
}

risk_predation_baseline <- risk_predation_baseline(1:nb_timesteps)
```

```
risk_predation_baseline<-
ifelse(risk_predation_baseline[1:nb_timesteps]<0.0001,0.0001,risk_predation_baseline[1:nb_
timesteps])
risk_predation_baseline<-
ifelse(risk_predation_baseline[1:nb_timesteps]>0.2,0.2,risk_predation_baseline[1:nb_timeste
ps])


get_predation <- function(time_current){
  predation_current <- risk_predation_baseline[time_current]
  return(predation_current)
}



# Light-dependent energetic foraging success cost (caused by interspecific competition)

competition_cost_baseline <- function(time_current){
  if (Light_scenario == "Trondheim") {
    comp_cost_current <- comp_cost_scenario + 0.09249 + 1.067e-02*time_current - 7.479e-
04*time_current^2 + 3.239e-05*time_current^3 - 6.291e-07*time_current^4 + 4.097e-
09*time_current^5
  }  else if (Light_scenario == "Nittedal") {
    comp_cost_current <- comp_cost_scenario + 0.11426 + 2.097e-02*time_current - 1.348e-
03*time_current^2 + 4.724e-05*time_current^3 - 8.336e-07*time_current^4 + 5.310e-
09*time_current^5
  }  else if (Light_scenario == "Bodø") {
    comp_cost_current <- comp_cost_scenario + 0.11145 + 1.199e-02*time_current - 7.351e-
04*time_current^2 + 3.132e-05*time_current^3 - 6.127e-07*time_current^4 + 3.977e-
09*time_current^5
  }  else if (Light_scenario == "Tromsø") {
    comp_cost_current <- comp_cost_scenario + 0.10306 + 1.000e-02*time_current - 7.121e-
04*time_current^2 + 3.063e-05*time_current^3 - 5.729e-07*time_current^4 + 3.585e-
09*time_current^5
  }  else if (Light_scenario == "Gamvik") {
    comp_cost_current <- comp_cost_scenario + 0.11079 + 6.323e-03*time_current - 1.990e-
04*time_current^2 + 8.119e-06*time_current^3 - 1.916e-07*time_current^4 + 1.395e-
09*time_current^5

  }

  return(comp_cost_current)
}

competition_cost_baseline <- competition_cost_baseline(1:nb_timesteps)

competition_cost_baseline<-
ifelse(competition_cost_baseline[1:nb_timesteps]<0.0001,0.0001,competition_cost_baseline[
1:nb_timesteps])
```

```
competition_cost_baseline<-
ifelse(competition_cost_baseline[1:nb_timesteps]>0.2,0.2,competition_cost_baseline[1:nb_ti
mesteps])


get_competition_cost<- function(time_current){
  competition_cost_current <- competition_cost_baseline[time_current]
  return(competition_cost_current)
}





#Metabolism, transform the hourly cost into a per timestep cost
standardize_metabo_cost <- function(cost_hourly){
  fraction <- nb_hours/nb_timesteps
  cost <- cost_hourly*fraction
  return(cost)
}


#Calculate metabolism per hour, torpor (patch 1)
#Energy expenditure for torpor, per bat with 0 extra fat per hour
get_cost_torpor_hourly <- function(temperature_roost_current){
  cost_torpor_hourly <-
(0.00006*exp(0.1284*temperature_roost_current))*(mass_zero_fat+1) # From the PBZ paper
with brown long-eared bats, adding 1 gram of stomach content / fat reserves as most bats
usually have some energy reserves
  if(cost_torpor_hourly<bmr*0.9){
    cost_torpor_hourly
  }
  else{
    cost_torpor_hourly <- bmr*0.9
  }
  return(cost_torpor_hourly)
}
get_cost_torpor_hourly <- Vectorize(get_cost_torpor_hourly, vectorize.args =
"temperature_roost_current")


#Calculate metabolism per hour, resting (patch 2)
#Energy expenditure for resting, per bat with 0 extra fat per hour
get_cost_resting_hourly <- function(temperature_roost_current){
  if(temperature_roost_current>tnz){
    cost_resting_hourly <- bmr
  }
  else{
```

```
    cost_resting_hourly <- (0.0357-(0.0011*temperature_roost_current))*(mass_zero_fat+1) # From Geiser & Brigham 2000, adding 1 gram of stomach content / fat reserves as most bats usually have some energy reserves
  }
  return(cost_resting_hourly)
}
get_cost_resting_hourly <- Vectorize(get_cost_resting_hourly, vectorize.args = "temperature_roost_current")




# Cost of rewarming from different temperatures
get_cost_arousing <- function(temperature_roost_current){
  cost_arousing <- 0.1158621 - 0.003586207 * temperature_roost_current # Turbill's equation (converted to mass loss), total expenditure for active arousals
  if (cost_arousing > 0.005) {
    cost_arousing <- cost_arousing
  } else {
    cost_arousing <- 0.005
  }
  return(cost_arousing)
}
get_cost_arousing <- Vectorize(get_cost_arousing, vectorize.args = "temperature_roost_current")

library(magrittr)



patch1 <- rep(0, times = nb_timesteps)
patch2 <- rep(0, times = nb_timesteps)
patch3 <- seq(1, nb_timesteps, by=1)
patch3_very_warm <- standardize_metabo_cost((get_prey(get_temperature_ext_very_warm(patch3)))*(energy_gain))
patch3_dynamic_warm <- standardize_metabo_cost((get_prey(get_temperature_ext_dynamic_warm(patch3)))*(energy_gain))
patch3_stable_warm <- standardize_metabo_cost((get_prey(get_temperature_ext_stable_warm(patch3)))*(energy_gain))
patch3_dynamic_cold <- standardize_metabo_cost((get_prey(get_temperature_ext_dynamic_cold(patch3)))*(energy_gain))
patch3_stable_cold <- standardize_metabo_cost((get_prey(get_temperature_ext_stable_cold(patch3)))*(energy_gain))
```

```
patch3_very_cold <-
standardize_metabo_cost((get_prey(get_temperature_ext_very_cold(patch3)))*(energy_gain)
)

foraging_benefit = array(c(patch1,patch2,patch3_very_warm,
                    patch1,patch2,patch3_dynamic_warm,
                    patch1,patch2,patch3_stable_warm,
                    patch1,patch2,patch3_dynamic_cold,
                    patch1,patch2,patch3_stable_cold,
                    patch1,patch2,patch3_very_cold),dim =c(nb_timesteps,3,length(weather)))
rm(patch1)
rm(patch2)
rm(patch3_very_warm)
rm(patch3_dynamic_warm)
rm(patch3_stable_warm)
rm(patch3_dynamic_cold)
rm(patch3_stable_cold)
rm(patch3_very_cold)




#make dataframe that stores timestep and corresponding predation risk (independent of
weather)
patch1 <- rep(0, times = nb_timesteps)
patch2 <- rep(0, times = nb_timesteps)
patch3 <- seq(1, nb_timesteps, by=1)
patch3 <- get_predation(patch3)
risk_predation_all <- data.frame(patch1, patch2, patch3)
rm(patch1)
rm(patch2)
rm(patch3)


#make dataframe that stores timestep and corresponding energetic competition cost
(independent of weather)
patch1 <- rep(0, times = nb_timesteps)
patch2 <- rep(0, times = nb_timesteps)
#patch3 <- rep(0, times = nb_timesteps)
patch3 <- seq(1, nb_timesteps, by=1)
patch3 <- get_competition_cost(patch3)
competition_cost_all <- data.frame(patch1, patch2, patch3)
rm(patch1)
rm(patch2)
rm(patch3)


patch1_very_warm <-
standardize_metabo_cost(get_cost_torpor_hourly(get_temperature_roost_very_warm(1:nb_ti
mesteps)))
```

```
patch2_very_warm <-
standardize_metabo_cost(get_cost_resting_hourly(get_temperature_roost_very_warm(1:nb_t
imesteps)))
patch1_dynamic_warm <-
standardize_metabo_cost(get_cost_torpor_hourly(get_temperature_roost_dynamic_warm(1:n
b_timesteps)))
patch2_dynamic_warm <-
standardize_metabo_cost(get_cost_resting_hourly(get_temperature_roost_dynamic_warm(1:
nb_timesteps)))
patch1_stable_warm <-
standardize_metabo_cost(get_cost_torpor_hourly(get_temperature_roost_stable_warm(1:nb_
timesteps)))
patch2_stable_warm <-
standardize_metabo_cost(get_cost_resting_hourly(get_temperature_roost_stable_warm(1:nb
_timesteps)))
patch1_dynamic_cold <-
standardize_metabo_cost(get_cost_torpor_hourly(get_temperature_roost_dynamic_cold(1:nb
_timesteps)))
patch2_dynamic_cold <-
standardize_metabo_cost(get_cost_resting_hourly(get_temperature_roost_dynamic_cold(1:n
b_timesteps)))
patch1_stable_cold <-
standardize_metabo_cost(get_cost_torpor_hourly(get_temperature_roost_stable_cold(1:nb_ti
mesteps)))
patch2_stable_cold <-
standardize_metabo_cost(get_cost_resting_hourly(get_temperature_roost_stable_cold(1:nb_t
imesteps)))
patch1_very_cold <-
standardize_metabo_cost(get_cost_torpor_hourly(get_temperature_roost_very_cold(1:nb_tim
esteps)))
patch2_very_cold <-
standardize_metabo_cost(get_cost_resting_hourly(get_temperature_roost_very_cold(1:nb_ti
mesteps)))
patch3 <- rep(standardize_metabo_cost(cost_flight_hourly), times = nb_timesteps)
metabolic_cost_all <- array(c(patch1_very_warm,patch2_very_warm,patch3,
                     patch1_dynamic_warm,patch2_dynamic_warm,patch3,
                     patch1_stable_warm,patch2_stable_warm,patch3,
                     patch1_dynamic_cold,patch2_dynamic_cold,patch3,
                     patch1_stable_cold,patch2_stable_cold,patch3,
                     patch1_very_cold,patch2_very_cold,patch3),dim
=c(nb_timesteps,3,length(weather)))
rm(patch1_very_warm)
rm(patch2_very_warm)
rm(patch1_dynamic_warm)
rm(patch2_dynamic_warm)
rm(patch1_stable_warm)
rm(patch2_stable_warm)
rm(patch1_dynamic_cold)
rm(patch2_dynamic_cold)
rm(patch1_stable_cold)
```

```
rm(patch2_stable_cold)
rm(patch1_very_cold)
rm(patch2_very_cold)
rm(patch3)


#make dataframe for metabolic cost of arousing at any timestep on good vs bad days
arousal_cost_very_warm <-
get_cost_arousing(get_temperature_roost_very_warm(1:nb_timesteps))
arousal_cost_dynamic_warm <-
get_cost_arousing(get_temperature_roost_dynamic_warm(1:nb_timesteps))
arousal_cost_stable_warm <-
get_cost_arousing(get_temperature_roost_stable_warm(1:nb_timesteps))
arousal_cost_dynamic_cold <-
get_cost_arousing(get_temperature_roost_dynamic_cold(1:nb_timesteps))
arousal_cost_stable_cold <-
get_cost_arousing(get_temperature_roost_stable_cold(1:nb_timesteps))
arousal_cost_very_cold <-
get_cost_arousing(get_temperature_roost_very_cold(1:nb_timesteps))
arousal_cost_all <- array(c(arousal_cost_very_warm,
                            arousal_cost_dynamic_warm,
                            arousal_cost_stable_warm,
                            arousal_cost_dynamic_cold,
                            arousal_cost_stable_cold,
                            arousal_cost_very_cold),dim =c(nb_timesteps,1,length(weather)))
rm(arousal_cost_very_warm)
rm(arousal_cost_dynamic_warm)
rm(arousal_cost_stable_warm)
rm(arousal_cost_dynamic_cold)
rm(arousal_cost_stable_cold)
rm(arousal_cost_very_cold)



#---- Empty arrays (do not change) ----

# We make an empty array to hold the fitness values
fitness <- array(data = NA, dim = c(length(fat_discretized), nb_timesteps+1, nb_days,
length(weather), length(temp_state)),
            dimnames = list(fat_discretized, 1:(nb_timesteps+1), 1:(nb_days),
1:length(weather), 1:length(temp_state)) )

# And for the decision loop (patch choice)
patch_choice <- array(data = NA, dim = c(length(fat_discretized), nb_timesteps, nb_days,
length(weather),length(temp_state)),
                 dimnames = list(fat_discretized, 1:nb_timesteps, 1:nb_days, 1:length(weather),
1:length(temp_state)))


#---- Functions (do not change) ----
```

```
#### Calculating terminal fitness at the end of each day####
# Terminal fitness is the probability of surviving the last day (i.e. day_current+1)

calculate_survival_proba <- function(fat_state) {
  if (fat_state < fat_survival_threshold) {
    survival_proba <- 0
  } else {
    if (fat_state < fat_survival_decrease) {
      survival_proba <- ((fat_state-fat_survival_threshold)/(fat_survival_decrease -
fat_survival_threshold))
    } else {
      survival_proba <- 1
    }
  }
  return(survival_proba)
}

# Calculate daily terminal fitness F(fat_state, timestep_current+1, day_current), for every
day, where fat_state=state
for (j in 1:length(fat_discretized)) {
  fat_state <- fat_discretized[j]
  fitness[j, 1:(nb_timesteps+1), 1:(nb_days), 1:length(weather), 1:length(temp_state)] <-
calculate_survival_proba(fat_state)
}
rm(j)



#---- Interpolating the discrete state variable ----
# NOTE this function actually returns a fitness value!
# The computer discretizes the state variable  but in reality energetic reserves is a continuous
variable.We overcome this using interpolation (see C&M 2.1)


interpolate <- function (fat_state, timestep_current, day_current, weather_current,
temp_state_current) {
  # This function returns a fitness value, based on the fat state.

  # Function that returns the index of the closest discrete fat_state value. Only the first value
is returned if multiple equidistant values.
  closest_discrete_x <- function(fat_state) {
    return(which(abs(fat_discretized - fat_state) == min(abs(fat_discretized - fat_state)))[1])
  }

  # Function that interpolates between a and b. dx (delta_x) is either 0 and 1, depending
whether it is closer to a or b (see below).
  linear_interpolation <- function (a, b, dx) {
    return((1-dx)*a+ b*dx)
  }
```

```
  # First, if reserves are negative, return 0 (the bat is dead).
  if (fat_state < 0) {
    return(0)
  }

  # Otherwise, find and store the closest discretized fat value to our actual fat value.
  closest <- closest_discrete_x(fat_state)

  #Then, check if this closest value is larger or smaller to our actual value
  if (fat_state < fat_discretized[closest]) {
    j1 <- closest -1 #this will be used to calculate delta_x
    j2 <- closest    #this will be used to calculate delta_x
  } else if (fat_state > fat_discretized[closest]) {
    j1 <- closest    #this will be used to calculate delta_x
    j2 <- closest +1 #this will be used to calculate delta_x
  }
  # If neither are true, the fitness value for fat_state is already present in the matrix. No need
to interpolate.
  else {
    return( fitness[closest, timestep_current, day_current, weather_current,
temp_state_current])
  }

  # Calculate how fat_state is positioned in relation to fat_discretized[j1] and
fat_discretized[j2].
  # 0: closer to fat_discretized[j1]; 1: closer to fat_discretized[j2]
  delta_x <- (fat_state-fat_discretized[j1])/(fat_discretized[j2]-fat_discretized[j1])

  # Interpolate.
  return(linear_interpolation(fitness[j1, timestep_current, day_current, weather_current,
temp_state_current], fitness[j2, timestep_current, day_current, weather_current,
temp_state_current], delta_x))
}
```

```
#### SCRIPT WITH BACKWARDS ITERATION ####

source("1. Prep. of parameters and functions.R")

# ---- Backwards Iteration ----

# Iterate backwards across days at nb_timesteps+1 (daily terminal fitness)
for (weather_current in 1:length(weather)) { # for all the weather values (1-6)
  for (day_current in nb_days:1) { # and then for all the days
    if (day_current > 1) {
      if (day_current != nb_days) { #if this is not the last day (for which fitness has already
been calculated)
        for (fat_current in 1:length(fat_discretized)) { # then for each fat level
          #message(paste0('Calculating end of day fitness for weather type ', weather_current, ',
on day ', day_current, ' with fat level #', round(fat_current, digits = 2), ' for both states...'))
          for (temp_state_current in 1:length(temp_state)) { # and then for each state
            fat_state <- fat_discretized[fat_current] #store the fat level in fat_state (this makes the
code more readable)
          } #end state loop
        } #end fat loop
      }


      # Iterate backwards from max nb_timesteps for each day
      for (timestep_current in nb_timesteps:1) { # for this day


        #message(paste0('Calculating for all states and decisions for weather type ',
weather_current, ', on day ', day_current, ' at timestep ', timestep_current, '...'))
        for (fat_current in 1:length(fat_discretized)) { # and for all fat states

          for (temp_state_current in 1:length(temp_state)) { # and for all behaviours

            fat_state <- fat_discretized[fat_current] #store the fat state to make the code more
readable
            fitness_all_choices <- vector(mode = 'numeric', length=3) #make a vector to
temporarily store fitness values for all three patches

            if (fat_state < fat_survival_threshold) {
              # Bat is dead. It will remain dead.
              fat_state <- 0
              fitness_all_choices <- 0
            } else {

              # Calculate resulting fitness of choosing each patch
              for (patch_chosen in 1:3) { # for all patches (patch 1 = torpor, patch 2 = rest, patch 3
= forage)
                if (temp_state_current == 1) { # if bat is in state 1 (torpid)

                  if (patch_chosen == 1) { #if this is patch 1 (torpid)
```

```
        if (timestep_current == nb_timesteps) { # If t=72, then the interpolation needs to replace t+1 (t=73) with t=1 from next day, accounting for the probability of day types (using a weighted average)

          fitness_if_staying_torpid <- numeric()

          for (weather_tomorrow in 1:length(weather)) {

            fat_when_torpid <- (fat_state - metabolic_cost_all[timestep_current,patch_chosen,weather_current]) #calculate torpid fat state
            fitness_if_staying_torpid <- c(fitness_if_staying_torpid,interpolate(fat_when_torpid, timestep_current+1, day_current, weather_tomorrow,1)) #calculate corresponding fitness (no predation, so just interpolating the fat state)

          }

          fitness_all_choices[patch_chosen] <- weighted.mean(fitness_if_staying_torpid,weather_prob) #store it


        } else { # If timestep is not 72
          fat_when_torpid <- (fat_state - metabolic_cost_all[timestep_current,patch_chosen,weather_current]) #calculate torpid fat state
          fitness_if_staying_torpid <- interpolate(fat_when_torpid, timestep_current+1, day_current, weather_current,1) #calculate corresponding fitness (no predation, so just interpolating the fat state)
          fitness_all_choices[patch_chosen] <- fitness_if_staying_torpid #store it

        }


      } else { #if not choosing to stay in torpid patch (i.e. if bat chooses to go to resting or foraging patch)
        if (timestep_current == nb_timesteps) { # If t=72, then the interpolation needs to replace t+1 (t=73) with t=1 from next day, accounting for the probability of day types (using a weighted average)

          fitness_both_food_and_no_food_patch_2 <- numeric()
          fitness_both_food_and_no_food_patch_3 <- numeric()

          for (weather_tomorrow in 1:length(weather)) {

            fat_if_food <- min(fat_max,(fat_state + foraging_benefit[timestep_current,patch_chosen,weather_current] - metabolic_cost_all[timestep_current,patch_chosen,weather_current] - arousal_cost_all[timestep_current,1,weather_current] -
```

```
competition_cost_all[timestep_current,patch_chosen]))# Ensure that fat_expected does not exceed fat_max.
            fat_if_no_food <- (fat_state - metabolic_cost_all[timestep_current,patch_chosen,weather_current] - arousal_cost_all[timestep_current,1,weather_current] - competition_cost_all[timestep_current,patch_chosen])

            fitness_if_food <- interpolate(fat_if_food, timestep_current+1, day_current, weather_tomorrow,2) - (risk_predation_all[timestep_current,patch_chosen]*exp(predation_risk_increase*fat_state) ) # må ha med torpor-state
            fitness_if_no_food <- interpolate(fat_if_no_food, timestep_current+1, day_current, weather_tomorrow,2) - (risk_predation_all[timestep_current,patch_chosen]*exp(predation_risk_increase*fat_state) )

            fitness_both_food_and_no_food_patch_2 <- c(fitness_both_food_and_no_food_patch_2,(fitness_if_food*(prob_finding_food*exp(foraging_success_decline*fat_state)) + fitness_if_no_food*(1-(prob_finding_food*exp(foraging_success_decline*fat_state))) + resting_fitness_benefit))
            fitness_both_food_and_no_food_patch_3 <- c(fitness_both_food_and_no_food_patch_3,(fitness_if_food*(prob_finding_food*exp(foraging_success_decline*fat_state)) + fitness_if_no_food*(1-(prob_finding_food*exp(foraging_success_decline*fat_state)))))
          }
          if (patch_chosen == 2) { #If the bats choose patch 2, there is an added fitness benefit
            #store mean fitness
            fitness_all_choices[patch_chosen] <- weighted.mean(fitness_both_food_and_no_food_patch_2,weather_prob) #store it
          } else if (patch_chosen == 3) {
            fitness_all_choices[patch_chosen] <- weighted.mean(fitness_both_food_and_no_food_patch_3,weather_prob) #store it
          }



        } else {  # If timestep is not 72

          fat_if_food <- min(fat_max,(fat_state + foraging_benefit[timestep_current,patch_chosen,weather_current] - metabolic_cost_all[timestep_current,patch_chosen,weather_current] - arousal_cost_all[timestep_current,1,weather_current] - competition_cost_all[timestep_current,patch_chosen]))# Ensure that fat_expected does not exceed fat_max.
          fat_if_no_food <- (fat_state - metabolic_cost_all[timestep_current,patch_chosen,weather_current] - arousal_cost_all[timestep_current,1,weather_current] - competition_cost_all[timestep_current,patch_chosen])

          #calculate corresponding fitness
```

```
                fitness_if_food <- interpolate(fat_if_food, timestep_current+1, day_current, weather_current,2) - (risk_predation_all[timestep_current,patch_chosen]*exp(predation_risk_increase*fat_state) ) # må ha med torpor-state
                fitness_if_no_food <- interpolate(fat_if_no_food, timestep_current+1, day_current, weather_current,2) - (risk_predation_all[timestep_current,patch_chosen]*exp(predation_risk_increase*fat_state) )


                if (patch_chosen == 2) { #If the bats choose patch 2, there is an added fitness benefit
                  #store mean fitness
                  fitness_all_choices[patch_chosen] <- fitness_if_food*(prob_finding_food*exp(foraging_success_decline*fat_state)) + fitness_if_no_food*(1-(prob_finding_food*exp(foraging_success_decline*fat_state))) + resting_fitness_benefit
                } else if (patch_chosen == 3) {
                  fitness_all_choices[patch_chosen] <- fitness_if_food*(prob_finding_food*exp(foraging_success_decline*fat_state)) + fitness_if_no_food*(1-(prob_finding_food*exp(foraging_success_decline*fat_state)))
                }
              }
            }


          } else if (temp_state_current == 2) { # if bat is in state 2 (awake)
            if (patch_chosen == 1) { #if this is patch 1 (torpid)

              if (timestep_current == nb_timesteps) { # If t=72, then the interpolation needs to replace t+1 (t=73) with t=1 from next day, accounting for the probability of day types (using a weighted average)

                fitness_if_going_torpid <- numeric()

                for (weather_tomorrow in 1:length(weather)) {

                  fat_if_going_torpid <- (fat_state - metabolic_cost_all[timestep_current,patch_chosen,weather_current])
                  fitness_if_going_torpid <- c(fitness_if_going_torpid,interpolate(fat_if_going_torpid, timestep_current+1, day_current, weather_tomorrow,1))

                }

                fitness_all_choices[patch_chosen] <- weighted.mean(fitness_if_going_torpid,weather_prob) #store it


              } else {  # If timestep is not 72
```

```
                    fat_if_going_torpid <- (fat_state -
metabolic_cost_all[timestep_current,patch_chosen,weather_current]) #calculate torpid fat
state
                    fitness_if_going_torpid <- interpolate(fat_if_going_torpid, timestep_current+1,
day_current, weather_current,1) #calculate corresponding fitness (no predation, so just
interpolating the fat state)
                    fitness_all_choices[patch_chosen] <- fitness_if_going_torpid #store it
                  }

                } else { # if patch chosen is 2 or 3, calculate fat if it finds food, and fat if it does
not

                  if (timestep_current == nb_timesteps) { # If t=72, then the interpolation needs to
replace t+1 (t=73) with t=1 from next day, accounting for the probability of day types (using
a weighted average)

                    fitness_both_food_and_no_food_patch_2 <- numeric()
                    fitness_both_food_and_no_food_patch_3 <- numeric()

                    for (weather_tomorrow in 1:length(weather)) {

                      fat_if_food <- min(fat_max,(fat_state +
foraging_benefit[timestep_current,patch_chosen,weather_current] -
metabolic_cost_all[timestep_current,patch_chosen,weather_current] -
competition_cost_all[timestep_current,patch_chosen]))# Ensure that fat_expected does not
exceed fat_max.
                      fat_if_no_food <- (fat_state -
metabolic_cost_all[timestep_current,patch_chosen,weather_current] -
competition_cost_all[timestep_current,patch_chosen])

                      fitness_if_food <- interpolate(fat_if_food, timestep_current+1, day_current,
weather_tomorrow,2) -
(risk_predation_all[timestep_current,patch_chosen]*exp(predation_risk_increase*fat_state) )
# må ha med torpor-state
                      fitness_if_no_food <- interpolate(fat_if_no_food, timestep_current+1,
day_current, weather_tomorrow,2) -
(risk_predation_all[timestep_current,patch_chosen]*exp(predation_risk_increase*fat_state) )

                      fitness_both_food_and_no_food_patch_2 <-
c(fitness_both_food_and_no_food_patch_2,(fitness_if_food*(prob_finding_food*exp(foragi
ng_success_decline*fat_state)) + fitness_if_no_food*(1-
(prob_finding_food*exp(foraging_success_decline*fat_state))) + resting_fitness_benefit))
                      fitness_both_food_and_no_food_patch_3 <-
c(fitness_both_food_and_no_food_patch_3,(fitness_if_food*(prob_finding_food*exp(foragi
ng_success_decline*fat_state)) + fitness_if_no_food*(1-
(prob_finding_food*exp(foraging_success_decline*fat_state)))))
                    }
                    if (patch_chosen == 2) { #If the bats choose patch 2, there is an added fitness
benefit
                      #store mean fitness
```

```
            fitness_all_choices[patch_chosen] <-
weighted.mean(fitness_both_food_and_no_food_patch_2,weather_prob) #store it
          } else if (patch_chosen == 3) {
            fitness_all_choices[patch_chosen] <-
weighted.mean(fitness_both_food_and_no_food_patch_3,weather_prob) #store it
          }



        } else {  # If timestep is not 72

          fat_if_food <- min(fat_max,(fat_state +
foraging_benefit[timestep_current,patch_chosen,weather_current] -
metabolic_cost_all[timestep_current,patch_chosen,weather_current] -
competition_cost_all[timestep_current,patch_chosen]))# Ensure that fat_expected does not
exceed fat_max.
          fat_if_no_food <- (fat_state -
metabolic_cost_all[timestep_current,patch_chosen,weather_current] -
competition_cost_all[timestep_current,patch_chosen])

          #calculate corresponding fitness
          fitness_if_food <- interpolate(fat_if_food, timestep_current+1, day_current,
weather_current,2) -
(risk_predation_all[timestep_current,patch_chosen]*exp(predation_risk_increase*fat_state) )
# må ha med torpor-state
          fitness_if_no_food <- interpolate(fat_if_no_food, timestep_current+1,
day_current, weather_current,2) -
(risk_predation_all[timestep_current,patch_chosen]*exp(predation_risk_increase*fat_state) )


          if (patch_chosen == 2) { #If the bats choose patch 2, there is an added fitness
benefit
            #store mean fitness
            fitness_all_choices[patch_chosen] <-
fitness_if_food*(prob_finding_food*exp(foraging_success_decline*fat_state)) +
fitness_if_no_food*(1-(prob_finding_food*exp(foraging_success_decline*fat_state))) +
resting_fitness_benefit
          } else if (patch_chosen == 3) {
            fitness_all_choices[patch_chosen] <-
fitness_if_food*(prob_finding_food*exp(foraging_success_decline*fat_state)) +
fitness_if_no_food*(1-(prob_finding_food*exp(foraging_success_decline*fat_state)))
          }
        }
      }

    } # end ifelse temp_state_current

  }  # end for patch_chosen loop

 }  # end ifelse fat_state
```

```
        # find and store highest fitness patch for this timestep/day/weather/state
        fitness[fat_current, timestep_current, day_current, weather_current,
temp_state_current] <- max(fitness_all_choices)[1]

        # choose lowest risk patch if multiple patches have the same fitness
        patch_choice[fat_current, timestep_current, day_current, weather_current,
temp_state_current] <- which(fitness_all_choices == max(fitness_all_choices))[1]

        patch_choice_stable_very_warm_state1 <- patch_choice[,,20,1,1] # optimal decision
matrix for a bat in torpid state on a very warm day
        patch_choice_stable_dynamic_warm_state1 <- patch_choice[,,20,2,1] # optimal
decision matrix for a bat in torpid state on a dynamic warm day
        patch_choice_stable_stable_warm_state1 <- patch_choice[,,20,3,1] # optimal decision
matrix for a bat in torpid state on a stable warm day
        patch_choice_stable_dynamic_cold_state1 <- patch_choice[,,20,4,1] # optimal
decision matrix for a bat in torpid state on a dynamic cold day
        patch_choice_stable_stable_cold_state1 <- patch_choice[,,20,5,1] # optimal decision
matrix for a bat in torpid state on a stable cold day
        patch_choice_stable_very_cold_state1 <- patch_choice[,,20,6,1] # optimal decision
matrix for a bat in torpid state on a very cold day

        patch_choice_stable_very_warm_state2 <- patch_choice[,,20,1,2] # optimal decision
matrix for a bat in awake state on a very warm day
        patch_choice_stable_dynamic_warm_state2 <- patch_choice[,,20,2,2] # optimal
decision matrix for a bat in awake state on a dynamic warm day
        patch_choice_stable_stable_warm_state2 <- patch_choice[,,20,3,2] # optimal decision
matrix for a bat in awake state on a stable warm day
        patch_choice_stable_dynamic_cold_state2 <- patch_choice[,,20,4,2] # optimal
decision matrix for a bat in awake state on a dynamic cold day
        patch_choice_stable_stable_cold_state2 <- patch_choice[,,20,5,2] # optimal decision
matrix for a bat in awake state on a stable cold day
        patch_choice_stable_very_cold_state2 <- patch_choice[,,20,6,2] # optimal decision
matrix for a bat in awake state on a very cold day


      } # end j loop
     } # end temp_state_current loop
    } # end timestep_current loop
    fitness[fat_current, nb_timesteps+1, day_current-1, weather_current, temp_state_current]
<- fitness[fat_current, timestep_current, day_current, weather_current, temp_state_current]
   } # end if day > 1
  } # end day_current loop
} #end good/bad weather loop

rm(patch_chosen)
rm(fat_current)

#END OF BACKWARDS ITERATION#
```

```
#### SCRIPT WITH FORWARD ITERATION ####

source("1. Prep. of parameters and functions.R")
source("2. Backwards iteration.R")


###### FORWARD ITERATION ######

# The below section implements forward iteration of the model (i.e. looking at the fate of
individuals in the model).

# Number of days to forward iterate for.
nb_days_forward  <- 30

# nb_timesteps/day is given from the model above.

# Number of individuals to iterate for each fat_state_init_forward Total number of
individuals
# is length(fat_state_init_forward)*nb_indiv_forward:
nb_indiv_forward <- 200

# Initial states for individual (A vector of indices in fat_discretized).
fat_state_init_forward <- 20 #11:30

temp_state_init_forward <- 2 #bats start in the awake state
#first_day <- 1 # For making figure 3 and 4

# state_forward_all[fat_state_init_forward, N, D, T]:
# State of individual N with initial state fat_state_init_forward, at day D and time T.
fat_state_forward_all <- array(data = NA, dim = c(length(fat_state_init_forward),
nb_indiv_forward, nb_days_forward*nb_timesteps +1), dimnames = list(
  fat_discretized[fat_state_init_forward],
  1:nb_indiv_forward,
  1:(nb_days_forward*nb_timesteps+1)
))

temp_state_forward_all <- array(data = NA, dim = c(length(fat_state_init_forward),
nb_indiv_forward, nb_days_forward*nb_timesteps +1), dimnames = list(
  rep(temp_state_init_forward, times = length(fat_state_init_forward)),
  1:nb_indiv_forward,
  1:(nb_days_forward*nb_timesteps+1)
))

decision_all <- array(data = NA, dim = c(length(fat_state_init_forward), nb_indiv_forward,
nb_days_forward*nb_timesteps +1), dimnames = list(
  fat_discretized[fat_state_init_forward],
  1:nb_indiv_forward,
  1:(nb_days_forward*nb_timesteps+1)
))
```

```
day_type_all <- array(data = NA, dim = c(nb_indiv_forward, nb_days_forward*nb_timesteps
+1), dimnames = list(
  1:nb_indiv_forward,
  1:(nb_days_forward*nb_timesteps+1)
))

# Keeping the sequence of day types days equal for all individuals
# weather_sequence <- sample(weather, size=nb_days_forward, replace=TRUE, prob =
weather_prob)

for (j in 1:length(fat_state_init_forward)) {
  fat_state_init_forward_discretized <- fat_discretized[fat_state_init_forward[j]]
  for (n in 1:nb_indiv_forward) {
    # Set the initial state.
    fat_state_forward_all[j, n, 1] <- fat_state_init_forward_discretized
    temp_state_forward_all[j, n, 1] <- temp_state_init_forward

    # All bats experience their own day type sequence. To keep this similar across individuals
move it out of the loop (see above)
    weather_sequence <- sample(weather, size=nb_days_forward, replace=TRUE, prob =
weather_prob)
    # weather_sequence <- c(first_day, sample(weather, size=nb_days_forward-1,
replace=TRUE, prob = weather_prob)) # For making figure 3 and 4

    # Iterate through the time.
    for (z in 1:(nb_days_forward*nb_timesteps) ) {
      timestep_current = (z-1) %%  nb_timesteps  +1 # nb_timesteps of day
      day_current = (z-1) %/% nb_timesteps  +1 # Day
      # The current state.
      fat_state <- fat_state_forward_all[j, n, z]
      temp_state_current <- temp_state_forward_all[j, n, z]

      if (fat_state < fat_survival_threshold) {
        # Bat is dead. It will remain dead.
        fat_state_new_forward <- -1
        temp_new <- 0
        h <- 0

      } else {
        # The best decision given the current state (Found by rounding fat_state to the closest
item
        # in fat_discretized).
        # TODO: interpolate the decision between the two closest optimal decisions.

        if (weather_sequence[day_current] == 1) {
          if (temp_state_current == 1) {
            h <- patch_choice_stable_very_warm_state1[which(abs(fat_discretized - fat_state) ==
min(abs(fat_discretized - fat_state)))[1], timestep_current]
            metabolism <- metabolic_cost_all[timestep_current,h,1]
            foraging   <- 0
```

```
          day_type <- 1

        } else if (temp_state_current == 2) {
          h <- patch_choice_stable_very_warm_state2[which(abs(fat_discretized - fat_state) ==
min(abs(fat_discretized - fat_state)))[1], timestep_current]
          metabolism <- metabolic_cost_all[timestep_current,h,1]
          foraging   <- foraging_benefit[timestep_current,h,1]
          day_type <- 1

        }

      } else if (weather_sequence[day_current] == 2) {
        if (temp_state_current == 1) {
          h <- patch_choice_stable_dynamic_warm_state1[which(abs(fat_discretized -
fat_state) == min(abs(fat_discretized - fat_state)))[1], timestep_current]
          metabolism <- metabolic_cost_all[timestep_current,h,2]
          foraging   <- 0
          day_type <- 2

        } else if (temp_state_current == 2) {
          h <- patch_choice_stable_dynamic_warm_state2[which(abs(fat_discretized -
fat_state) == min(abs(fat_discretized - fat_state)))[1], timestep_current]
          metabolism <- metabolic_cost_all[timestep_current,h,2]
          foraging   <- foraging_benefit[timestep_current,h,2]
          day_type <- 2
        }

      } else if (weather_sequence[day_current] == 3) {
        if (temp_state_current == 1) {
          h <- patch_choice_stable_stable_warm_state1[which(abs(fat_discretized - fat_state)
== min(abs(fat_discretized - fat_state)))[1], timestep_current]
          metabolism <- metabolic_cost_all[timestep_current,h,3]
          foraging   <- 0
          day_type <- 3

        } else if (temp_state_current == 2) {
          h <- patch_choice_stable_stable_warm_state2[which(abs(fat_discretized - fat_state)
== min(abs(fat_discretized - fat_state)))[1], timestep_current]
          metabolism <- metabolic_cost_all[timestep_current,h,3]
          foraging   <- foraging_benefit[timestep_current,h,3]
          day_type <- 3
        }

      } else if (weather_sequence[day_current] == 4) {
        if (temp_state_current == 1) {
          h <- patch_choice_stable_dynamic_cold_state1[which(abs(fat_discretized - fat_state)
== min(abs(fat_discretized - fat_state)))[1], timestep_current]
          metabolism <- metabolic_cost_all[timestep_current,h,4]
          foraging   <- 0
          day_type <- 4
```

```
        } else if (temp_state_current == 2) {
          h <- patch_choice_stable_dynamic_cold_state2[which(abs(fat_discretized - fat_state)
== min(abs(fat_discretized - fat_state)))[1], timestep_current]
          metabolism <- metabolic_cost_all[timestep_current,h,4]
          foraging   <- foraging_benefit[timestep_current,h,4]
          day_type <- 4
        }

      } else if (weather_sequence[day_current] == 5) {
        if (temp_state_current == 1) {
          h <- patch_choice_stable_stable_cold_state1[which(abs(fat_discretized - fat_state) ==
min(abs(fat_discretized - fat_state)))[1], timestep_current]
          metabolism <- metabolic_cost_all[timestep_current,h,5]
          foraging   <- 0
          day_type <- 5

        } else if (temp_state_current == 2) {
          h <- patch_choice_stable_stable_cold_state2[which(abs(fat_discretized - fat_state) ==
min(abs(fat_discretized - fat_state)))[1], timestep_current]
          metabolism <- metabolic_cost_all[timestep_current,h,5]
          foraging   <- foraging_benefit[timestep_current,h,5]
          day_type <- 5
        }

      } else if (weather_sequence[day_current] == 6) {
        if (temp_state_current == 1) {
          h <- patch_choice_stable_very_cold_state1[which(abs(fat_discretized - fat_state) ==
min(abs(fat_discretized - fat_state)))[1], timestep_current]
          metabolism <- metabolic_cost_all[timestep_current,h,6]
          foraging   <- 0
          day_type <- 6

        } else if (temp_state_current == 2) {
          h <- patch_choice_stable_very_cold_state2[which(abs(fat_discretized - fat_state) ==
min(abs(fat_discretized - fat_state)))[1], timestep_current]
          metabolism <- metabolic_cost_all[timestep_current,h,6]
          foraging   <- foraging_benefit[timestep_current,h,6]
          day_type <- 6
        }
      }

      if (runif(1) <=
risk_predation_all[timestep_current,h]*exp(predation_risk_increase*fat_state)) { # Predation
risk.

        # Negative fat_state = dead.
        fat_state_new_forward <- -1
        temp_new <- 0
```

```
        } else {

          if (runif(1) <= prob_finding_food) {
            success <- 1
          } else {
            success <- 0
          }

          fat_state_new_forward <- fat_state + foraging*success - metabolism

          if (h == 1) {
            temp_new <- 1
          } else {
            temp_new <- 2
          }
        }
      }

      # Set new state.
      fat_state_forward_all[j, n, z+1] <- fat_state_new_forward
      decision_all[j, n, z+1] <- h
      temp_state_forward_all[j, n, z+1] <- temp_new
      day_type_all[n, z] <- day_type
    }
  }
}

rm(h)
rm(j)
rm(z)


#END OF FORWARD ITERATION#
```
